# The transcriptional hallmarks of intra-tumor heterogeneity across a thousand tumors

**DOI:** 10.1101/2021.12.19.473368

**Authors:** Avishai Gavish, Michael Tyler, Dor Simkin, Daniel Kovarsky, L. Nicolas Gonzalez Castro, Debdatta Halder, Rony Chanoch-Myers, Julie Laffy, Michael Mints, Alissa R. Greenwald, Adi Wider, Rotem Tal, Avishay Spitzer, Toshiro Hara, Amit Tirosh, Sidharth V. Puram, Mario L. Suva, Itay Tirosh

## Abstract

Each tumor contains malignant cells that differ in genotype, phenotype, and in their interactions with the tumor micro-environment (TME). This results in distinct integrated cellular states that govern intra-tumor heterogeneity (ITH), a central challenge of cancer therapeutics. Dozens of recent studies have begun to describe ITH by single cell RNA-seq, but each study typically profiledonly a small number of tumors and provided a narrow view of transcriptional ITH. Here, we curate, annotate and integrate the data from 77 different studies to reveal the patterns of ITH across 1,163 tumor samples covering 24 tumor types. Focusing on the malignant cells, we find thousands of transcriptional ITH programs that can be described by 41 consensus meta-programs (MPs), each consisting of dozens of genes that are coordinately upregulated in subpopulations of cells within many different tumors. The MPs cover diverse cellular processes and differ in their cancer-type distribution. General MPs associated with processes such as cell cycle and stress vary within most tumors, while context-specific MPs reflect the unique biology of particular cancer types, often resembling developmental cell types and suggesting the co-existence of variable differentiation states within tumors. Some of the MPs are further associated with overall tumor proliferation or immune state, highlighting their potential clinical significance. Based on functional similarities among MPs, we propose a set of 11 hallmarks that together account for the majority of observed ITH programs. Given the breadth and scope of the investigated cohort, the MPs and hallmarks described here reflect the first comprehensive pan-cancer description of transcriptional ITH.

## Introduction

Intra-tumor heterogeneity (ITH) is a fundamental property of tumors that is driven by genetics, epigenetics and micro-environmental influences, and is central to treatment failure, metastasis and other cancer phenotypes (Marusyk et al., 2012; McGranahan and Swanton, 2015). Single cell RNA-sequencing (scRNA-seq) efficiently enables the characterization of ITH, and has seen a rapid expansion of its use across virtually all major cancer types (Suva and Tirosh, 2019). One emerging concept from recent studies applying scRNA-seq to tumor samples is the existence of ITH “expression programs”, consisting of sets of dozens of genes with coordinated variability in their expression across cells within a given tumor. In melanoma, a skin pigmentation-related program driven by *MITF* and an EMT-like program associated with *AXL*, varied within multiple individual tumors and had important functional consequences (Rambow et al., 2018; Shaffer et al., 2017; Tirosh et al., 2016a). In glioblastoma, four expression programs were identified as a central source of transcriptional heterogeneity (Neftel et al., 2019). In head and neck squamous cell carcinoma (HNSCC), EMT-like and Epithelial Senescence (EpiSen) programs were identified and shown to affect the likelihood for metastasis and drug responses (Kinker et al., 2020; Puram et al., 2017). A stress-response program was found in multiple cancer types with important functional implications (Baron et al., 2020).

The examples of ITH expression programs described above share three important features: First, they often account for a large fraction of the expression variability among malignant cells within a tumor, e.g. reflecting the first or second principal components or accounting for the main clusters of cells in analysis of expression heterogeneity. Second, the program gene-sets are enriched with specific functional annotations, thus delineating their likely biological function. Third, while the programs are first defined *within* each individual tumor, there are highly significant similarities between the ITH programs that are identified across tumors of the same cancer type, and in some cases even across distinct cancer types (Baron et al., 2020; Kinker et al., 2020), in spite of their distinct genetic background. These similarities suggest that the ITH expression programs may reflect essential aspects of the basic biology of tumors. To capture this basic biology, we seek to identify the shared genes among ITH programs identified in different tumors, reasoning that these are more likely to reflect the underlying biological process and are less dependent on patient-specific genetic and epigenetic features. We therefore define sets of genes that are highly shared among multiple ITH programs with overall similarities, and we refer to such consensus gene-sets of ITH programs as “meta-programs” (or MPs).

High expression of any particular MP may be considered as defining a *cellular state*. However, it is important to note that MPs tend to be limited to dozens of genes whose expression is superimposed on the cells’ baseline expression profile and therefore reflect a *relative* cellular state. For example, two subpopulations of cells from two distinct tumors may upregulate the same MP (e.g. cell cycle) while retaining the extensive expression differences between these two tumors (e.g. due to unique driver mutations). These two subpopulations would be in a different *global* cellular state (reflecting the tumor identity and genetics) but in the same *relative* cellular state (reflecting the activation of a particular MP). In this work we focus primarily on relative cellular states by defining the expression programs (and meta-programs) of each subpopulation of cells relative to the other cells from the same tumor, hence highlighting the patterns of *intra*-tumor heterogeneity as opposed to *inter*-tumor variability.

The functional and clinical significance of MPs identified previously and the emerging notion that they may reflect a central feature of ITH raises the need to comprehensively define the MPs in cancer and further explore their biology and function. We previously profiled 198 cancer cell lines by scRNA-seq and uncovered 12 *in vitro* MPs (Kinker et al., 2020). Here, we aim to expand this analysis to tumor samples derived directly from patients. To that end, we integrate scRNA-seq data across 77 studies that profiled patient samples from 24 cancer types, define 41 MPs and investigate their functional enrichments, distribution across cancers and potential implications as hallmarks of ITH in cancer.

## Results

### Comprehensive curation of cancer scRNA-seq datasets

With the expanding use of scRNA-seq, many publications have profiled tumors from most major cancer types (Gonzalez Castro et al., 2021). While these studies provide various insights into cancer biology, they frequently did not define ITH programs or MPs. The few studies which did define ITH programs together represent only a minority of cancer types, and in most cases had insufficient cohorts to robustly delineate MPs. Moreover, these studies typically did not integrate their findings with those of other studies, yielding insights that were mostly specific to small cohorts from one cancer type or subtype. By combining data from all of these studies we set to systematically define cancer MPs and identify similarities and differences between cancer types (**Fig. 1A, Fig. S1A**).

**Figure 1.**
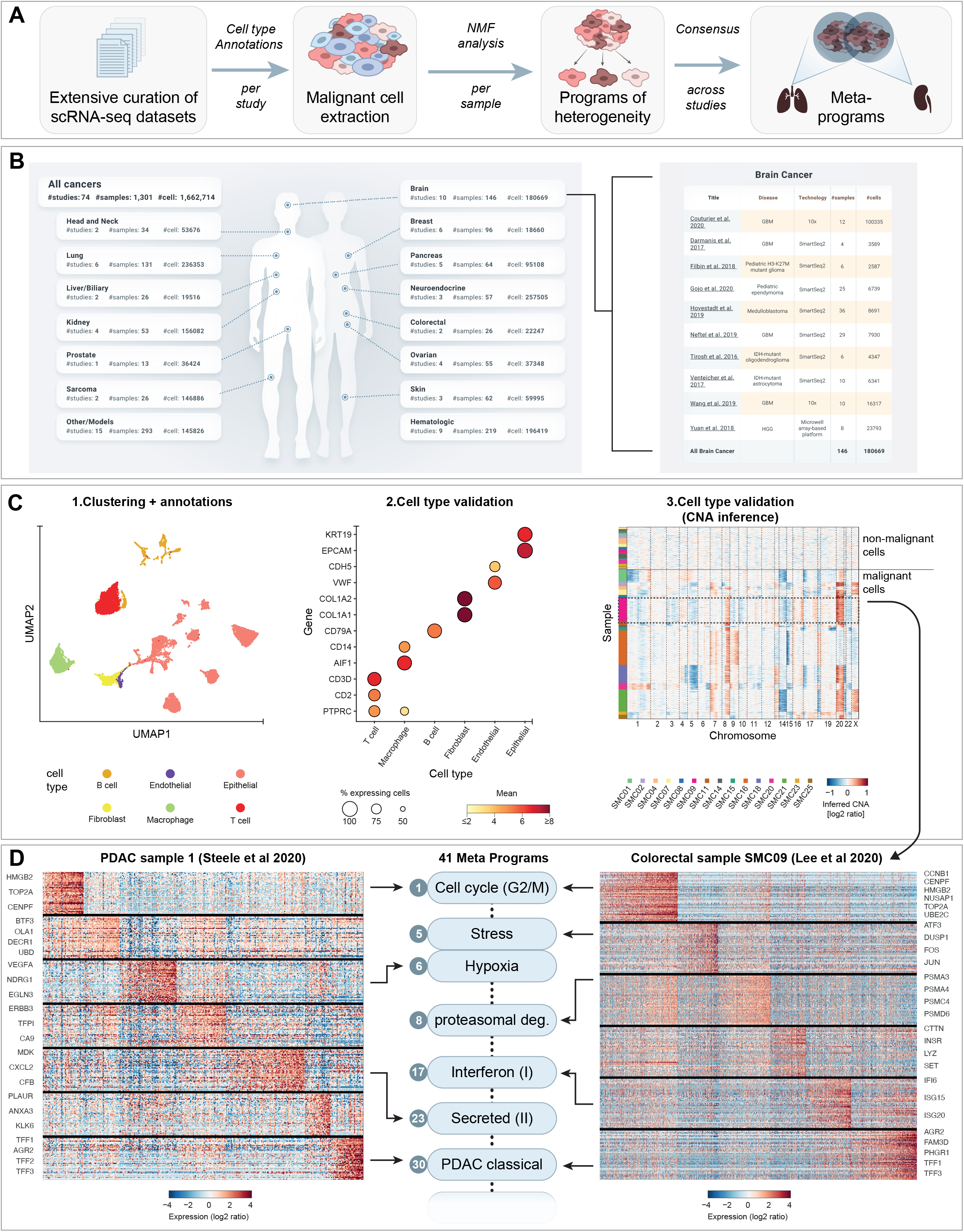
Defining transcriptional ITH from a curated and annotated collection of scRNA-seq datasets. (**A**) Workflow. (**B**) Left: the analyzed scRNA-seq datasets are summarized by the number of studies, number of profiled tumors and number of profiled cells, per cancer type (rows) and for all cancer types combined (top row). Right: summary of the brain cancer scRNA-seq datasets, with each row corresponding to one study, and listing the number of profiled tumors, number of profiled cells, type of brain cancer analyzed and scRNA-seq platform used. (**C**) Cell annotations of the Lee et al. 2020 dataset by three steps, exemplifying the approach for annotation of all datasets. 1: UMAP plot of all cells colored by their cell type assignment as determined by the original study. 2: Manual validation of the cell type assignments based on expression of canonical marker genes. 3: inference of CNAs from gene expression profiles separates malignant cells with CNAs (bottom) from non-malignant cells without CNAs (top). (**D**) Expression programs of heterogeneity as defined by NMF, for the SMC9 tumor from Lee et al. 2020 (right) and the ‘sample 1’ tumor from Steele et al. 2020 (left). In each case, a heatmap shows the relative expression, for the top 50 genes from each of the robust NMF programs (rows), across all malignant cells from the tumor (columns). Genes are arranged by NMF programs and selected genes are labeled. The middle section shows the inclusion of 7 of the 41 NMF programs in MPs and the associated MP names.

We systematically searched for all studies that reported scRNA-seq data of human tumors. We prioritized studies based on the amount of data for malignant cells (as opposed to non-malignant immune and/or stromal cells), the ability to obtain such data from public databases or upon request from the authors and the quality of the data (Methods). We further included several unpublished scRNA-seq datasets from ongoing studies by our group on neuroendocrine tumors, head and neck cancer and schwannoma. Finally, while we prioritized data for human tumors, we also incorporated a few selected datasets of either mice models or cell models. Altogether, we obtained data from 77 studies, encompassing 1456 samples covering 24 cancer types and profiling 2,591,545 cells (**Fig. 1B**, **Table S1**).

### Annotation and availability of a cancer scRNA-seq compendium

We used two complementary approaches to annotate cells from each dataset. First, we assigned cells to cell types (**Fig. 1C**, **step 1**). Cell type assignments from the original study were publically available for ~28% of the datasets, were directly obtained from the authors upon specific request for another ~67% of the datasets and were missing in the remaining ~5% of datasets, which we annotated by clustering and manual annotation based on differentially expressed genes. We then verified and refined these annotations based on expression of canonical cell type markers, and by excluding questionable clusters and apparent doublets (**Fig. 1C**, **step 2**). In some cases the original study’s annotations were partial (e.g. assigning some cells only as malignant or non-malignant), but for most studies we obtained a detailed assignment of the non-malignant cells into 7 common cell types and 31 other cell types (**Table S1**).

Second, for each study, we inferred copy-number alterations (CNAs) from the gene expression profiles, as described previously (Tirosh et al., 2016a), to assign cells as malignant based on tumor-specific CNAs (**Fig. 1C, step 3**). This distinction by CNAs is important given that a large proportion of tumors contain malignant and non-malignant cell types with globally similar expression profiles (e.g. 67% of carcinoma samples appear to contain non-malignant epithelial cells, such that an epithelial assignment is not sufficient to identify malignant cells). Cells with borderline CNA signals were excluded from further analysis as they likely reflect either doublets or cancer cells in which low-quality data is limiting the ability to detect CNAs and thus might also limit any further analysis of their cellular state. Based on this procedure we defined 686,690 high-quality malignant cells and 1,199,312 non-malignant cells that were further divided into specific cell types.

This large curated and annotated pan-cancer compendium is used here to define cancer MPs, but it also constitutes a resource of major utility for various future analyses. For most of the datasets included in the compendium the cell annotations were only available upon request from the authors and the available annotations were often lacking. These limitations highlight the difficulty for researchers to re-analyze scRNA-seq datasets and the importance of an easily accessible, consistently formatted, and comprehensively annotated compendium. Therefore, we provide all data through a dedicated website called the Curated Cancer Cell Atlas, or 3CA (https://weizmann.ac.il/sites/3CA). Apart from providing the original datasets and the refined cell annotations described above, the 3CA includes inferred CNAs, UMAP plots, associated statistics and other advanced analyses that are described below and in upcoming publications. To ensure the utility, comprehensiveness and accuracy of 3CA data, we will continue to curate and annotate cancer scRNA-seq datasets and will include on the 3CA website the updated results along with extended functionalities and graphical interfaces.

### Defining expression programs of malignant intra-tumor heterogeneity

Having built 3CA, we turned to these curated data to comprehensively define expression patterns of intra-tumor heterogeneity among malignant cells. The large number of studies and diverse experimental and computational methodologies used to profile malignant cells introduces a challenge for integrating these data and avoiding batch effects. Recent studies developed multiple computational methods for integration of scRNA-seq data (Barkas et al., 2019; Korsunsky et al., 2019; Stuart et al., 2019), but these are typically designed for integration of a small number of studies rather than dozens of independent studies. Moreover, even efficient integration methods cannot fully distinguish between technical and biological variability, such that batch removal strategies will also remove many genuine biological patterns. Notably, since our primary interest is in variability *within* individual tumors (rather than *between* tumors), direct integration of datasets is not essential for our analyses. Thus, we avoided the use of direct integration methods and instead first defined expression programs that vary *within* each individual tumor (with each program represented by a set of correlated genes) and subsequently compared the programs across studies. The rationale behind this approach is that while absolute expression profiles often suffer from prominent batch effects, the patterns of variability *within* tumors are defined only from comparisons within the same batch and hence are much less sensitive to batch effects and recover high similarities across diverse studies (**Fig. S1B**).

For each tumor, we utilized non-negative matrix factorization (NMF) to characterize the expression programs that vary among its malignant cells, each summarized by its top-scoring 50 genes (**Fig. 1D**). NMF was applied with multiple values of the rank parameter (K = 4..9) and the NMF programs that were consistently identified in a tumor by multiple rank values were denoted as *robust* NMF expression programs (Methods). Overall, we identified 5547 robust NMF programs that were retained for further analysis (**Table S2**).

### Recurrent meta-programs covering diverse biological functions

Since our goal was to identify recurrent patterns of ITH, we clustered the robust NMF programs by their fractions of shared top genes (quantified by a Jaccard index, see **Fig. S1A**), and filtered out programs that were associated with low quality or doublets. This resulted in 41 clusters of programs (**Fig. 2A-B, Table S2**). All clusters were derived from multiple studies and 83% of them were derived from multiple cancer types, highlighting their robustness and pan-cancer significance. The clusters covered 66% of all robust NMF programs, indicating that most patterns of ITH reflect recurrent patterns that may be described by meta-programs (MPs). Thus, for each cluster, a meta-program was defined as the set of genes most commonly shared between programs from that cluster (see Methods). 54% of all malignant cells were enriched for at least one of the MPs, indicating that MPs capture a high fraction of the ITH patterns. Similarly, when clustering the malignant cells in each sample by Louvain clustering (see methods), 59% of the clusters were enriched with at least one MP (**Fig.S2**).

**Figure 2.**
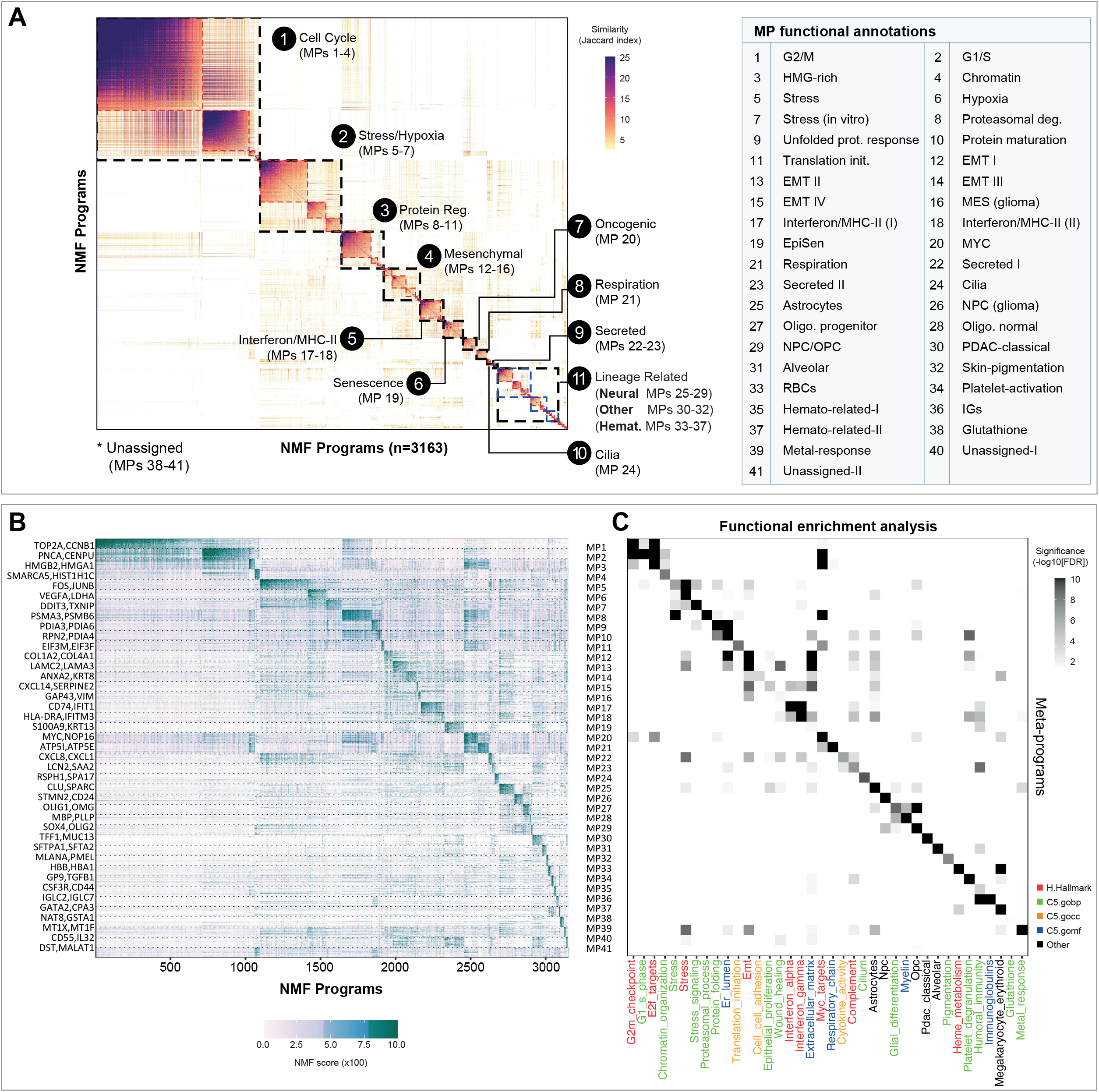
ITH meta-programs and their functional annotations. (**A**) *Left*: heatmap showing Jaccard similarity indices for comparisons among 3163 robust NMF programs based on their top 50 genes. Programs are ordered by clustering (see Methods) and grouped into MPs (marked by red dashed lines) and families of MPs with relate functions (marked by black dashed lines); MP families are numbered and labeled. *Right*: table listing all MP names. (**B**) Heatmap showing NMF scores for all MP genes (rows) across all robust NMF programs (columns, arranged as in A). Dashed lines separate the genes of different MPs; MP numbers (right) and selected genes (left) are indicated. (**C**) Significance (−log10(FDR)) of the enrichment of selected functional annotations (columns) in all of the MPs (rows, ordered as in B). Annotations are colored by their database origin, as indicated (right legend). See **Table S3** for additional functional annotations.

MPs were named based on their diverse functional enrichments (**Fig. 2C, Table S3**), and MPs with related functions were further grouped into 11 MP families (**Fig. 2A**). The 41 MPs are further described and visualized in additional detail in **Supplementary Note 1**, which includes functional enrichments, selected genes, and expression patterns in selected tumors.

The most abundant MP families in tumors were largely associated with expected and well-described patterns of ITH, including cell cycle (MP1-4), stress/hypoxia (MP5-7), and mesenchymal states (MP12-16). Nonetheless, specific MPs within these families reflected interesting variants of these known ITH patterns that were either not described previously or remain poorly understood. For example, among cell cycle MPs, apart from the canonical G2/M and G1/S MPs (MP1,2, respectively), two less frequent MPs consisted of cell cycle-related genes but were also specifically enriched with genes encoding for HMG-box proteins (MP3) or for chromatin regulators (MP4). Among mesenchymal MPs, we found multiple variants of EMT that differ in their abundance across cancer types, as well as a “hybrid” program that includes activation of both mesenchymal and epithelial markers (MP14), and a mesenchymal MP that was specific to glioma (MP15).

MP families of intermediate abundance across tumors included some that resembled previously described ITH patterns (protein regulation, interferon response, epithelial senescence, cilia) but also several MPs that to our knowledge were not described previously in tumor scRNA-seq datasets, including for example MYC-targets (MP20). MPs with lower abundance (<1% of NMF programs) were primarily not described previously to our knowledge, highlighting the increased sensitivity in detecting recurrent ITH programs with such a large and diverse compendium. Many of the low-abundance MPs were enriched with functional annotations linked to the biology of a specific tissue or lineage (e.g. brain or blood-related MPs) and therefore we grouped these together in a MP family denoted as “lineage-related”.

### Distribution of meta-programs across cancer types

To systematically define the distribution and context-specificity of MPs, we classified the abundance of each MP in each broadly-defined cancer type as either absent, low, medium, high, or high and significantly enriched (**Fig. 3A**; see Methods). We then examined the number of cancer types in which each MP is found at a medium or higher frequency (**Fig. 3B**). Seven MPs were each found in most of the cancer types, thereby defining the “general” MPs. These include MPs associated with the G1/S and G2/M cell cycle phases; stress, hypoxia and interferon responses; EMT-III; and MYC-targets.

**Figure 3.**
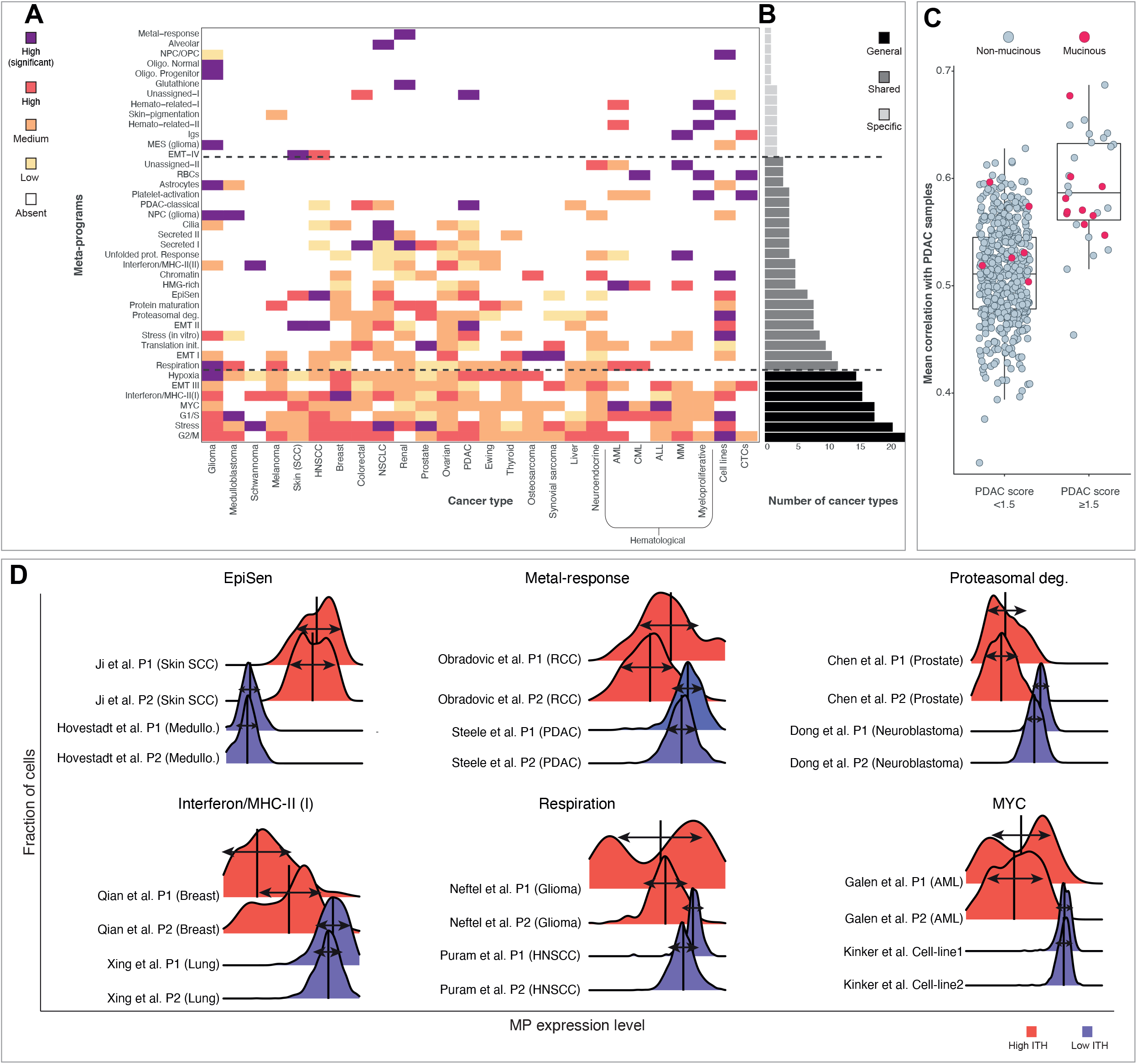
Variability and absolute expression of meta-programs across cancer types. **(A)** Abundance of each MP (rows) in each cancer type (columns), defined as either absent, low, medium, high or high and significant (see Methods for exact definitions). (**B**) Bar plot showing, for each MP, the number of cancer types with a medium or higher abundance. Rows correspond to the MPs as labeled in (A). MPs are further divided into three abundance categories by the dashed black lines. (**C**) Average correlation of each LUAD TCGA samples to all PDAC TCGA samples. LUAD samples are divided into those with or without high expression of MP30 (score>1.5) and are colored based on their histological classification as mucinous or as non-mucinous lung adenocarcinomas. (**D**) For each of six selected MPs (panels), the distribution of MP scores across malignant cells from a particular tumor is shown for four different tumors - two from a cancer type with low variability across cells (i.e. narrow distributions, shown in blue), and two from a cancer type with high variability across cells (i.e. wide distributions, shown in red). The plots demonstrate that for some MPs (e.g. EpiSen), cancer types with high average expression also tend to have high within-tumor variability, while for many other MPs (i.e. the five other examples), high within-tumor variability is not associated with high average expression.

On the other end of the spectrum, 13 MPs were found only in one or two cancer types, defining “context-specific” MPs. Note that while context-specific MPs tend to also be lineage-related MPs, these are distinct definitions based on the MPs occurrence across cancer types vs. their functional enrichments, respectively. The context-specific MPs were primarily specific to glioma, melanoma, hematologic malignancies, squamous cell carcinoma (head and neck and skin), and to renal or lung adenocarcinoma. The remaining MPs (21 out of 41), which we denote as “shared” were neither general nor context-specific, as they were detected in 3 to 12 cancer types.

For most of the *context*-*specific* MPs and for some of the shared MPs, the observed distribution was expected based on their known biology, such that *lineage*-*related* MPs were observed as variable in the cancer types derived from the corresponding lineages. However, several MPs had unexpected distributions, such as MP30, which is linked to pancreatic adenocarcinoma (PDAC) but is observed as variable more broadly, as described below. While the exact biological meaning of MP30 is unclear, this MP is remarkably consistent with the signature of the “classical” subtype of PDAC (Moffitt et al., 2015). Thus, this expression program distinguishes between the two main PDAC subtypes - classical and basal - but also varies within individual PDAC tumors, regardless of their subtype (Chan-Seng-Yue et al., 2020).

As expected, MP30 was primarily observed as variable within PDAC tumors, yet it was also observed as variable within several lung, colorectal, liver and head and neck cancers (**Fig. 1D**, **Fig. 3A**). This observation was most robust among lung adenocarcinomas (LUADs), which we investigated further. MP30 was observed as variable within 5 LUADs from 3 studies. These tumors also had high absolute expression of MP30 genes as well as of other PDAC-enriched genes (**Fig. S3**), suggesting that they reflect a subset of LUADs with certain features that resemble PDAC. To better understand this subset, we searched for such tumors among TCGA LUAD samples, as the TCGA cohort is large (n=508) and well annotated genetically and clinically. The LUAD tumors with highest scores for MP30 also had overall increased expression similarity to PDAC tumors (**Fig. 3C**) and an enrichment of KRAS mutations which are most common in PDAC (*p*=0.0032, hypergeometric test). These LUAD tumors with features of PDAC were highly enriched with histological classification as invasive mucinous adenocarcinoma (**Fig. 3C,** *P*=9.7e^−10^), suggesting a link between this histology and the PDAC features, consistent with previous results (Shim et al., 2015).

### Absolute expression vs. variability of meta-programs

While the PDAC-related MP30 is observed more broadly than expected, other MPs are conversely observed in a more narrow distribution than might be expected. For example, two MPs enriched with glutathione-related genes and with metal response genes, respectively, were found to be variable within tumors exclusively (but consistently) in renal cancer, although activity of these genes and associated processes are not specific to the kidney. The genes associated with both of these MPs were on average more highly expressed in other cancer types but their coherent variability was found only in renal cancer (**Fig. S4**).

This result highlights an important but often overlooked distinction between the specificity of an expression program’s activity and its variability: an expression program may be expressed in various contexts but may vary significantly only within certain contexts, which are not necessarily the ones with the highest absolute expression. A systematic comparison between activity and variability shows that for most MPs, variability is linked to the activity of the program but also highlights many exceptions, including the two renal-specific MPs noted above (**Fig. 3D, Fig. S4**).

Additional examples include: (i) Proteasomal degradation and translation initiation, which are most often variable in prostate cancer as well as in cell lines, despite lower than average activity in these contexts. (ii) Interferon, which is most often variable in breast cancer despite moderate activity. (iii) Hypoxia, which is more often variable in sarcoma and glioma than in multiple carcinoma types with higher absolute expression. (iv) Respiration, which is most often variable in brain cancers (glioma and medulloblastoma) despite low activity. (v) MYC-targets, which is most often variable in leukemia but is higher in many other contexts. Notably, MYC targets have both the highest activity and the least variability in cell lines, perhaps indicating that MYC activity is selected for during the establishment of cell lines such that its variability is avoided in culture. These examples may provide clues to specific mechanisms that facilitate (or hinder) the variability of such programs within specific tumors and decouple them from absolute activity. This decoupling limits the ability to infer ITH patterns from bulk profiles, which reflect only absolute (i.e. average) activity.

### Meta-program associations with proliferation

To further explore the functional significance of MPs we examined their association with two central cancer phenotypes, namely proliferation and immune state. Proliferation scores were defined as the maximal expression scores of the cell cycle MPs. In each tumor, we calculated the correlation between proliferation scores of cells and their expression of all other MPs detected in that tumor. We then averaged those correlations across all tumors to define the overall propensity of cells in each state to proliferate (**Fig. S5**). Most MPs are slightly negatively correlated with proliferation, perhaps indicating that cycling cells tend to repress other programs to divert their energy to proliferation. The most significant negative correlation (*P*=9.8e^−32^) was found for MP19, consistent with its annotation as epithelial senescence (EpiSen). EpiSen was previously described in HNSCC (Kinker et al., 2020; Puram et al., 2017), but in this work we find that it is an abundant program observed across 131 tumors from 9 tumor types.

On the other extreme, we find three programs that are positively associated with proliferation: MPs 21, 8 and 20, annotated as respiration, proteasomal degradation and MYC-targets, respectively (*P*=7.8e^−11^, 8.8e^−14^ and 1.8e^−23^). The MYC-target program had the strongest correlation with proliferation, consistent with the known oncogenic function of MYC. This result suggests that rather than a uniform MYC activation at an early stage of oncogenesis, MYC-targets are further upregulated in a subset of cells that are prone to proliferate and that may constitute an efficient therapeutic target. The association of respiration with proliferation may suggest that despite the tendency of many cancers to rely primarily on glycolysis (Vander Heiden et al., 2009), subsets of cells with increased respiration may have increased proliferation capacity.

### T-cell states and their association with malignant MPs

Next, to examine the potential relevance of cancer MPs to immunotherapies we turned to focus on their associations with T cell states. Although our curation strategy prioritized datasets with malignant cells, the resulting compendium contained 195,881 annotated T-cells from 34 studies, which represent 20 cancer types. Thus, we applied to these T cells the same analysis as described above for the malignant cells, and identified 16 T-cell meta-programs (MP_T_s; **Fig. 4A**). These include four highly abundant MP_T_s associated with the cell cycle and with cytotoxic, naïve, and regulatory T-cells, that together cover the majority (76%) of robust NMF programs. The additional 12 MP_T_s were much less abundant, although they were mostly shared across different cancer types (**Fig. 4B**), in contrast to the malignant MPs of low abundance.

**Figure 4.**
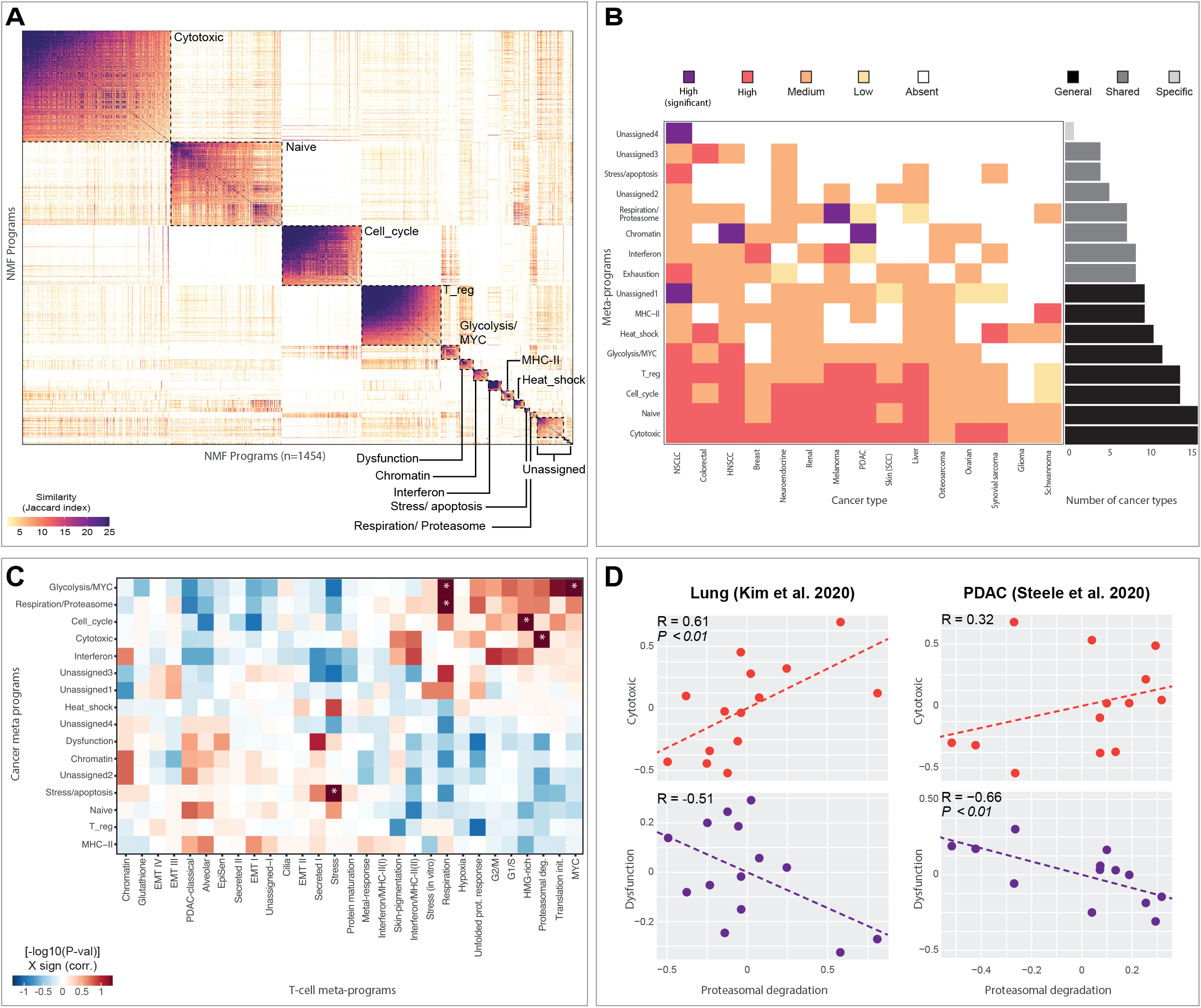
T-cell states and their association with malignant MPs. (**A**) Clustering of T-cell programs to define MP_T_s. Heatmap shows Jaccard similarity indices for comparisons among 1454 robust NMF T-cell programs based on their top 50 genes. Programs are ordered by clustering (see Methods) and grouped into MP_T_s (marked by black dashed lines) which are also labeled. (**B**) *Left*: abundance of each MP_T_ (rows) in each cancer type (columns), defined as either absent, low, medium, high or high and significant (see Methods for exact definitions). *Right*: Bar plot showing, for each MP_T_, the number of cancer types with a medium or higher abundance. MP_T_s are further divided into three abundance categories (General, Shared and Specific). (**C**) Heatmap showing the significance of the average correlations between malignant MPs (columns) and T-cell MP_T_s (rows), defined as −log10(P)*sign(R). R refers to the Pearson correlation between tumor scores for the corresponding MP and MP_T_; the correlation was calculated separately for each study and then averaged across studies. P refers to the p-value defined by t-test over the correlations from the individual studies. Asterisks denote significant values (P<0.05). (**D**) Scatterplots show scores for the proteasomal degradation MP (X-axis) vs. either the cytotoxic MP_T_ (Y-axis, top) or dysfunction MP_T_ (Y-axis, bottom). Dots correspond to tumors from two separate studies (left and right panels). Dashed lines show linear regression and the correlations are indicated for each plot along with indication of significant p-values.

Eight of the lower-abundant MP_T_s were annotated based on functional enrichments and marker genes (**Table S5**). MP_T_6 was annotated as “dysfunction” as it included several genes encoding for important co-inhibitory receptors that serve as immunotherapy targets (PD1, TIGIT, CD161, BTLA) (Lee et al., 2021; Mathewson et al., 2021), along with dysfunction/exhaustion-related transcription factors (TOX and TOX2) (Seo et al., 2019) and secreted factors (CXCL13) (Thommen et al., 2018). The other 7 annotated MP_T_s were not associated with T cell-specific states but rather with more generic cellular processes, resembling some of the malignant cell MPs. These include glycolysis/MYC-targets (MP_T_5), chromatin (MP_T_7), interferon response (MP_T_8), MHC-II (MP_T_9), heat shock (MP_T_10), stress/apoptosis (MP_T_11) and respiration/proteasome (MP_T_12). Despite the similarities between these MP_T_s and some of the malignant cell MPs, we note that the same functional enrichments are often represented by different genes in the two cell types, and that associations between functional enrichments were unique to each cell type (e.g. interferon response and MHC-II were coupled within malignant MPs, but associated with different T-cell MP_T_s).

Finally, we asked if malignant MPs and T-cell MP_T_s may be correlated. We defined tumor-level MP scores and MP_T_ scores (by averaging the scores of malignant cells and of T-cells within each tumor, respectively) and then examined their correlations across tumors. Due to the large differences between datasets of distinct studies, we calculated these correlation within each individual study, and then averaged the correlations across studies to identify consistent associations across diverse cancer types (**Fig. 4C**). We found six significant positive associations (by t-test across studies with FDR<0.05). Of these, four associations involved similar processes in the malignant cells and the T-cell: cell cycle, stress, respiration and MYC-targets. This may suggest that certain aspects of the TME may often induce similar processes in the malignant cells and the T cells.

One of the remaining two positive associations was between proteasomal degradation in the malignant cells (MP8) and cytotoxicity in the T-cells (MP_T_1) (**Fig. 4C-D**). Furthermore, we noted that MP8 was also negatively associated with T-cell dysfunction (MP_T_6), and while this association was not statistically significant when averaging all studies, it was significant in some of the individual studies (**Fig. 4C-D**). These results suggest an overall association between proteasomal degradation and immune activation, perhaps reflecting the role of proteasomal degradation in antigen-presentation through MHC-I, which could facilitate the recognition of cancer cells by T-cells and the subsequent activation of T-cells. As our pan-cancer analysis will expand into additional scRNA-seq tumor studies and to those with larger cohorts and more cell types, these analyses will be better powered to fully uncover such associations between cell types and to delineate their context-specificity.

## Discussion

While each tumor has a unique ITH pattern, ITH also involves variability of expression programs that are shared across tumors. Some aspects of ITH are therefore predictable and may warrant particular attention and potentially the development of novel therapeutic strategies. Here we aim to systematically uncover such shared patterns, by assembling an unprecedented number of cancer scRNA-seq datasets and performing integrative analysis. We uncovered 41 recurrent malignant programs that we term MPs and further characterized their functional enrichments, distribution across 24 cancer types and other associations.

Which patterns of ITH may have been overlooked in our analysis? First, rare patterns that either occur in a small proportion of tumors or in a small proportion of cells may not be identified in our current datasets. These may include patterns unique to cancer subtypes that were not sampled sufficiently in this cohort, or rare cells with abundance too low to be sampled robustly in the tumors that harbor them. Second, our unbiased approach defines variable expression programs based on the combined signal from thousands of genes, and hence is efficient in detecting large-scale programs with many co-expressed genes. Programs that involve only a handful of genes or that primarily involve proteins or metabolites, rather than changes at the mRNA level, are likely to be missed, and may require a supervised approach to be detected based on prior knowledge or particular genes of interest.

In future analyses, additional scRNA-seq datasets will undoubtedly enable the identification of novel MPs, as well as the refinement of existing MPs. However, we reason that this initial set of MPs is based on a sufficiently large and diverse tumor cohort to capture the canonical transcriptional ITH patterns, especially those that are commonly expressed across cancers. Accordingly, the MPs reported here may be regarded as a preliminary description of the hallmarks of ITH at the level of gene expression. Each MP (or family of related MPs), except for those of particularly low abundance, is considered here as one hallmark, thereby defining 11 hallmarks of transcriptional ITH (**Fig. 5A**). We named the hallmarks broadly (e.g. “oncogenic” rather than MYC-targets, and “lineage-related” rather than naming by specific lineages) to ensure their consistency with future data and additional MPs, but also included hallmarks that occur in a limited proportion of tumors. The two highly abundant hallmarks (cell cycle and stress) together cover most observed patterns of transcriptional ITH, followed by nine hallmarks with variable frequencies and context-specificity (**Fig.5A-B**).

**Figure 5.**
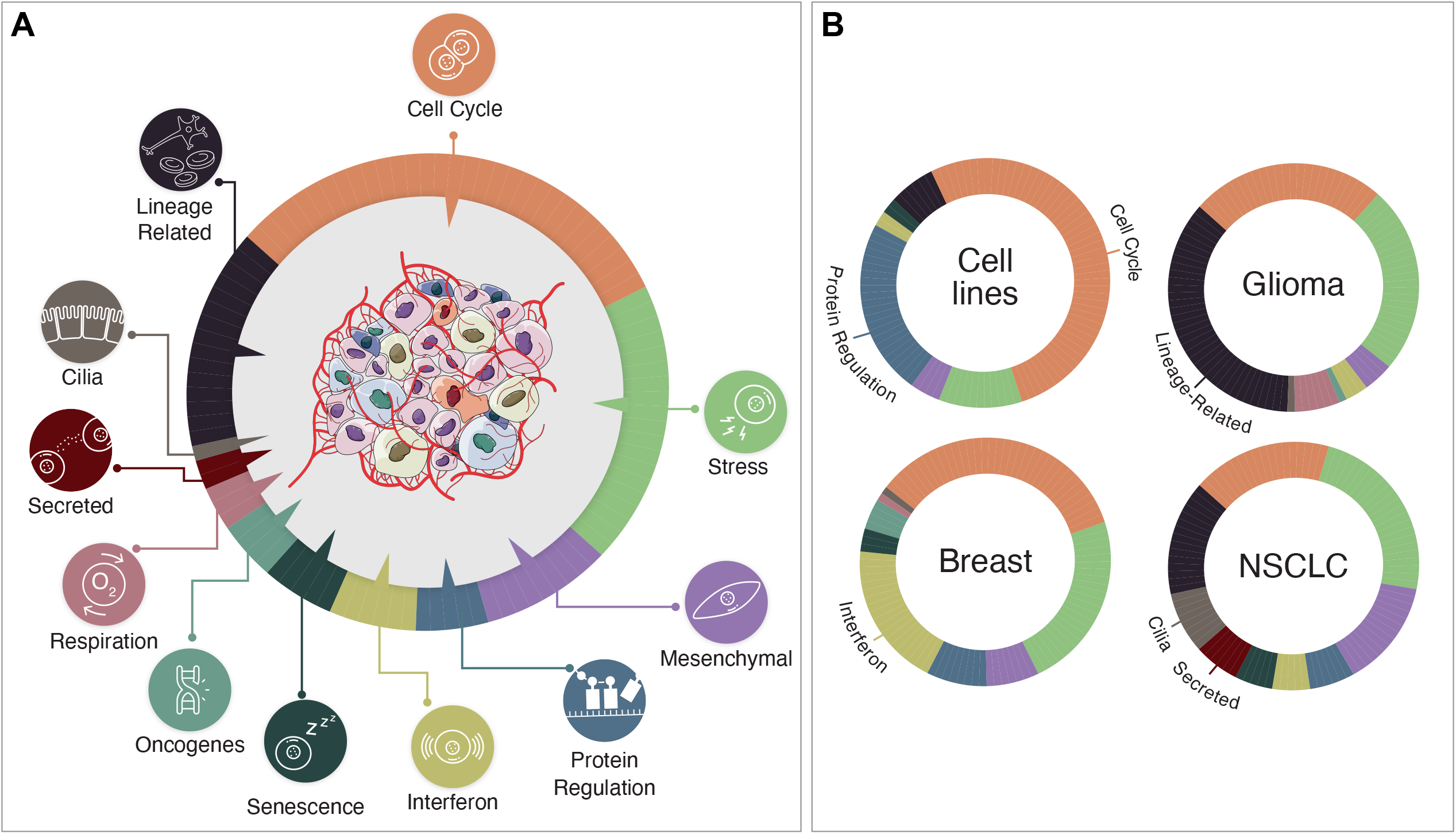
MP families reflect hallmarks of transcriptional ITH with varying frequencies across cancer types. (**A**) The circle reflects the 11 hallmarks of transcriptional ITH, with each corresponding to one family of MPs. The size of each section is proportional to the abundance of the MP family across all tumor samples (see **Table S4**). (**B**) Same as A, with each panel corresponding to hallmark frequencies in a particular type of samples – cell lines, glioma, breast cancers and lung cancers. Hallmarks with high frequency in each type of samples, compared to all other samples, are labeled.

Many of the hallmarks are largely recapitulated in cancer cell lines, highlighting the utility of cell lines for studying at least some common aspects of ITH (**Fig. 5B**). The hallmarks that are entirely absent in cell lines include hypoxia (likely reflecting the higher oxygen concentration of standard cell cultures), MYC-targets (invariably high in cell lines, possibly reflecting selection for MYC activity during establishment of cell lines), respiration, cilia, and secreted. In contrast, the protein regulation hallmark was particularly abundant in cell lines. Future studies may utilize the ITH patterns in mouse models, patient-derived xenografts, organoids and co-culture models to investigate the influence of particular non-malignant cell types on the variability of these ITH patterns that are absent in standard cell lines.

The lineage-related hallmark includes expression programs that are related to developmental cell types and typically linked to their differentiation (e.g. skin pigmentation, alveolar, astrocyte, neuronal progenitors, oligodendrocyte progenitor). Thus, variability of these programs within tumors implies variability in the degree of differentiation and supports the notion that ITH recapitulates developmental trajectories. These programs may also support the potential utility of differentiation therapies, perhaps implying that such therapies may be specifically relevant for cancers in which ITH is dominated by differentiation patterns.

Interestingly, abundance of lineage-related programs varied markedly between cancer types (**Fig.5B, Table S4**). Glioma had four different MPs belonging to this hallmark that together covered 36% of the robust NMFs, consistent with our prior work (Filbin et al., 2018; Neftel et al., 2019; Tirosh et al., 2016b). In contrast, lineage-related MPs were not detected at all in some cancer types. These include cancer types with limited scRNA-seq data, for which more data may be needed to detect such MPs (e.g. osteosarcoma), but also in common cancer types with extensive scRNA-seq data, such as ovarian and breast cancer. In those cancer types, lineage-related MPs might be relatively rare or too weak to be detected by the current analysis. Consistent with this possibility, when using a more lenient method for MP detection we identified variability of an additional Androgen Receptor (AR) program in a small fraction of breast cancers (**Fig. S6, Table S2**). Thus, while rare lineage-specific MPs may exist in any cancer type, the dominant patterns of ITH in most cancer types do not seem to directly reflect their specific lineage and development but rather points to variability of fundamental cellular processes, such as cell cycle, stress and anti-viral responses, protein regulation, and respiration, which all tend to be observed across most cancer types. These results highlight the broad similarities of ITH patterns across cancer types and are consistent with the emerging trend to consider pan-cancer treatments and tailor them to the molecular features of each tumor (e.g. the dominant MPs), regardless of the affected organ (Chen et al., 2021).

In summary, we curated and annotated a large scRNAseq cancer atlas that is publicly available for use by the research community and which enabled us to define the first comprehensive pan-cancer map of transcriptional ITH patterns. This map highlights 11 hallmarks of ITH, and while these will be refined and investigated further by future studies they are likely to endure due to th size and breadth of the present cohort. Hence, these hallmarks will guide our understanding of ITH and help to illuminate the path towards cancer therapies that consider not only tumor type and genetics but also internal cellular diversity.

## Acknowledgements

This work was supported by grants from the Israel Science Foundation (ISF), the Zuckerman STEM Leadership Program, the Mexican Friends New Generation, the Neuroendocrine Tumor Research Foundation, the Israel Cancer Research Fund (ISCR), the Benoziyo Endowment Fund, and Elías Harari. I.T. is the incumbent of the Dr. Celia Zwillenberg-Fridman and Dr. Lutz Zwillenberg Career Development Chair.

## Methods

### Data curation

A total of 77 Single cell RNA-seq (scRNA-seq) datasets, which constitute 1456 samples/tumors, were curated. Most of these datasets are available at https://www.weizmann.ac.il/sites/3CA aside from unpublished datasets and datasets for which we did not obtain sharing permission from the authors of the original studies. For data selection, we first constructed a list of potentially relevant studies through pubmed search, and continuously updated it through literature review. Each study was then examined for the type and amount of scRNA-seq data generated in it, prioritizing studies with data for multiple patient tumors and including a reasonable fraction of malignant cells. When seeking to add each new dataset to our cohort, we initially checked whether the data is publicly available for downloading (e.g. via the Gene Expression Omnibus database repository). Most publications freely provided the data in the form of an expression-matrix, together with a list of associated genes and cells (barcodes). Several datasets were only available upon author consent, in which case we contacted the authors for permission. The cohort also includes unpublished and published datasets that were previously analyzed in our lab, and that were already sequenced, aligned and processed. Most of the external datasets we downloaded, even when freely available, did not include the cell annotations presented in the published manuscript. In these cases, we contacted the leading authors and requested that they provide us with the annotations they used. In some cases, annotations of the original study were either not provided by the authors or were limited (e.g. not distinguishing between malignant and non-malignant epithelial cells), in which case we inferred the annotations ourselves.

### Verification of cell annotation

For each dataset – both for author-based annotations and for the annotations that we defined – we performed the following analyses to ensure the validity of the annotations. First, we performed dimensionality reduction using UMAP and examined whether the cells annotated as different cell types clustered separately. Second, we validated the annotations by verifying that the top differentially expressed genes of each cell type match known marker genes. Finally, we inferred Copy Number Aberrations (CNAs) (using the package available at https://github.com/jlaffy/infercna), to verify the annotation of cells as malignant. Some samples in which we could not resolve the annotations, possibly due to low data quality, were excluded during this process and were not used for further analysis.

### Data pre-processing

The following preprocessing steps were performed before conducting downstream analysis:

i. *Cell filtering:* We excluded cells with a low number of detected genes (#genes). For 10x data our cutoff was typically #genes > 1000. For smart-seq2 data we typically used a higher cutoff of #genes > 2000 genes. For other single cell platforms we adapted the cutoff and in some cases used a threshold lower than 1000 genes, but never lower than the cutoff used by the original study.
ii. *Sample filtering:* After cell filtering, we excluded samples having fewer than 10 malignant cells. Where relevant, we also excluded samples with unresolved CNA patterns. In total, 1163 samples were retained for downstream analysis.
iii. *Gene filtering:* Given an expression matrix *A* with *n* genes (rows) and *m* cells (columns), the mean expression of gene *i* across cells is given by 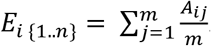. For most analysis we kept the 7000 genes with the highest *E*_*i*_ value in each sample.
iv. *Normalization:* UMI counts were converted to CPM (counts per million). For most analyses each entry in the matrix was then normalized according to 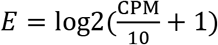. The same normalization was used for TPM values. The values were divided by 10 since the actual complexity is assumed to be in the realm of ~100,000 and not 1 million as implied by the CPM and TPM measures. (v) For most analyses the data was centered (each gene was centered across all cells). Centering was done separately for each study.

### Defining a non-redundant set of robust NMF programs

We performed non-negative matrix factorization (NMF) for each sample separately, to generate programs that capture the heterogeneity with each sample. Negative values in each centered expression matrix were set to zero. Since application of NMF requires a “K” parameter that influences the results, we ran NMF using different values (K=4,5,6,7,8,9), thereby generating 39 programs for each tumor. Each NMF program was summarized by the top 50 genes based on NMF coefficients. We reasoned that the most meaningful NMF programs are those that would recur across different values of K as well as across tumors; such programs (denoted here as *robust* NMF programs) were defined by the following three criteria:

i. *Robust within the tumor:* A program that is represented by multiple *similar* NMF programs, as defined for the same tumor when analyzed by multiple K value. Two NMF programs were considered as similar if they had at least 70% gene overlap (35 out of 50 genes).
ii. *Robust across tumors:* NMF programs that had at least 20% similarity (by top 50 genes) with any NMF program in any of the other tumors analyzed.
iii. *Non-redundant within the tumor:* within each tumor, NMF programs were ranked by their similarity (gene overlap) with NMFs from other tumors and selected in decreasing order. Once an NMF was selected, any other NMF within the tumor that had 20% overlap (or more) with the selected NMF was removed, to avoid redundancy. This approach yielded 5547 *robust* NMF programs.

### Defining Meta-programs (MPs)

We next clustered the *robust* NMF programs according to Jaccard similarity. The clustering was performed using a custom approach (see Figure S1A), that defined clusters of NMF programs and a list of 50 genes that constitute the MP. Briefly, each robust NMF was compared to all other robust NMFs to assess the degree of gene overlap between programs. Considering overlap instances of at least 10 genes, the NMF with the maximal number of considerable overlaps was selected as a potential founder of a new cluster. If the number of overlapping NMF programs (>10 genes) exceeded 5 cases, the NMF program with highest gene overlap to the founder NMF program was added, and thus a cluster was formed. The MP for the cluster was initially defined by the genes that appeared in both NMFs. To complete the list to 50 genes, the genes with the top NMF scores (in either NMF) were selected. The process was then repeated by searching for the NMF program with maximal overlap with the MP, adding it to the cluster while the overlap was at least 10 genes, and updating the MP (selecting genes that appeared in the highest number of NMF programs, and completing the MP list to 50 genes according to NMF scores in the . This way the MP was updated after addition of each NMF, reflecting the genes common to the NMFs constituting the cluster. The cluster was completed when no NMF could be added, and an attempt to form a new cluster was made as described above.

This approach yielded 67 initial MPs. We further removed MPs that were: (i) suspected to reflect low quality data and had a strong enrichment of either ribosomal protein genes or mitochondrial-encoded genes; or (ii) included NMF programs from only a single study, even though that study reflected a cancer type that is also examined by additional studies; or (iii) suspected to reflect doublet cells based on high similarity to the expression profile of a non-malignant cell type (e.g. T-cells or macrophages). We retained 41 MPs, and assessed their enrichment in functionally annotated gene-sets.

### Gene-sets for functional enrichment analyses

We primarily used signatures from MsigDB, including the following collections of gene-sets: Gene Ontology (C5.GOBP,C5.GOCC,C5.GOMF), Hallmark (H), and Cell-Types (C8). We also added selected additional signatures (not taken from MsigDB) related to brain/glioma, alveolar and PDAC signatures. Signatures with an FDR-adjusted P<0.05 (hypergeometric test) were considered significantly enriched (**Table S3**). The 41 MPs were further grouped by functional similarities into 11 hallmarks (**Figure 2**).

### MP abundance assessment

We first calculated, for each MP, the *observed* number of MP-related NMF programs in each cancer type. The *expected* abundance was then defined by multiplying the number of MP-related NMF programs (i.e. the MP size) by the total number of NMF programs identified in the cancer type, across all MPs, and dividing that by the total number of robust NMF programs. Finally, for each combination of cancer type and MP we calculated 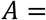 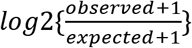 and the Bonferroni-adjusted p-value using a hypergeometric test. The abundance classification in Figure 3A was defined as follows:

- **Absent**: 0 MP-related NMF programs in that cancer type.
- **Low**: 1 MP-related NMF programs in that cancer type ***and*** −1.5 < *A* ≤ 0.
- **Medium**: Between 2 to 10 MP-related NMF programs in that cancer type ***or*** 0 < *A* ≤ 1.
- **High**: >10 MP-related NMF programs in that cancer type ***or*** *A* > 1,
- **High** (significant): Same as high ***and*** adjusted p-value < 0.05.

### Analysis of MP30 (PDAC-classical) in LUAD

Expression data for LUAD and PDAC tumors, in the form of RSEM ‘scaled estimates’, was obtained from TCGA via the Broad GDAC Firehose website (gdac.broadinstitute.org). These expression levels were converted to transcripts per million (TPM) and log-transformed, and patient samples were removed if they had no accompanying mutations data or if multiple tumor samples existed for the same patient. LUAD tumors were scored for the 50-gene PDAC-classical signature using a method described previously (Tirosh et al. 2016b), where signature gene expression is measured relative to a control gene set, which is chosen to have a similar distribution of expression levels but no coherent biological function. By manual inspection of the distribution of these scores, a long tail was observed above 1.5, hence this threshold was chosen to distinguish PDAC-classical-high LUAD tumors. Hypergeometric tests were used to quantify the enrichment of mucinous samples, and likewise the enrichment of KRAS mutations, among the PDAC-classical-high tumors. The mean correlation of each LUAD sample with PDAC samples was calculated as follows: first, for each of LUAD and PDAC, we computed the variability of all genes by median absolute deviation from the median and selected the 2500 genes with the highest such variability. Restricting to the intersection of these two gene sets (~1500 genes), we computed the pairwise correlations between LUAD and PDAC expression profiles. For each LUAD sample, we then computed the mean such correlation across PDAC samples (**Fig. 3C**). We also showed that LUAD samples that contributed an NMF program to the PDAC-classical MP had higher resemblance to PDAC samples by averaging the top 100 differentially expressed PDAC genes (obtained by comparing all PDAC samples to all LUAD samples) in LUAD samples (**Fig. S3**, *P*<0.05 by two-sample t-test).

### Global expression vs variability

To calculate the mean (global) expression of a MP and its variability within individual samples (Figure S4) we first averaged the (non-centered) expression of MP genes in each cell and log2-normalized each value. The mean of these values across cells represents the global MP expression in the sample, whereas the distribution’s width represents the degree of variability as shown in Figure 3D. To compare the global MP expression and variability within each cancer type, the global MP expression was averaged across samples from the same caner type. The variability of the MP expression in a given cancer type was determined as the fraction of tumors (out of the total number of tumors in that cancer type) that contributed at least one NMF program to the MP. For each MP we then calculated the Pearson correlation between global expression and variability across all cancer types.

### Estimating the fraction of NMF programs and Louvain clusters accounted for by MPs

Each individual cell was considered to score positively to a MP based on a one-sample t-test across the 50 genes of the MP, with null=0 since the input expression data has been centered per gene. We used a threshold of P<0.05 after adjusting for multiple comparisons. To determine whether a NMF scored significantly to a MP we first centered all the NMFs in each sample (centering each rank separately), and then scored against MPs as we did with cells. We then performed Louvain clustering for each sample using k=10, which resulted in an average of ~4 ± 1.8 clusters per tumor (33 tumors that had only 1 cluster were removed from this analysis). Clusters with more than 50% cells that scored positive to a MP were considered as being accounted for by an MP and their frequency in each sample was compared to the frequency of robust NMF programs in the sample that scored significantly positive a MP (**Figure S2**).

### MP association with proliferation

In each sample, we scored every cell to the 41 MPs and calculated the correlations between the scores (across cells) of the 4 cell cycle-related MPs and the rest of the MPs that were identified as variable within that sample. We kept the maximal correlation out of these 4, to represent correlation of a MP with cell cycle. We averaged the correlations of each MP across samples, retaining only the 29 MPs that were represented in at least 7 samples, and using one-sample t-test (with null=0) to define its significance (**Figure S5**).

### Defining T-cell Meta-programs (MPs)

A total of 195,881 cells from 34 studies were annotated as T-cells. We adopted an identical approach for defining T-cell MPs as described above for the cancer cells, which yielded 1,792 robust NMFs and 22 initial T-cell MPs. 16 MPs were finally selected after removing programs that we suspected represented low-quality cells or miss-annotations (e.g. B-cells or macrophages). Four of these MPs we could not assign to a functional annotations (**Fig. 4A**).

### Downstream analysis of T-cell (MP_T_s)

Abundance across cancer types (**Fig. 4B**) was defined by the same approach as described above for malignant cells (**Fig. 3A**). To test the significance of the correlations between cancer MPs and T-cell MP_T_s, we selected studies that contained at least 7 samples with both cancer cells and T-cells (a total of 19 studies). We next scored cancer cells for all cancer MPs, and similarly scored T-cells for all T-cell MPs (using the ‘sigScores’ function from https://github.com/jlaffy/scalop). Scores were averaged across cells to define tumor-level scores, and the correlations between tumor-level scores was calculated in each study for all cancer MPs vs. all T-cell MP_T_s. For each MP-MP_T_ combination, the mean correlation across studies was calculated, and its significance was evaluated by FDR-adjusted one-sample two-tailed t-test (**Fig. 4C**).

### Data availability

All curated data is available at (https://www.weizmann.ac.il/sites/3CA), aside from samples from unpublished studies that will be added when possible.

### Code availability

Any relevant code is available upon reasonable request from the corresponding author. Future versions of the website will also include the code used for downstream analysis.

## Notes

https://www.weizmann.ac.il/sites/3CA/

### Competing Interest Statement

The authors have declared no competing interest.

