## Supplemental Figures 1-6 for "The transcriptional hallmarks of intra-tumor heterogeneity across a thousand tumors"

Figure S1

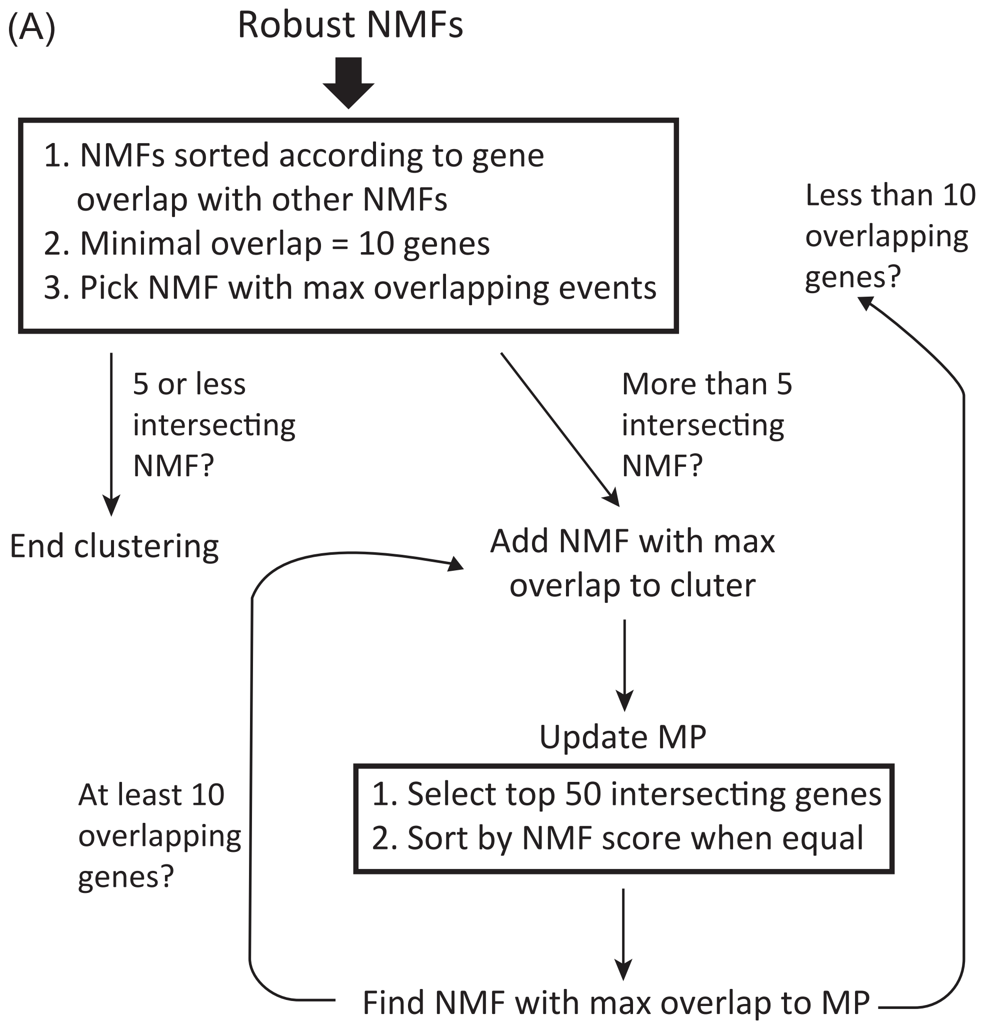

(A)
Robust NMFs
1. NMFs sorted according to gene overlap with other NMFs
2. Minimal overlap = 10 genes
3. Pick NMF with max overlapping events
5 or less intersecting NMF?
More than 5 intersecting NMF?
Less than 10 overlapping genes?
End clustering
Add NMF with max overlap to cluter
Update MP
1. Select top 50 intersecting genes
2. Sort by NMF score when equal
At least 10 overlapping genes?
Find NMF with max overlap to MP


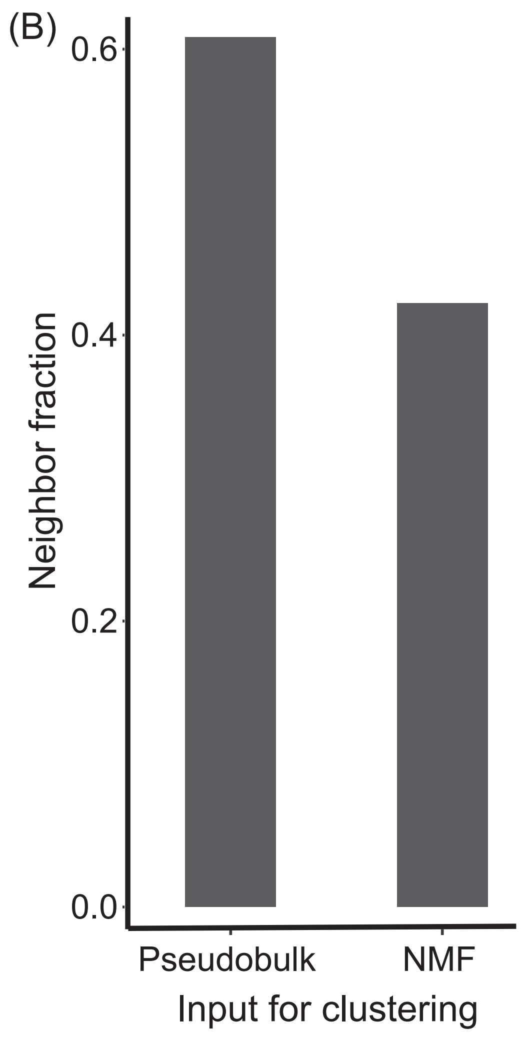

(B)
0.6
0.4
0.2
0.0
Neighbor fraction
Pseudobulk
NMF
Input for clustering

**Figure S1. Clustering robust NMF programs.** (**A**) Diagram describing the clustering process for robust NMFs. 5547 robust NMF programs were selected for initial clustering, out of which 3163 programs formed clusters and participated in the meta-programs (Table S2). See methods for further description. (**B**) Comparison between the fractions of neighboring programs (i.e. immediately adjacent in the hierarchical clustering dendrogram tree) that originate from the same study, in the NMF vs. the pseudobulk analysis. The fraction of in the NMF approach is significantly lower ($P$=1.84e$^{-31}$).

Figure S2

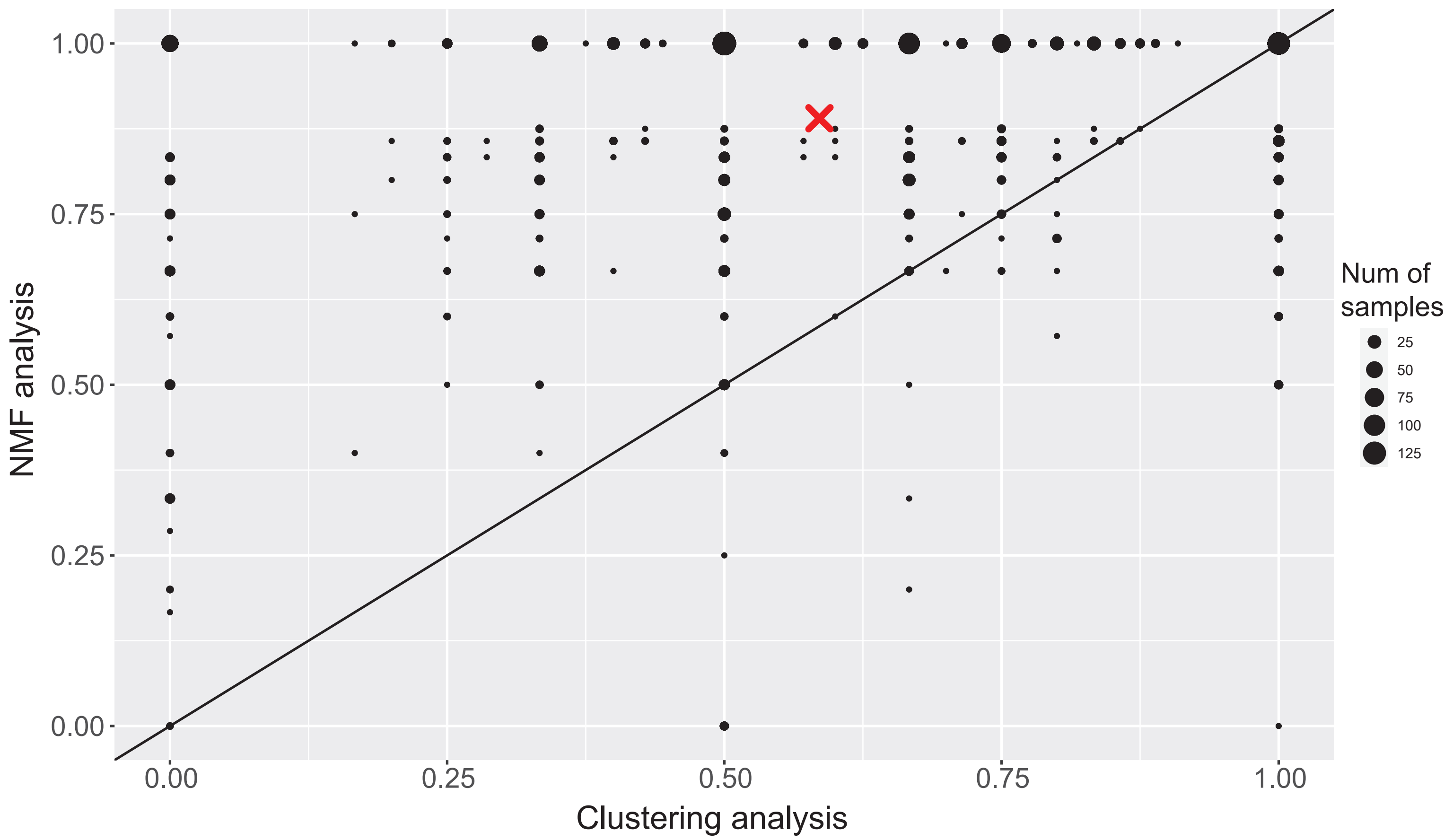

1.00
0.75
0.50
0.25
0.00
NMF analysis
0.00
0.25
0.50
0.75
1.00
Clustering analysis
Num of samples
25
50
75
100
125

**Figure S2. Fraction of ITH programs (defined by NMF or by Louvain clustering) that may be accounted for by the meta-programs.** The Y-axis shows, for each sample, the percentage of robust NMF programs identified in that sample that have a significant positive score for at least one MP. The X-axis shows a similar analysis with an alternative method to define ITH programs, based on Louvain clustering (with k=10). Cells in each Louvain cluster were scored against the MPs, and the fractions of clusters from each sample where more than 50% of the cells scored positive was computed. Since may samples had the exact same X,Y values, the number of samples in each coordinate is refleted by the size of the circle, as noted in the legend at the right.

Figure S3

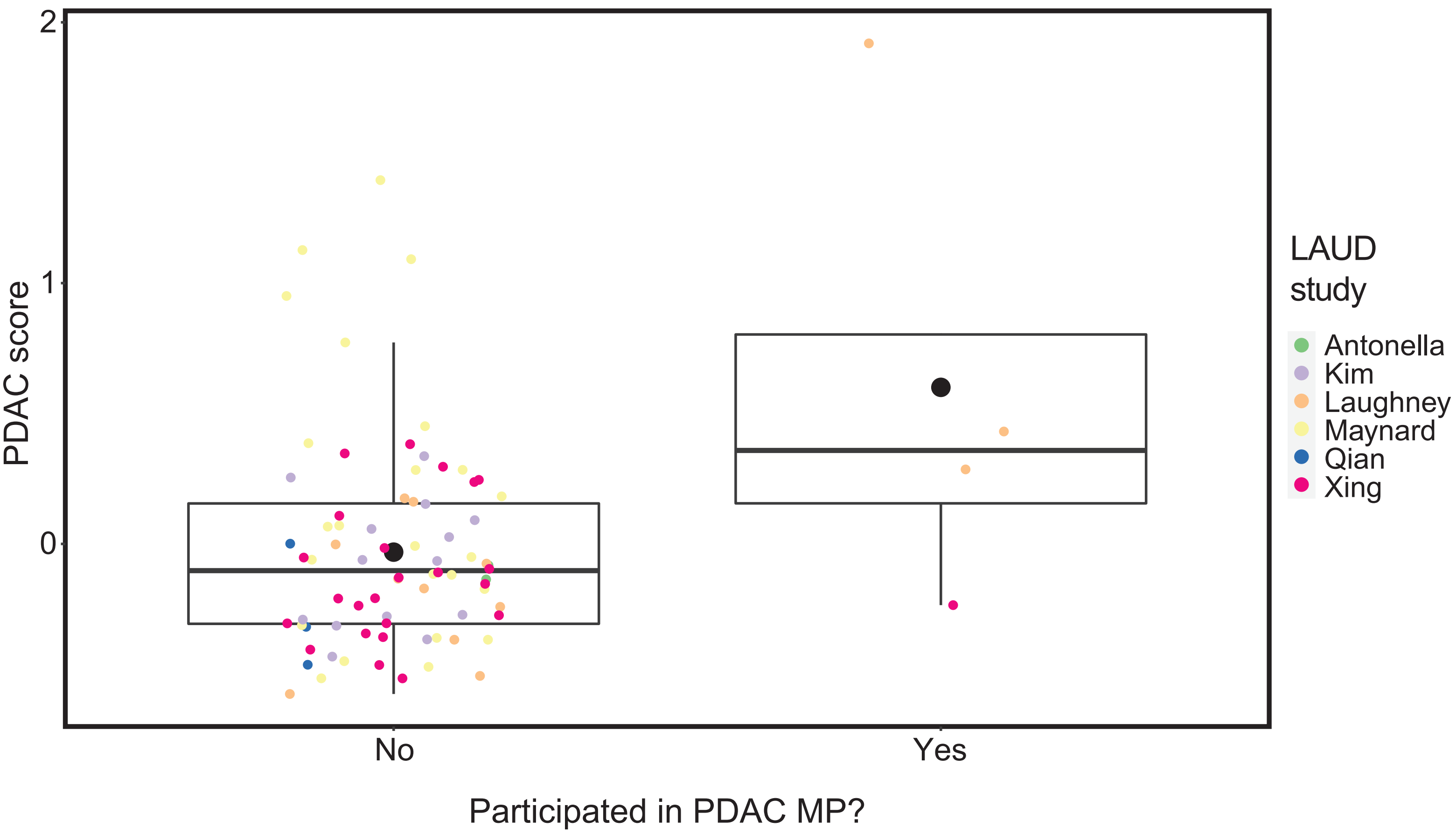

2
1
0
PDAC score
No
Yes
Participated in PDAC MP?
LAUD study
Antonella
Kim
Laughney
Maynard
Qian
Xing

**Figure S3. Overall expression of PDAC genes in lung adenocarcinoma (LUAD) samples.** The top 100 genes by higher expression in PDAC than in LUAD samples were defined as a PDAC signature. Average expression of the PDAC signature genes in malignant cells from each tumor was defined as a PDAC score. The PDAC scores were centered and are shown by box plots for two subsets of LUAD samples, based on whether MP30 (PDAC-classical) was identified as variable within these tumors. LUAD Samples with MP30 variability had significantly higher PDAC scores ($P$=0.002, t-test). Sample are also colored by their study as indicated at the right legend.

Figure S4, page 1

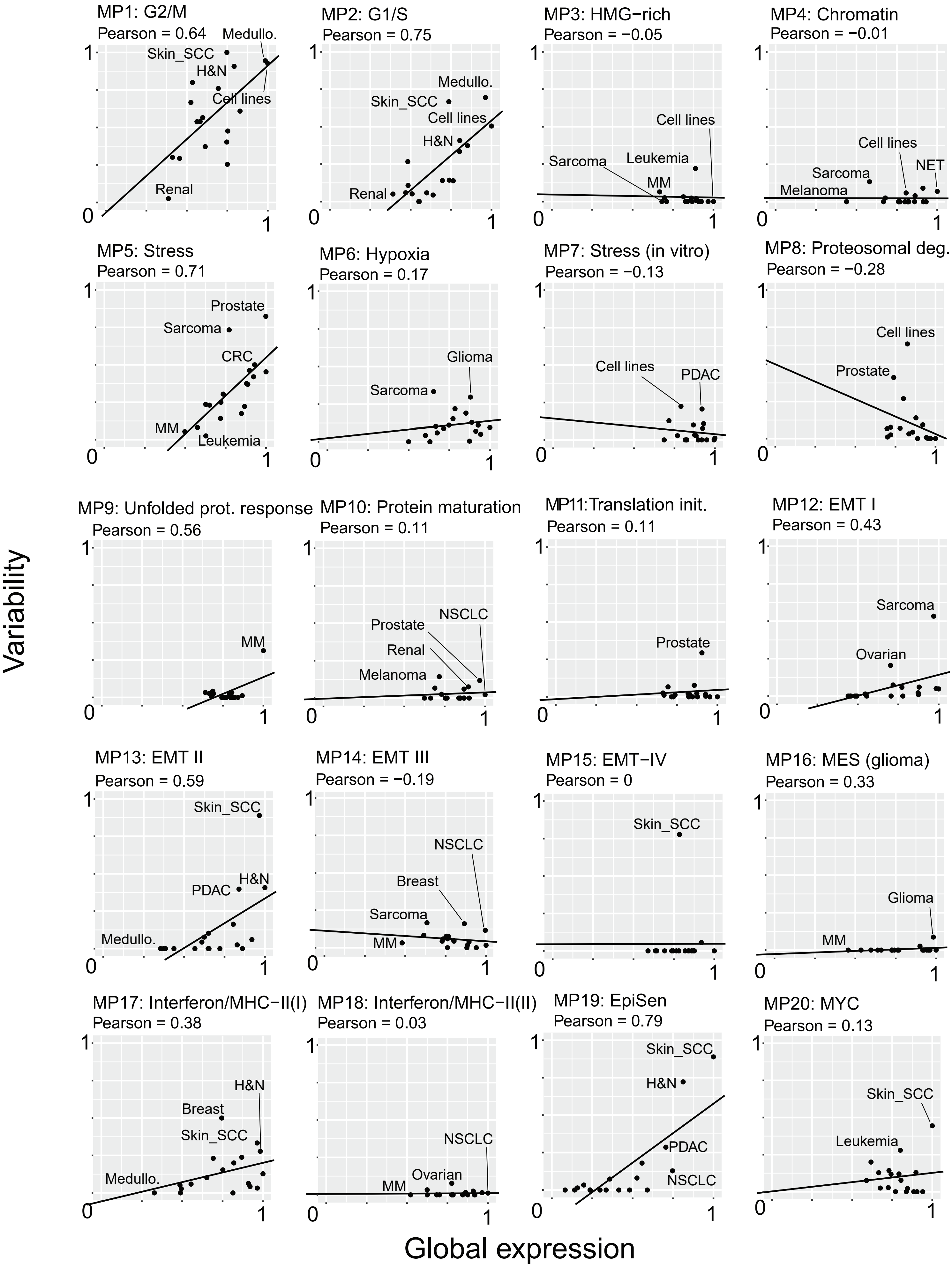

Figure S4, page 2

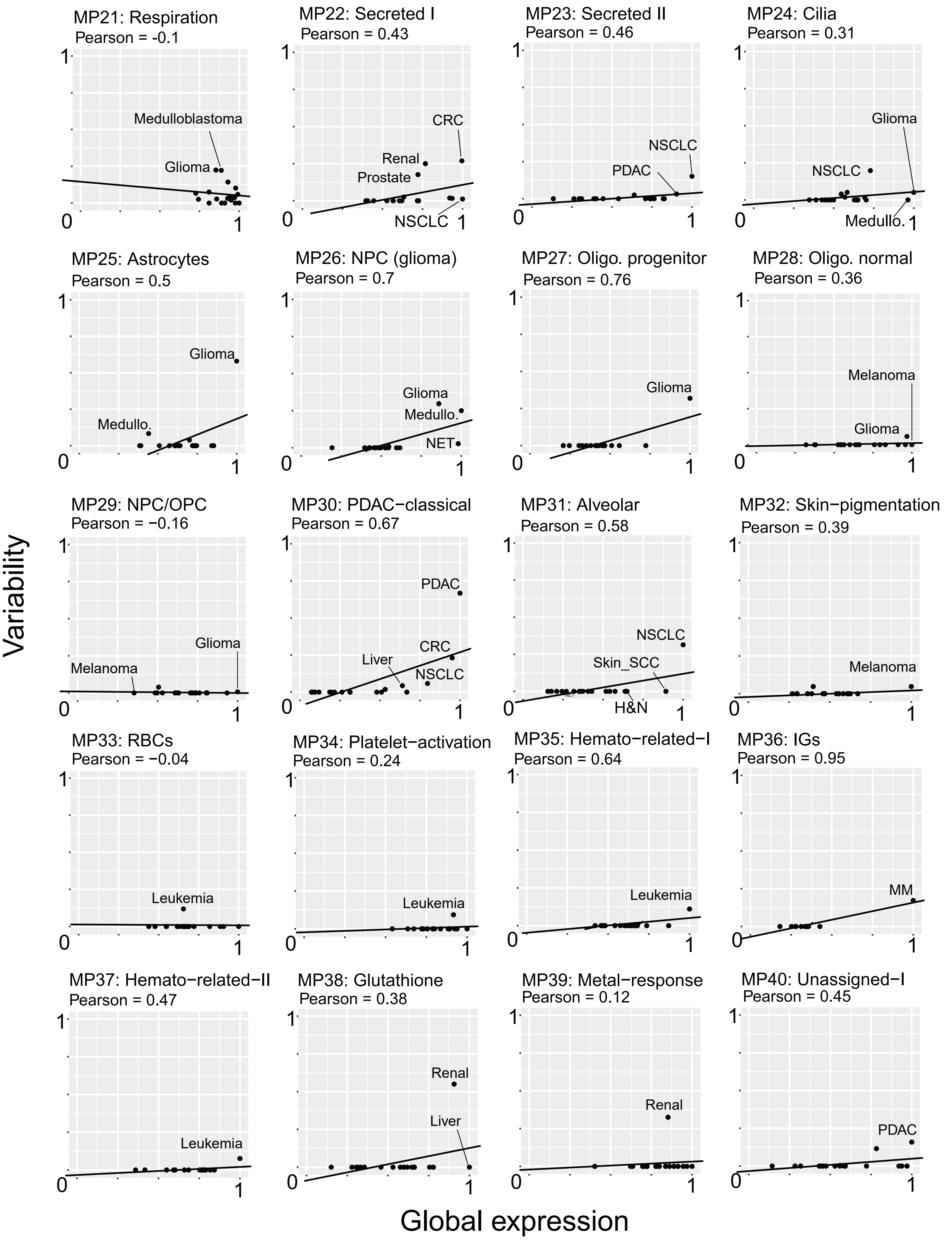

Figure S4, page 3

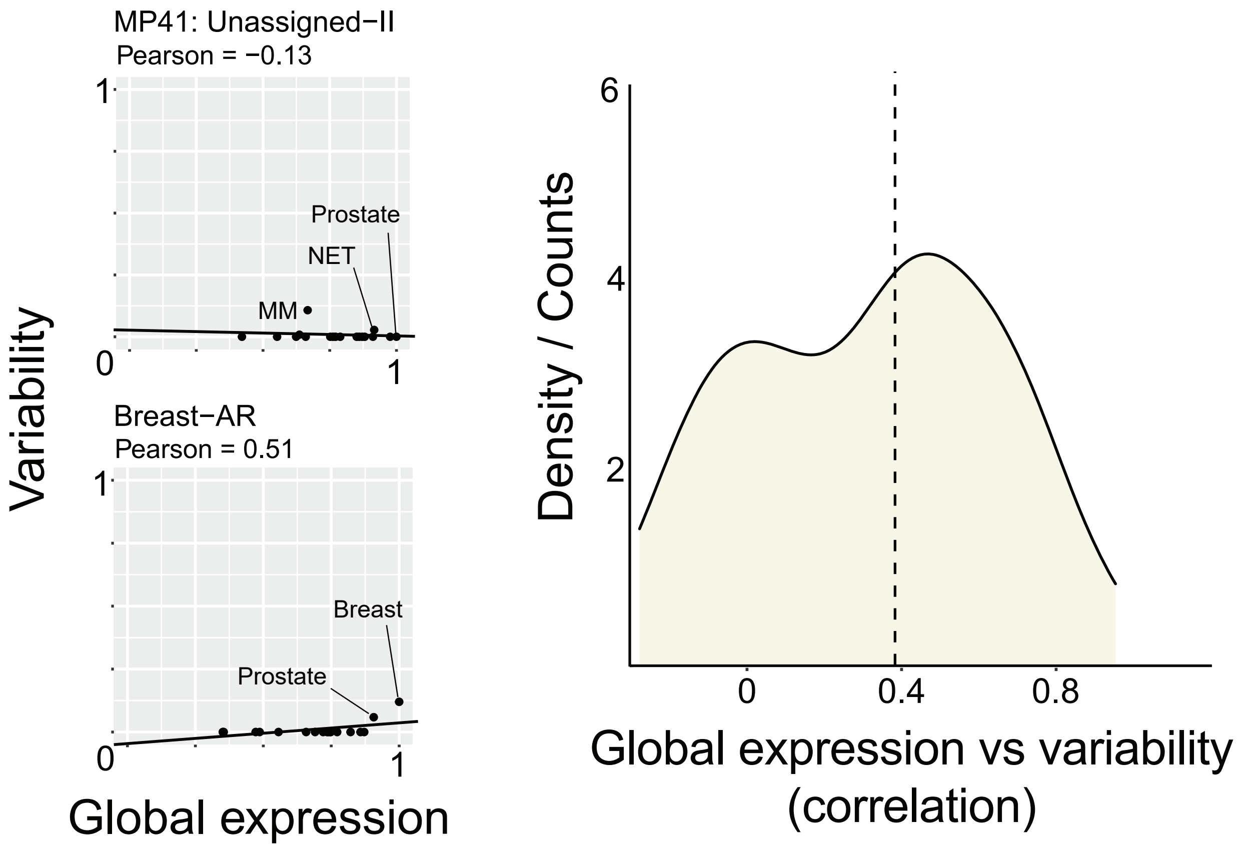

**Figure S4. Global (mean) expression vs. variability of MP expression across different cancer types.** Each panel shows global expression (X-axis) and variability of expression (Y-axis) of a specific MP across all cancer types analyzed (dots), along with a Pearson correlation and labels for selected cancer types. Global expression is defined as the mean expression of the MP genes across malignant cells from all samples of a particular cancer type; global expression is further normalized into [0..1]. Variablity of the MP in a given cancer type is defined as the fraction of tumors of that cancer type in which at least one of the robust NMF programs was assigned to the MP. The last panel shows the distribution of correlations between global expression and variability across all MPs, with a dashed line indicating the median correlation (R=0.38). See methods for further description.

Figure S5

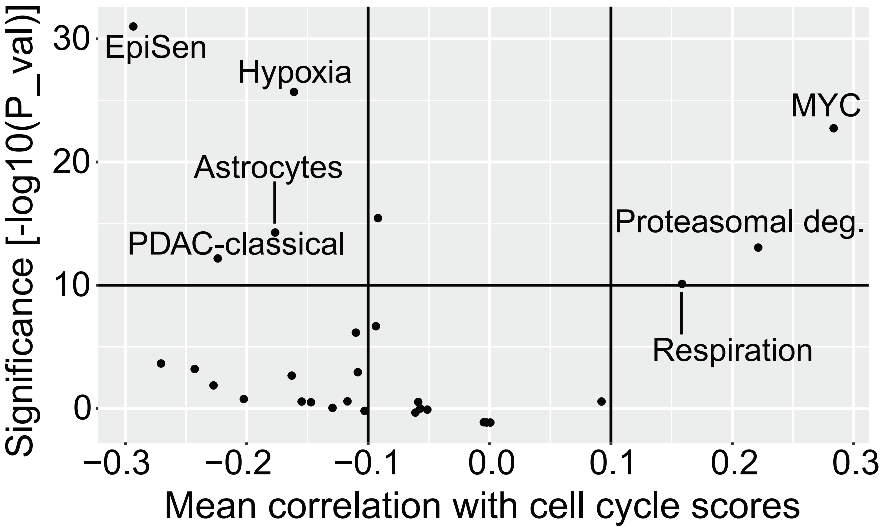
EpiSen
Hypoxia
MYC
Astrocytes
PDAC-classical
Proteasomal deg.
Respiration
Significance [-log10(P_val)]
30
20
10
0
−0.3
−0.2
−0.1
0.0
0.1
0.2
0.3
Mean correlation with cell cycle scores

**Figure S5. Meta-program associations with proliferation.** For each sample, the correlation between cell scores for the cell cycle MPs (maximum over the scores for MP1-4) and to different MPs was calculated. The mean values of these correlations are shown for different MPs against the significance (obtained by a one-sample t-test). The horizontal and vertical lines delineate the thresholds above (or below) which we considered values to be significant.

### Figure S6

(A)

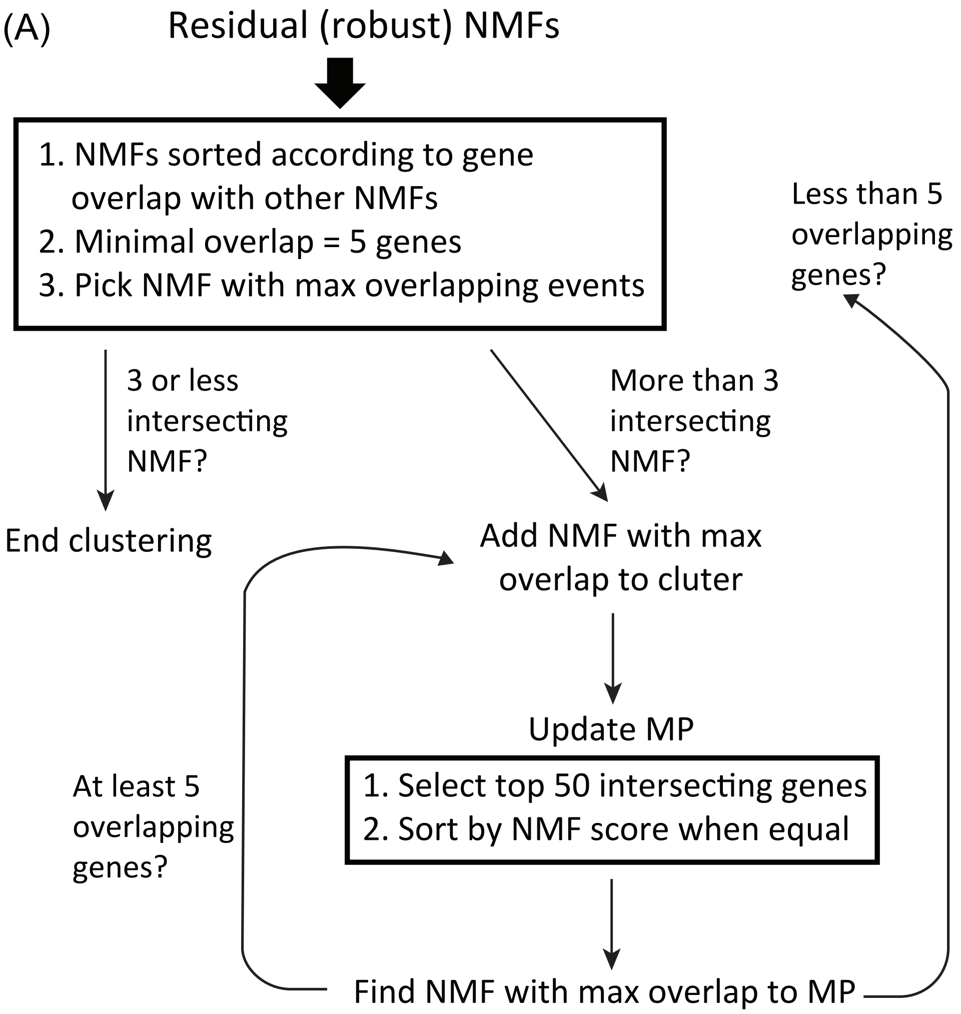


(B)

AR (breast) MP

| Cancer type | Sample | Subtype |
|---|---|---|
| Breast | 0029_9C (Pal) | ER+ |
| Breast | LN_0167 (Pal) | ER+ |
| Breast | 0176 (Pal) | HER2 |
| Breast | 0308 (Pal) | HER2 |
| Breast | 0337 (Pal) | HER2 |
| Breast | 89 (Karaayvas) | TNBC |
| Breast | CTC (Jordan) | |
| Prostate | 13 (Chen) | |

(C)

Breast sample HER2_0176 (Pal et al 2021)

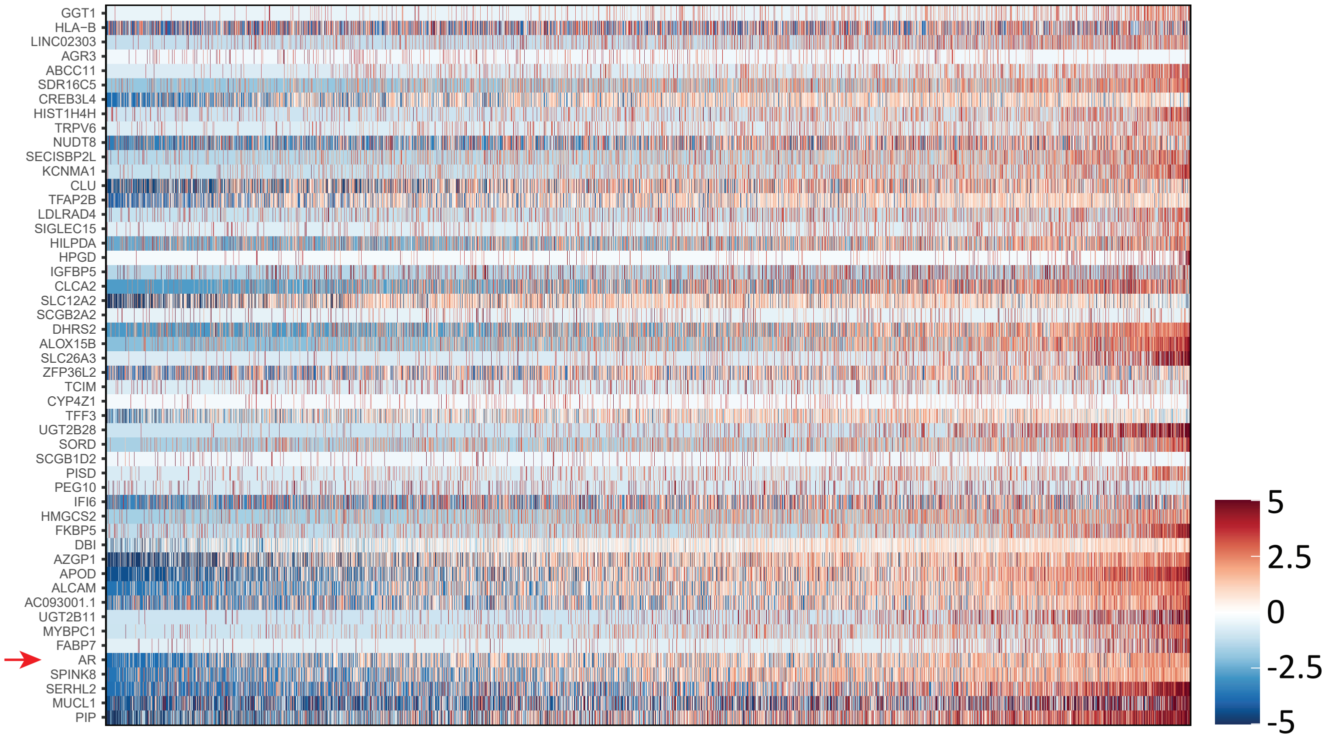

**Figure S6. Residual meta-program generation.** (**A**) Similar to the process described in figure S1A, the residual robust NMFs that were not included in the initial MPs were clustered using more lenient parameters. 59 residual MPs were generated (**Table S2**). (**B**) Residual MP50 was mostly identified within breast cancer samples and was enriched for the Hallmark Androgen Response signature (**Table S3**). (**C**) Heatmap for the HER2_0176 breast cancer sample from Pal et al. (2021), showing relative expression of residual MP50 genes (rows) across malignant cells (columns) ordered by MP50 expression. The Androgen Receptor (AR) gene is highlighted by a red arrow.

**Table S1.** TME composition across studies. 7 common cells types are seen across many of the studies. Additional 31 cell subtypes are defined in a smaller fraction of studies (sometimes only one study). Several cell subtypes that were context specific were combined or removed.

**Table S2.** 41 meta-programs, robust NMFs that were used to generate the MPs and the residual MPs defined using robust NMFs that were not initially assigned to a MPs running the clustering algorithm with more lenient parameters.

**Table S3.** Signatures for the MPs and residual MPs. Signatures are sorted from top to bottom for each MP according to corrected p-value (only significant signatures after correction for multiple comparisons are shown). Signatures that were chosen for figure 2C are highlighted in red.

**Table S4.** Hallmark distribution across different cancer types. The main panel in figure 4 depicts the distribution across solid tumors (first column).

**Table S5.** 16 T-cell meta-programs and their top signatures sorted from top to bottom according to corrected p-value (only significant signatures after correction for multiple comparisons are shown).
