## Supplemental Note 1 for "The transcriptional hallmarks of intra-tumor heterogeneity across a thousand tumors"

### Supplementary Note 1: detailed description of meta-programs

**Table of contents**

Each page contains text description and an example heatmap (genes X cells) of one MP.

MP1: Cell cycle (G2/M) ……………….....2
MP2: Cell cycle (G1/S) ……………………..3
MP3: Cell cycle (HMG-rich)………………..4
MP4: Cell cycle (chromatin)…………….....5
MP5: Stress……………………………..6
MP13: EMT-II……………………….....14
MP14: EMT-III……………………….....15
MP15: EMT-IV………………………..16
MP16: MES (glioma)…………………17
MP17: Interferon/MHC-II (#1)………..18
MP18: Interferon/MHC-II (#2)………..19
MP19: Epithelial senescence (EpiSen)..20
MP20: MYC-targets……………………..21
MP21: Respiration……………………22
MP22: Secreted I……………………....23
MP23: Secreted II……………………...24
MP24: Cilia……………………………25
MP25: Astrocytes……………………..26
MP26: NPC (glioma)………………….27
MP27: Oligo. progenitor……………….28
MP28: Oligo. Normal………………….29
MP29: NPC/OPC………………………30
MP30: PDAC classical………………....31
MP31: Alveolar……………………...32
MP32: Skin pigmentation……………....33
MP33: RBC…………………………….34
MP40: Unassigned I……....……………...41
MP41: Unassigned II…………………...42

#### MP1: Cell cycle (G2/M)

**Brief description:**
*General* cell cycle program reflecting the G2/M phases.

**Functional enrichments:**
Highly enriched with many cell cycle related functional annotations including H.HALLMARK_G2M_CHECKPOINT

**Selected genes:**
MKI67 (standard cell cycle marker); CCNB1 and CCNB2 (cycling B); TOP2A (topoisomerase); CENPA, CENPE and CENPF (centromere).

**Additional comments:**
The most frequently detected meta-program, highly similar across different contexts and highly consistent with programs defined in previous studies.

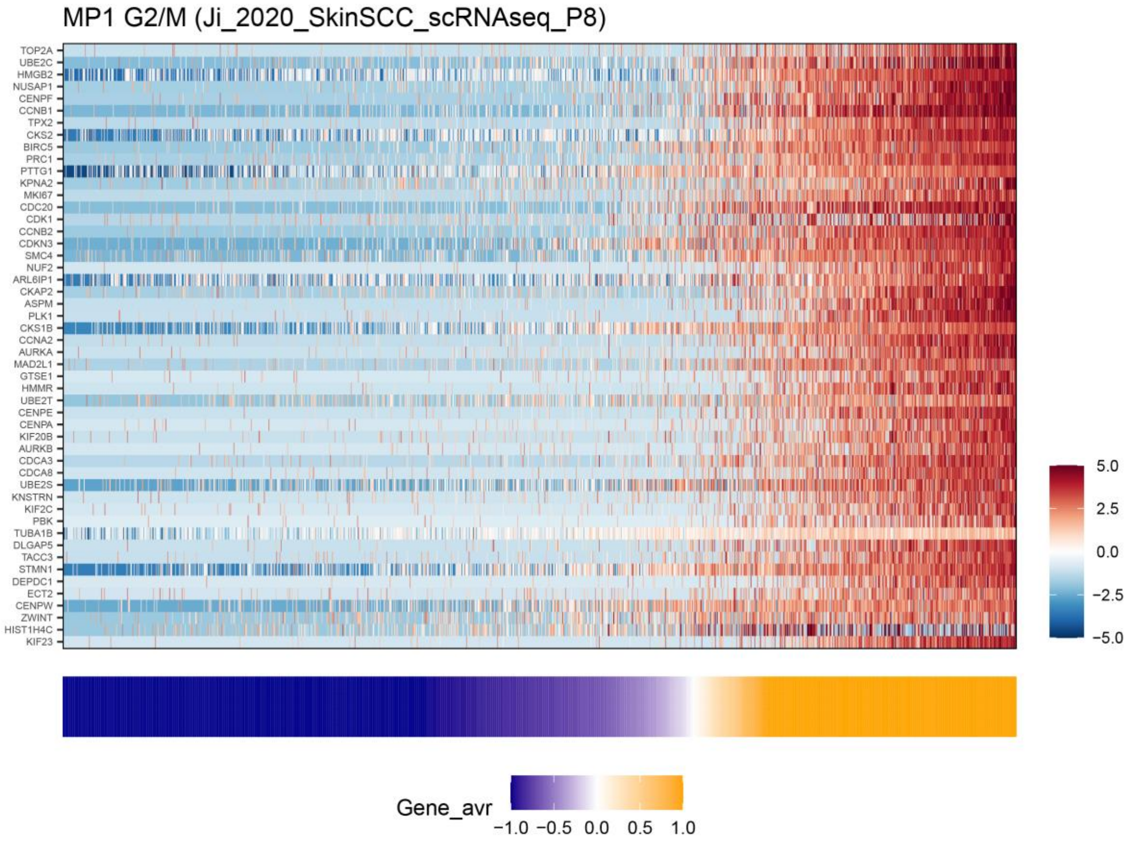

<u>**MP2: Cell cycle (G1/S)**</u>

**Brief description:**
Canonical cell cycle program reflecting the G1/S phases.

**Functional enrichments:**
Highly enriched with many cell cycle related functional annotations including C5.GOBP_CELL_CYCLE_G1_S_PHASE_TRANSITION

**Selected genes:**
PCNA (replication marker); MCM3-7 (MCM complex); GMNN (Geminin).

**Additional comments:**
The second most frequently detected meta-program, highly similar across different contexts and highly consistent with programs defined in previous studies.

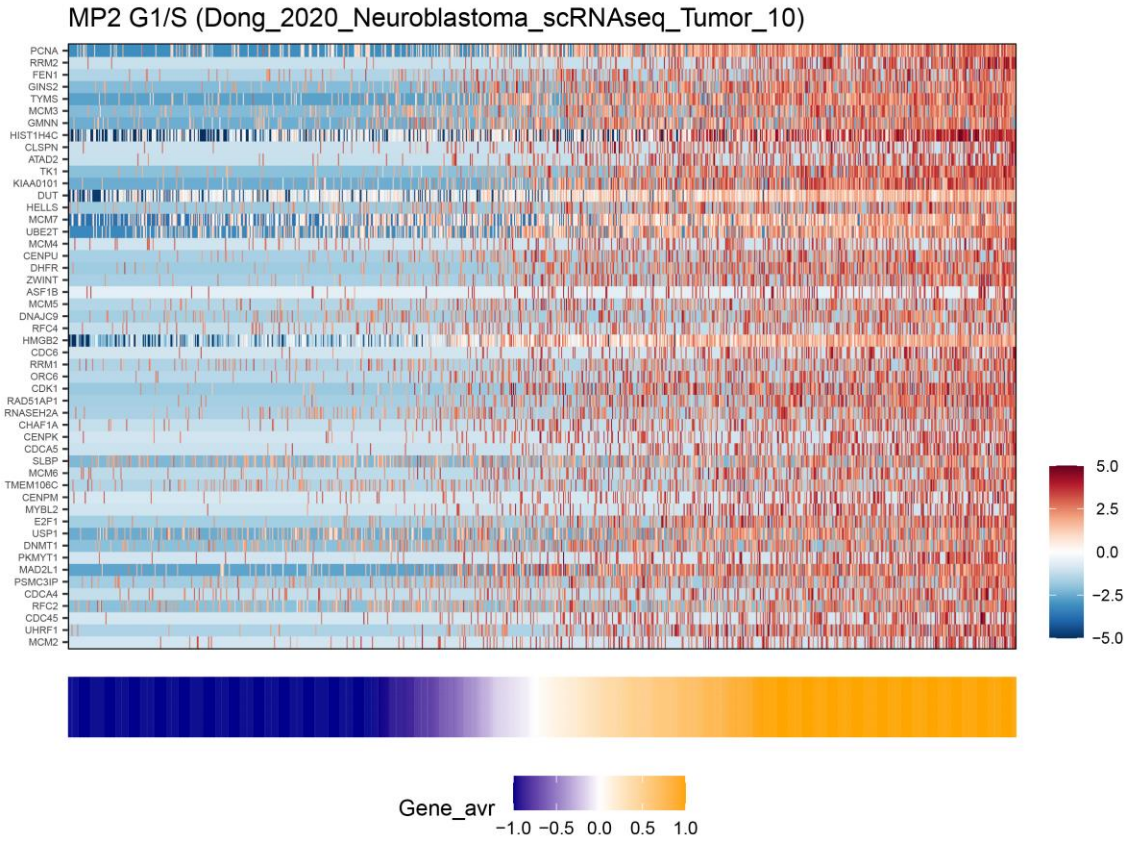

#### MP3: Cell cycle (HMG-rich)

**Brief description:**
Rare cell cycle program enriched with HMG-box proteins.

**Functional enrichments:**
Enriched with many cell cycle related functional annotations (H.HALLMARK_E2F_TARGETS), but also with annotations related to HMG-box proteins including C5.GOBP_DNA_GEOMETRIC_CHANGE.

**Selected genes:**
HMGA1, HMGB1, HMGB2, HMGN1 and HMGN2 (HMG-box proteins); PCNA, MCM7, CKS1B (cell cycle genes); H2AFV, H2AFZ, HIST1H4C (histones); HSPB11, HSPD1, HSPE1 (heat-shock proteins).

**Additional comments:**
Mostly observed in hematological malignancies but also found rarely in other cancer types (lung and pancreatic). Previously undescribed to our knowledge.

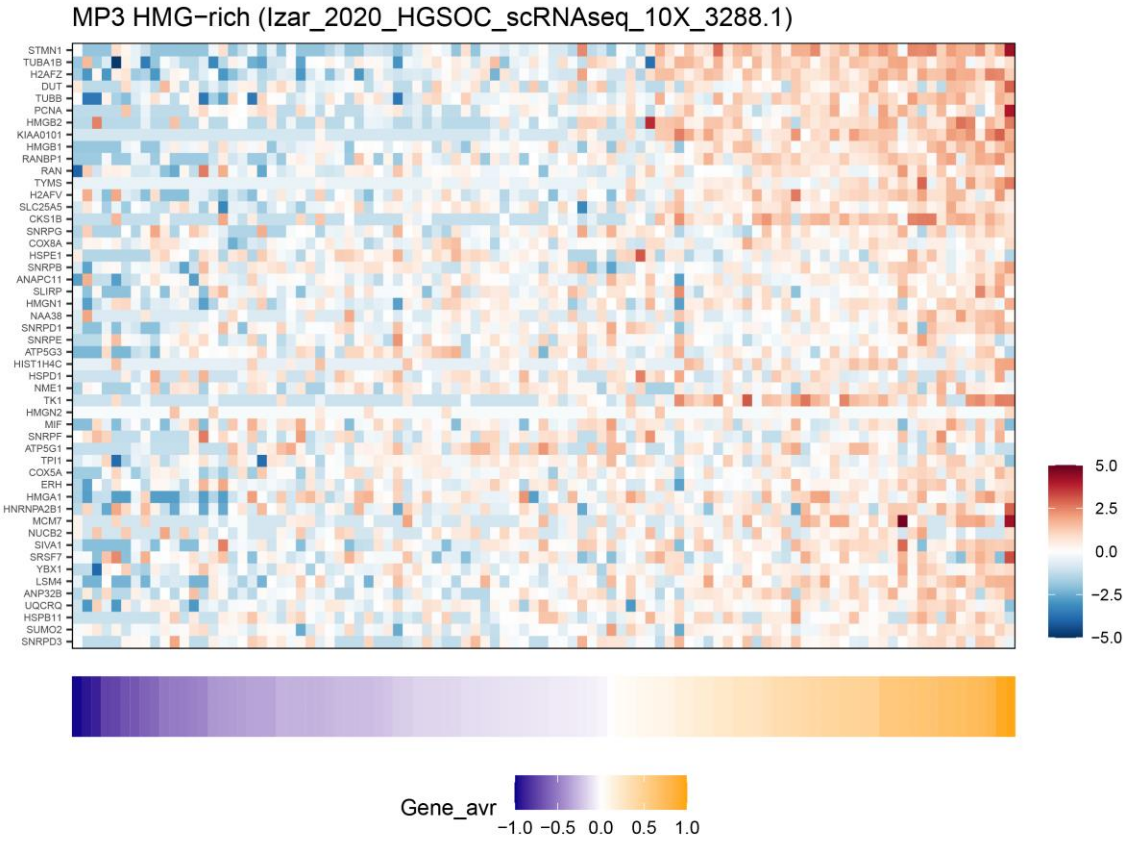

### MP4: Cell cycle (Chromatin)

**Brief description:**

*Shared* MP that is highly enriched with chromatin-related genes, including histones and chromatin regulators. Many of these genes are functionally important during the cell cycle and/or are also upregulated during the cell cycle (e.g. histones and SMC proteins) and therefore we speculate that this program characterizes a subset of cycling cells.

**Functional enrichment:**

Enriched with many chromatin related functional annotations (e.g. C5.GOBP_CHROMATIN_ORGANIZATION_INVOLVED_IN_REGULATION_OF_TRANSCRIPTION) and to epigenetic related functional annotation (e.g. C5.GOBP_REGULATION_OF_GENE_EXPRESSION_EPIGENETIC).

**Selected genes:**

Chromatin regulators (SMARCA5, PHF3, ASH1L, SETD2, SUZ12, KMT2A, ATAD2, ATAD5), histone genes (HIST1H1C, HIST2H2AC, HIST1H1B), SMC DNA-repair complex (SMC1A, SMC5, SMCHD1).

**Additional comments:**

Mostly observed in cell-lines, but also abundant in osteosarcoma and neuroendocrine tumors. Previously undescribed to our knowledge.

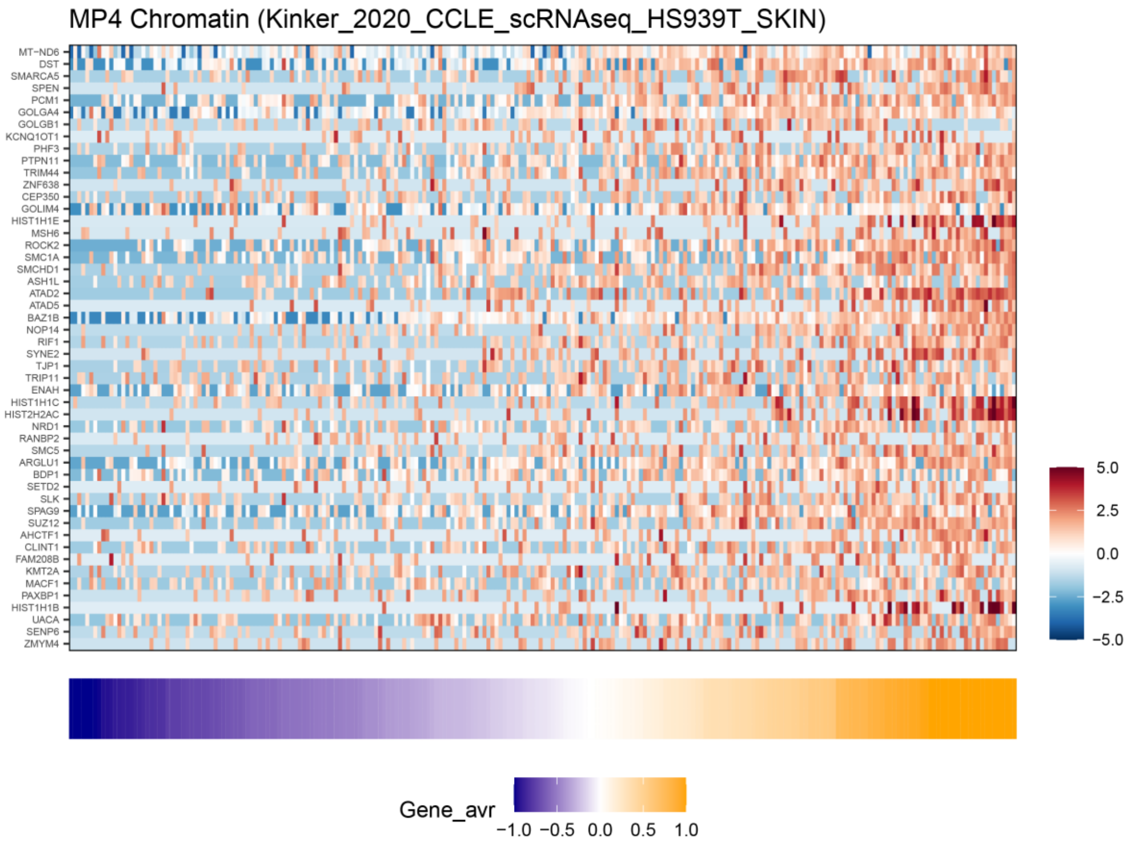

### MP5: Stress

**Brief description:**
Canonical stress-related program.

**Functional enrichments:**
Enriched with many stress-related functional annotations, including C5.GOBP_REGULATION_OF_DNA_TEMPLATED_TRANSCRIPTION_IN_RESPONSE_TO_STRESS, H.HALLMARK_UV_RESPONSE_UP, H.HALLMARK_APOPTOSIS, H.HALLMARK_HYPOXIA,

**Selected genes:**
FOS, FOSB, JUN and JUNB (AP-1 transcription factors); IER2, IER3, EGR1 and ATF3 (immediate early genes); HSPA1A, HSPA1B and DNAJB1(heat-shock proteins).

**Additional comments:**
Observed broadly. Could be associated with tumor dissociation, but at least in certain cases it is observed also in the absence of dissociation. Described and investigated by Baron et al. (2020).

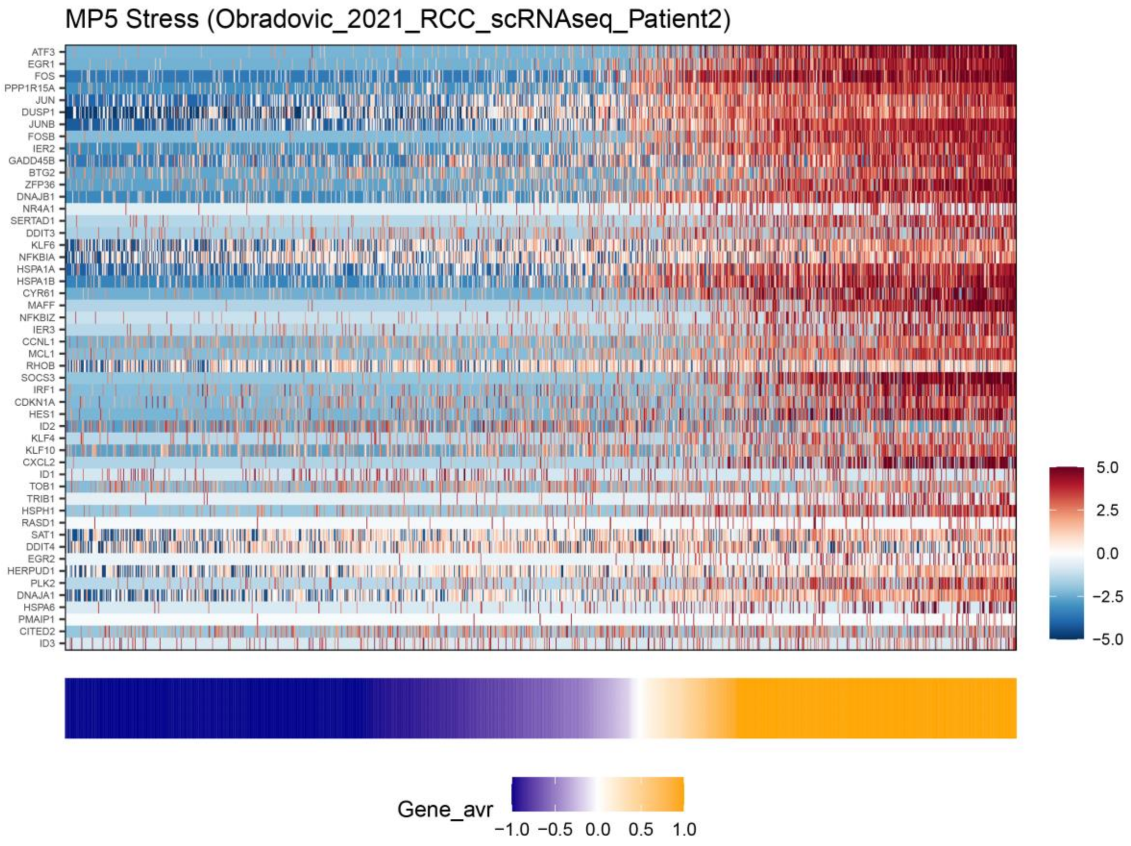

#### MP6: Hypoxia

**Brief description:**
*General* MP that belongs to the stress hallmark and is enriched with hypoxia-related genes, including hypoxia response and glycolysis.

**Functional enrichment:**
Enriched with many hypoxia related functional annotations (e.g. H.HALLMARK_HYPOXIA) or oxygen related metabolic processes (e.g. C5.GOBP_ATP_METABOLIC_PROCESS).

**Selected genes:**
hypoxia response (VEGFA, HILPDA, EGLN3), stress (NDRG1, DDIT3, DDIT4), glycolysis (ENO1, ENO2, LDHA, SLC2A1)

**Additional comments:**
Significantly abundant in glioma, and also abundant in breast, PDAC, sarcoma and thyroid cancer. Not observed in hematological malignancies, cells lines or CTCs. Variants of such hypoxia response has been described by many previous studies.

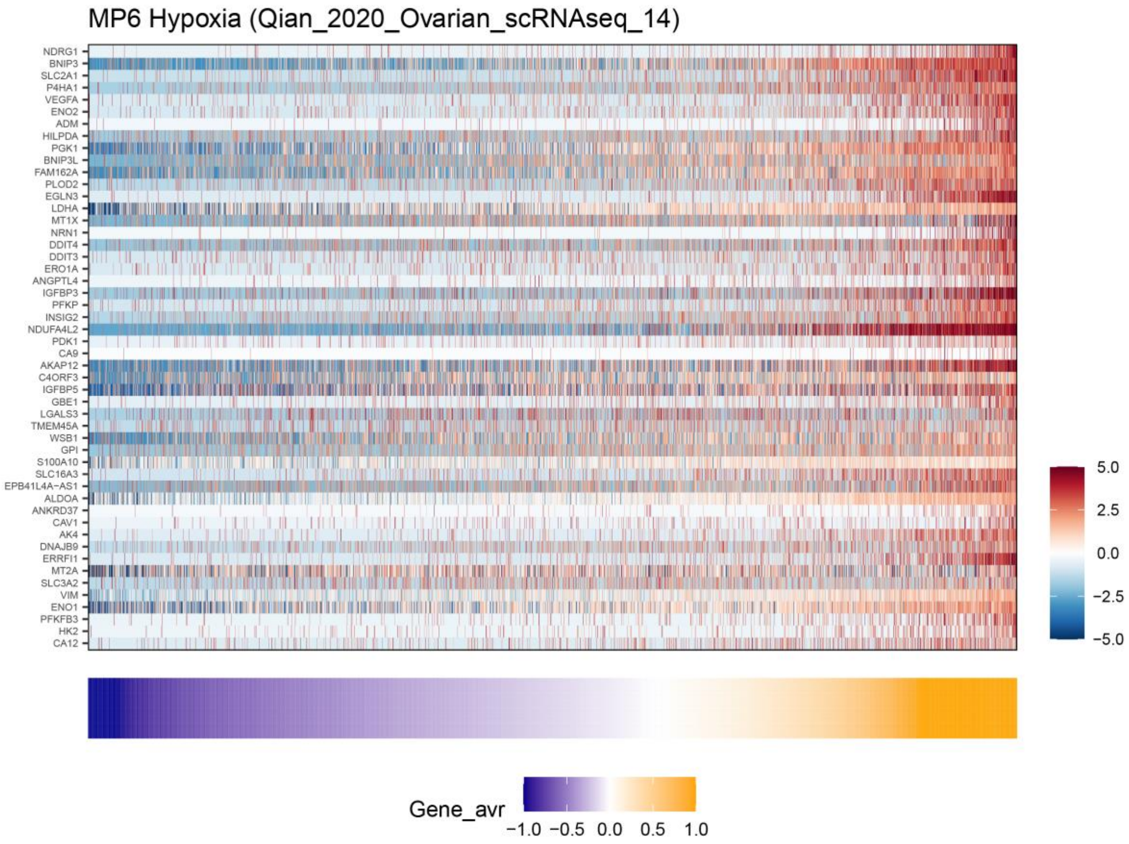

**<u>MP7 – Stress (in vitro)</u>**

<u>Brief description:</u> *Shared* stress MP seen mostly in cell-lines.

<u>Functional enrichment:</u> Enriched with stress related functional annotations (e.g. C5.GOBP_INTEGRATED_STRESS_RESPONSE_SIGNALING).

<u>Selected genes:</u> ATF4, ATF3, DDIT3, DDIT4, HSPA5, HSPA9, SQSTM1, XBP1, CEBPG, PPP1R15A.

<u>Additional comments</u>: Aside from cell-lines also observed in glioma and PDAC.

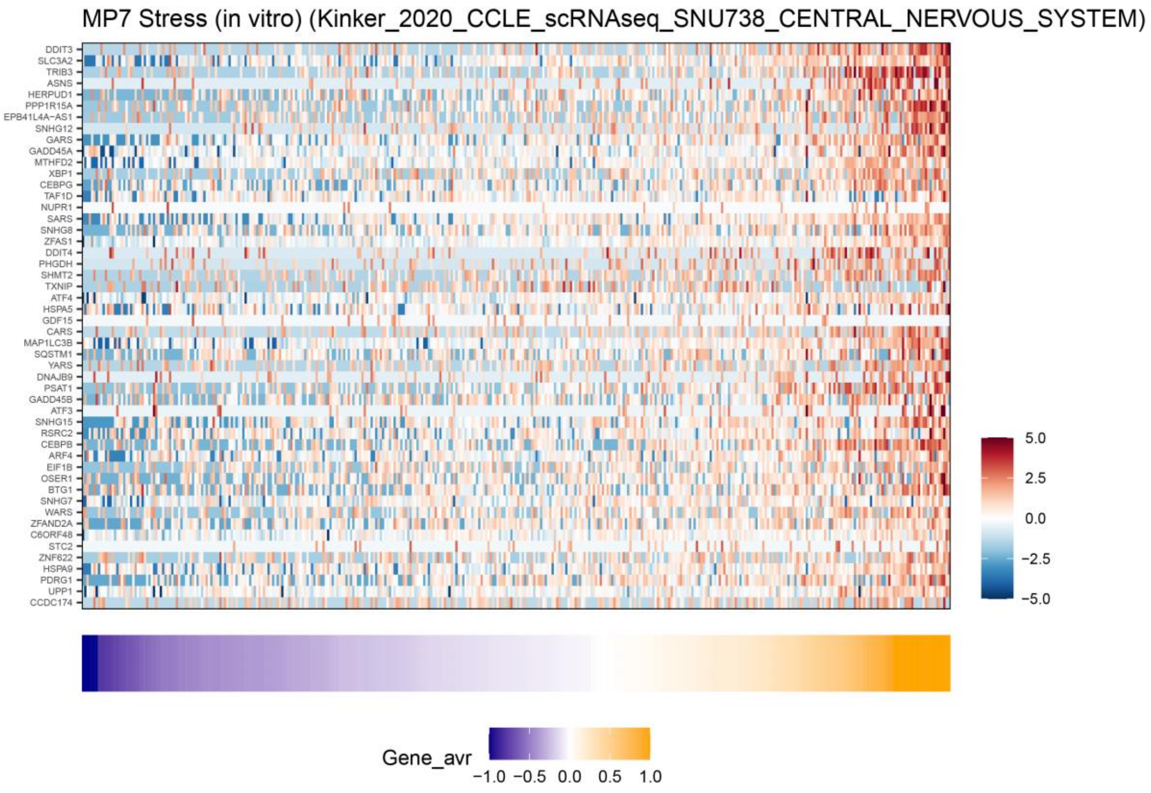

#### MP8 – Proteasomal degradation

Brief description: *Shared* MP that belongs to the protein-regulation hallmark and consists of many proteasome subunits.

Functional enrichment: Enriched with proteasomal catabolic related functional annotation (e.g. C5.GOBP_SCF_DEPENDENT_PROTEASOMAL_UBIQUITIN_DEPENDENT_PROTEIN_CATABOLIC_PROCESS).

Selected genes: proteasome subunits (PSMA3, PSMA4, PSMB1, PSMB3, PSMB6, PSMC2, PSMC4, PSMD13, PSME2), CCT complex (CCT5, CCT7, CCT8), translation initiation (EIF3I, EIF4A1, EIF4A3).

Additional comments: most abundant in cell-lines, followed by prostate cancer. Described previously as heterogeneous within multiple cell lines by Kinker et al.

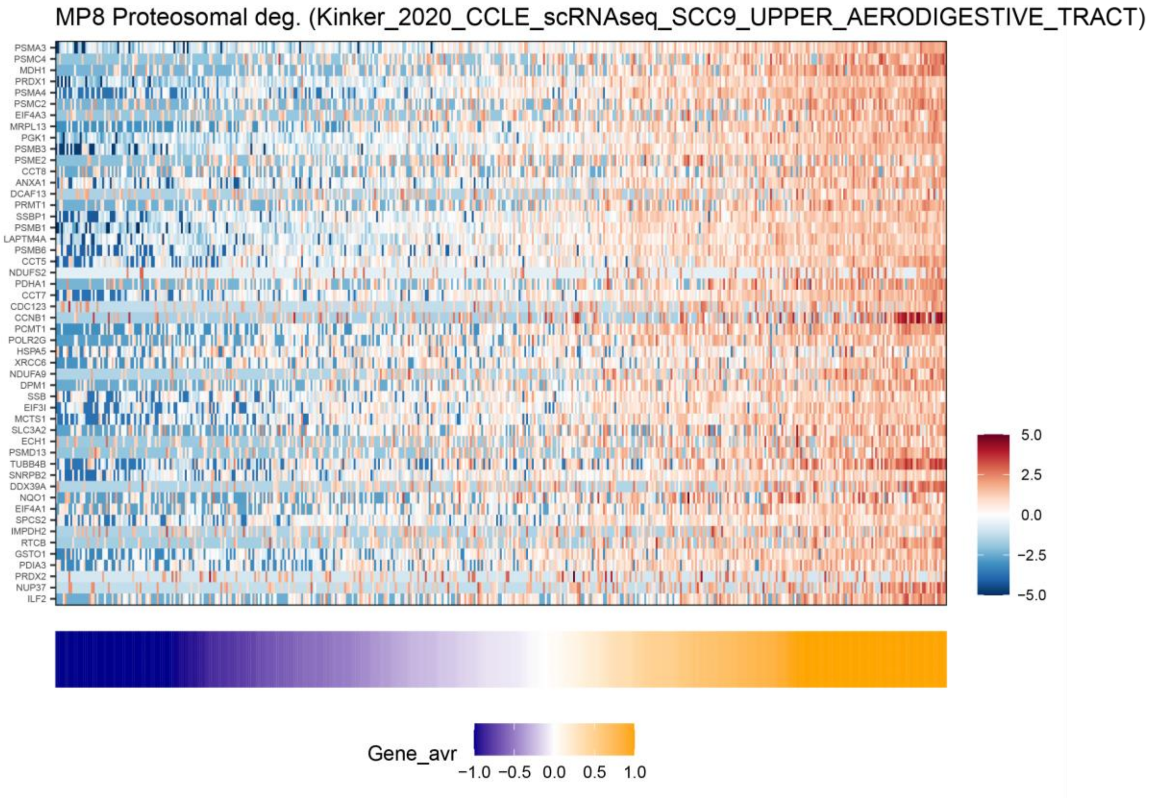

#### MP9 – Unfolded protein response

Brief description: *Shared* MP that belongs to the protein-regulation hallmark and consists of many genes involved in the unfolded protein response.

Functional annotations: Enriched with protein-folding related functional annotations (e.g. H.HALLMARK_UNFOLDED_PROTEIN_RESPONSE).

Selected genes: DNAJB9, DNAJB11, DNAJC3, HSPA5, HSP90B1, HSP90B2P, SRPRA, HYOU1, CALR,

Additional comments: Most abundant in multiple myeloma. Well known cellular response, but to our knowledge not previously described in the context of programs of intra-tumor heterogeneity.

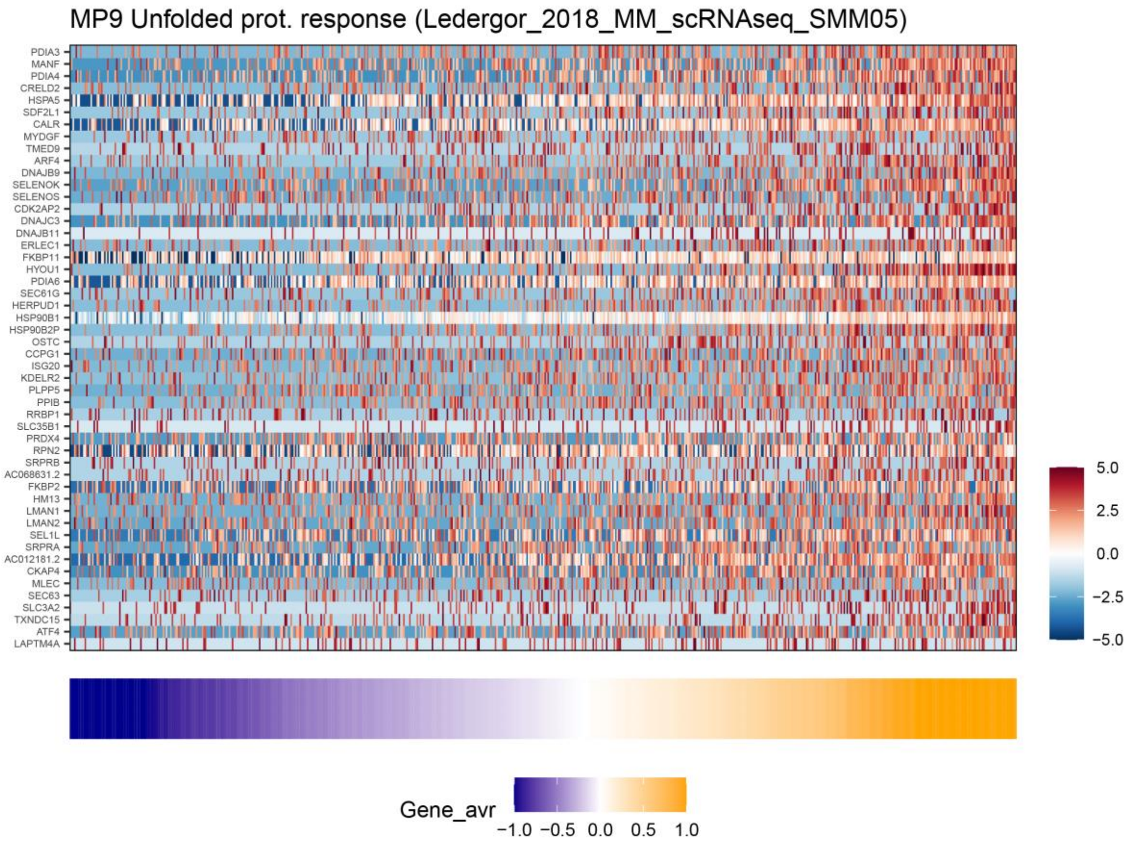

#### MP10 – Protein maturation

Brief description: *Shared* MP that belongs to the protein-regulation hallmark and consists of many ER genes involved in protein maturation.

Functional annotation: Enriched with endoplasmic-reticulum related functional annotations (e.g. C5.GOCC_ENDOPLASMIC_RETICULUM_LUMEN).

Selected genes: PDIA3, PDIA4, PDIA6, RPN2, PIGT, PPIB, OS9, HSPA5, LMAN2, SIL1, DDOST

Additional comments: Common in melanoma, renal and prostate cancer.

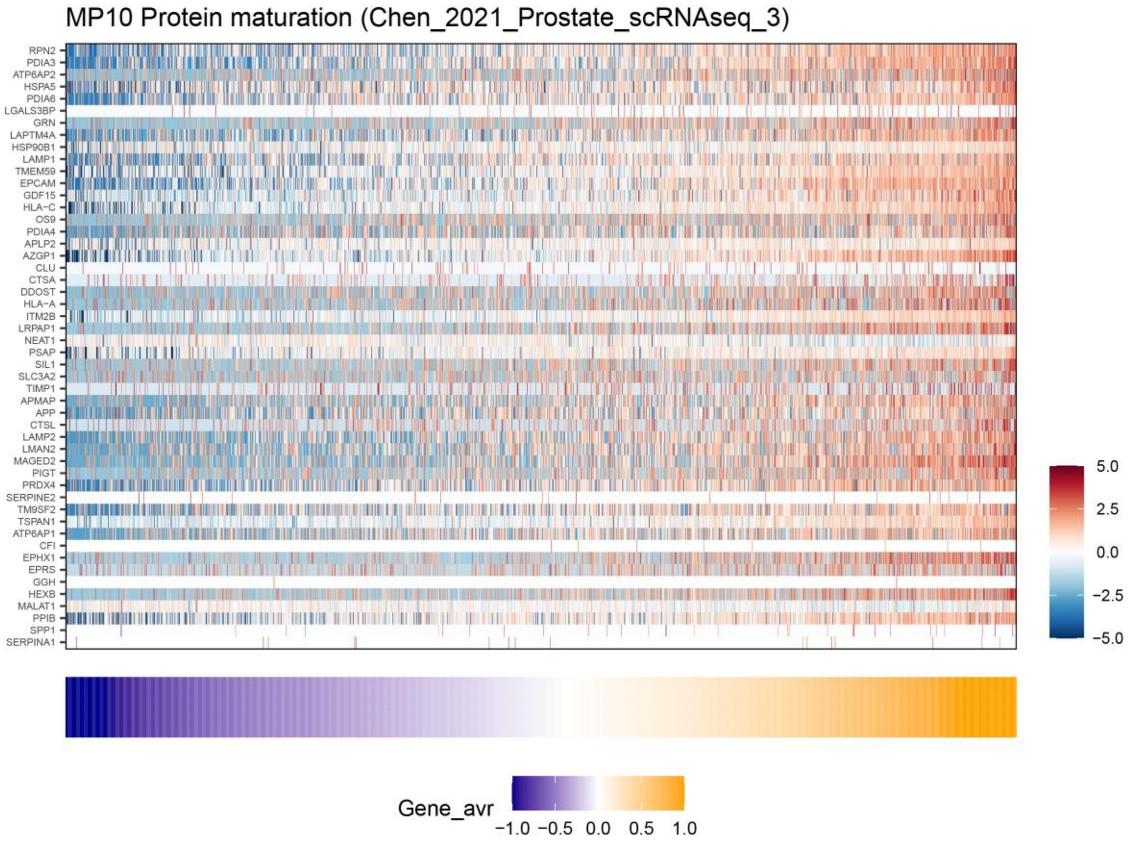

**MP11 – translation initiation**

Brief description: *Shared* MP that belongs to the protein-regulation hallmark and consists of many translation-related genes.

Functional annotation: Enriched with many translation-related functional annotations (e.g. C5.GOMF_TRANSLATION_INITIATION_FACTOR_ACTIVITY).

Selected genes: EIF2A, EIF4B, EIF3D, EIF3E, EIF3F, EIF2S3, EIF3M,

Additional comments: Most abundant in prostate cancer followed by colorectal-cancer.

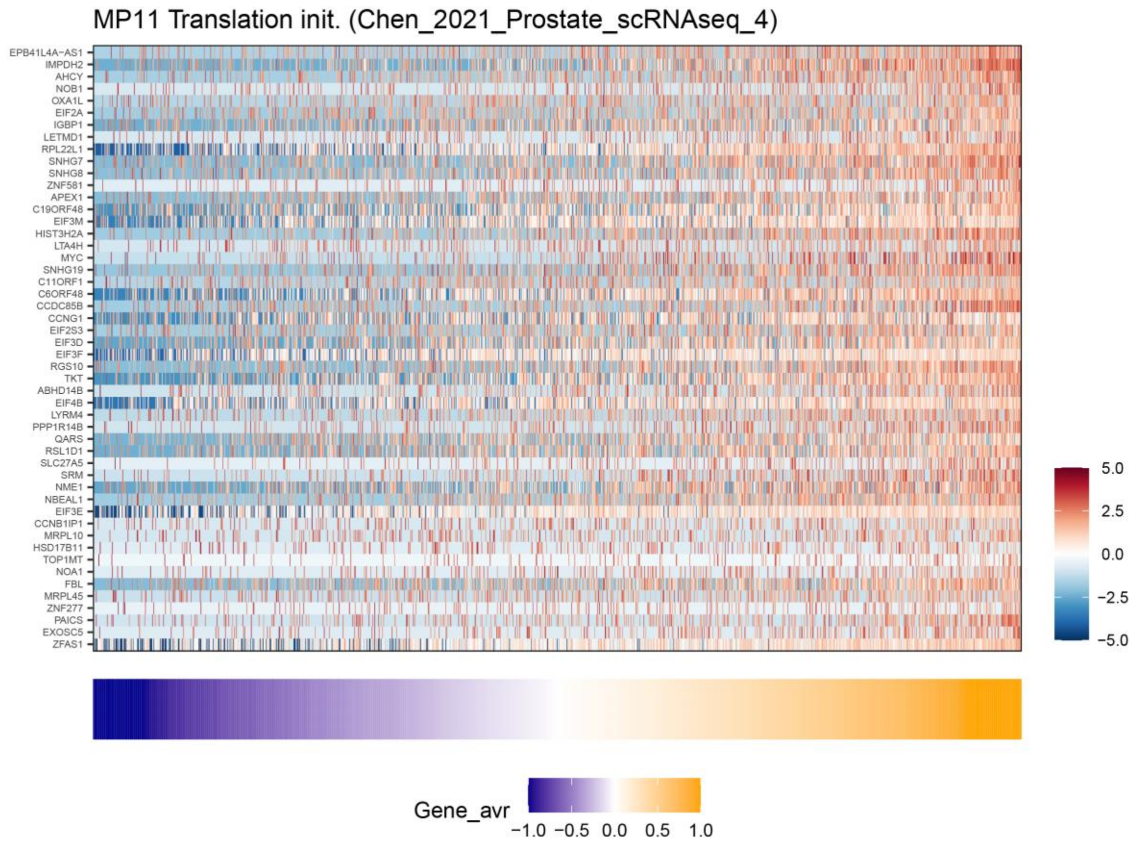

### MP12 – EMT-I

Brief description: *Shared* MP that belongs to the mesenchymal hallmark. Found in carcinoma and in sarcoma and hence might reflect more of a "full" mesenchymal program compared to other MPs that reflect a partial EMT.

Functional annotation: Enriched with epithelial-mesenchymal transition and extracellular functional annotations (e.g. H.HALLMARK_EPITHELIAL_MESENCHYMAL_TRANSITION).

Selected genes: collagen (COL1A1, COL1A2, COL3A1, COL4A1, COL4A2, COL6A1, COL6A2, COL6A), other mesenchymal markers (VIM, FN1, ACTA2).

Additional comments: Most abundant in sarcoma, followed by ovarian and thyroid cancer.

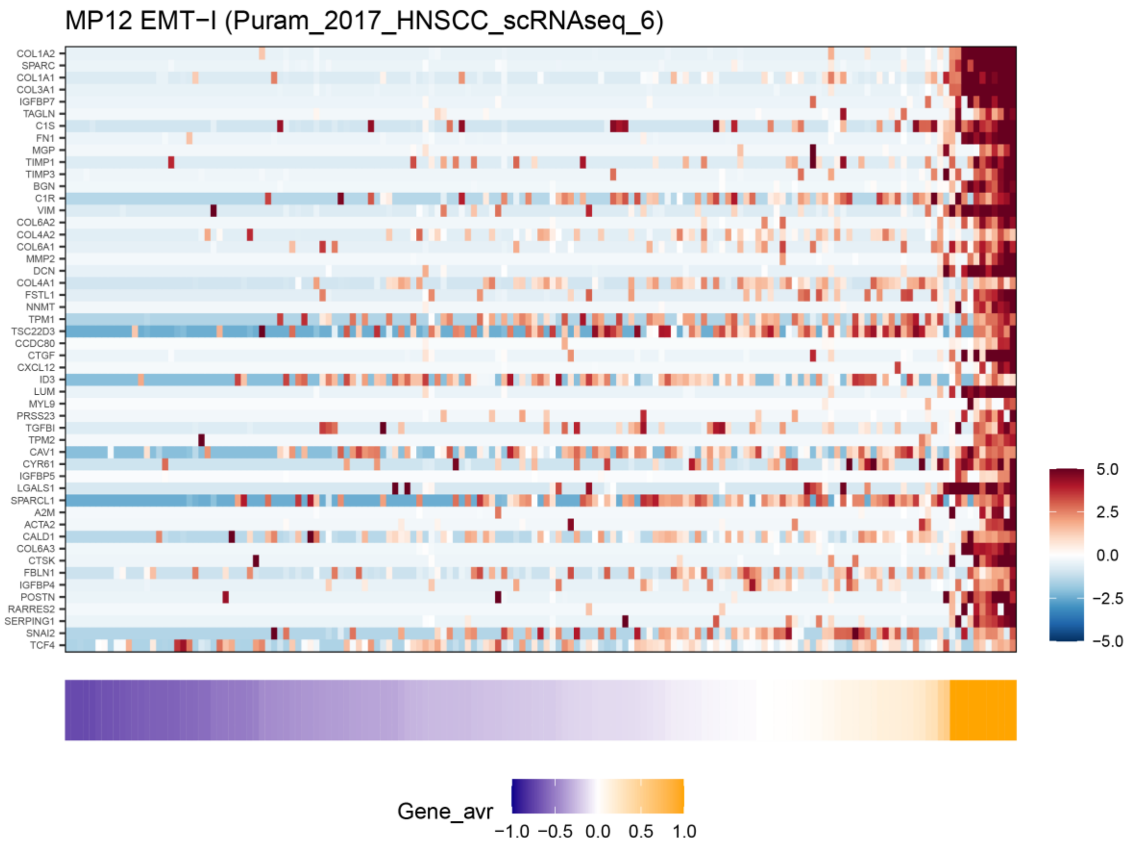

#### MP13 – EMT-II

Brief description: *Shared* MP that belongs to the mesenchymal hallmark and enriched in multiple types of squamous cell carcinoma.

Functional annotation: Enriched with epithelial-mesenchymal transition and extracellular functional annotations (e.g. H.HALLMARK_EPITHELIAL_MESENCHYMAL_TRANSITION).

Selected genes: laminins (LAMC2, LAMB3, LAMA3), integrins (ITGA3, ITGA6, ITGAV, ITGB1, ITGB6), other mesenchymal genes (PDPN, FN1, VIM, TGFBI, TNC, PLAUR, SERPINE1, SERPINE2)

Additional comments: abundant in skin (SCC), HNSCC, PDAC and NSCLC. Highly consistent with the EMT program described previously by Puram et al. and Kinker et al.

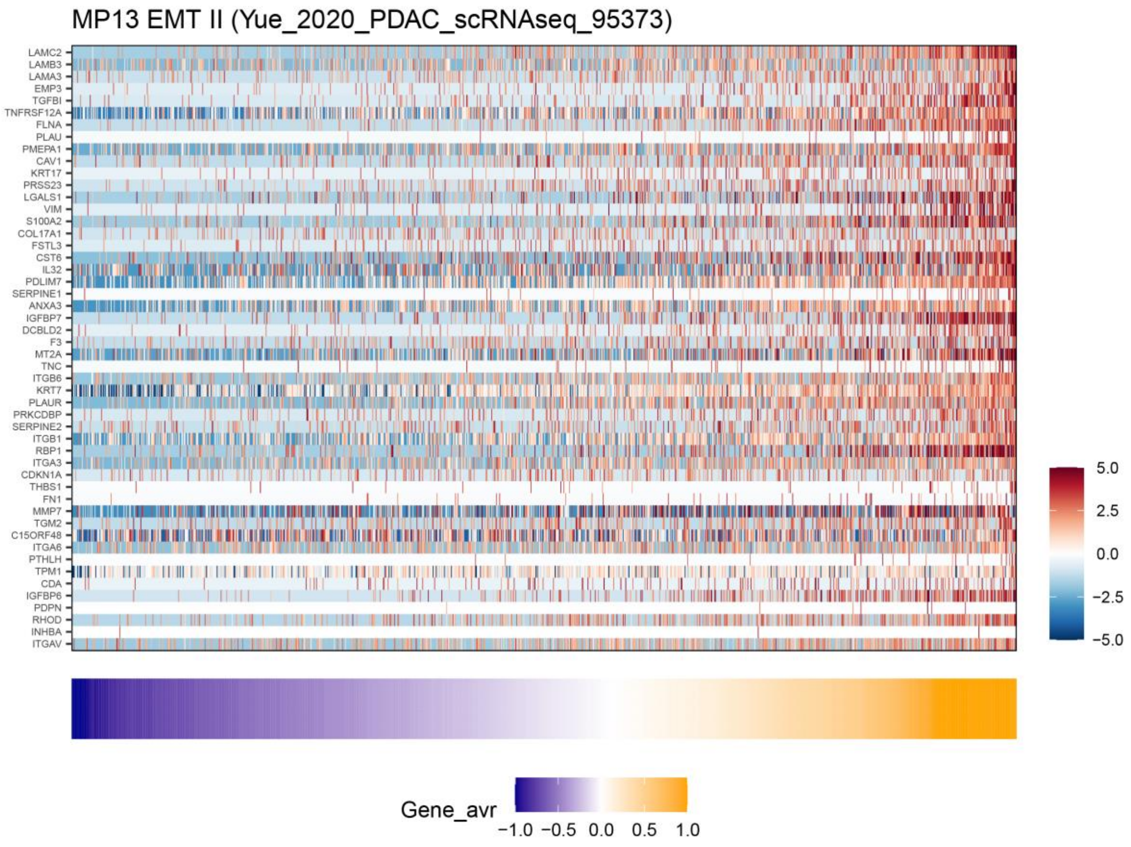

### <u>MP14 – EMT-III</u>

<u>Brief description:</u> *General* MP that belongs to the mesenchymal hallmark. Includes both mesenchymal and epithelial markers, consistent with a hybrid cellular state.

<u>Functional annotation:</u> Enriched with functional annotations related to cadherin binding, muscle and extracellular activity (e.g. C5.GOMF_CELL_CELL_ADHESION_MEDIATOR_ACTIVITY).

<u>Selected genes:</u> cytokeratins (KRT7, KRT8, KRT18, KRT19), Annexins (ANXA1, ANXA2, ANXA3), S100 proteins (S100P, S100A4, S100A6, S100A10, S100A11, S100A14, S100A16), mesenchymal genes (VIM, TAGLN2, MMP7).

<u>Additional comments:</u> A “hybrid” EMT program that includes activation of both mesenchymal and epithelial markers. Abundant in breast, synovial-sarcoma, neuroendocrine tumors, ALL and CTCs.

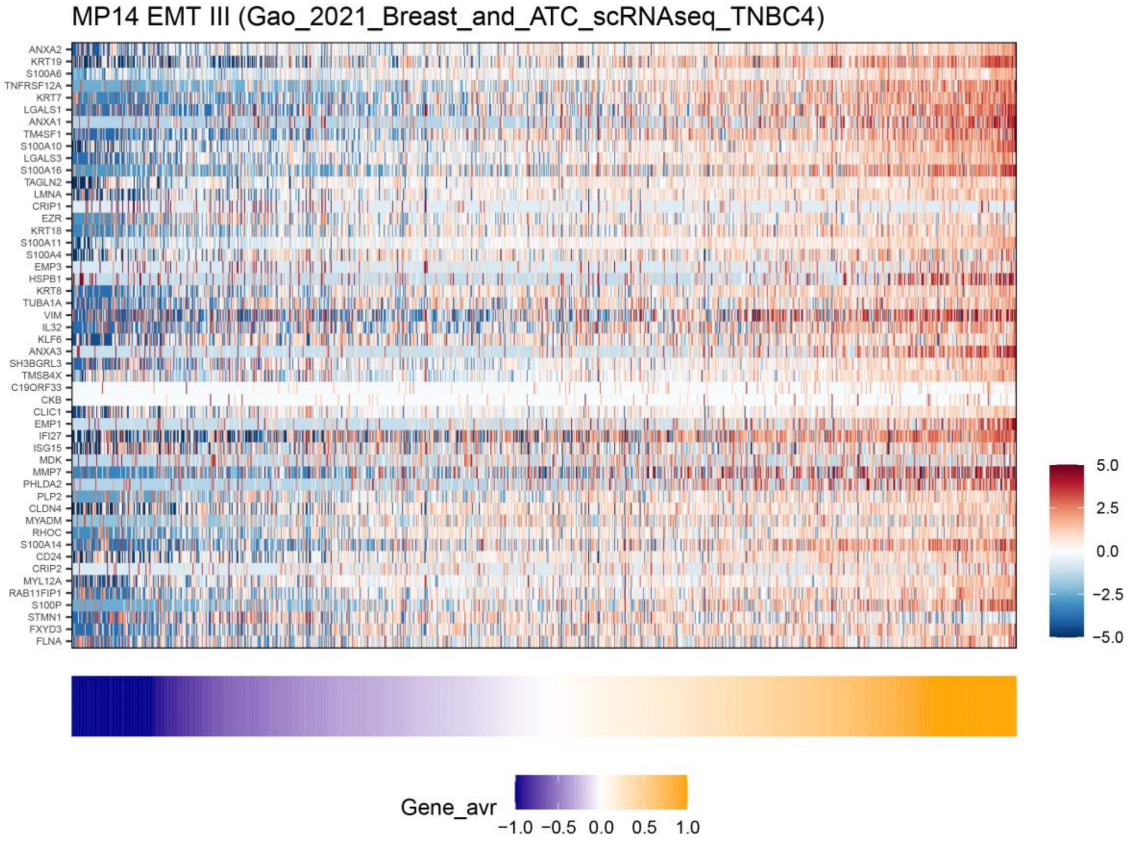

**MP15 – EMT-IV**

Brief description: *Specific* MP that belongs to the mesenchymal hallmark. Includes mesenchymal genes, as well as SOX4 and immune-related genes.

Functional annotation: Enriched with functional annotations related to extracellular matrix (e.g. C5.GOBP_CONNECTIVE_TISSUE_DEVELOPMENT)

Selected genes: Mesenchymal genes (SNAI2, COL17A1, TIMP1, THBS2, DCN, SPARC, CAV1), SOX4, ALDH3A1, NFIB, immune-related genes (CCL2, CXCL14, CD74, IFIT3, IFITM1, C1R, C1S).

Additional comments: Observed in skin (SCC) and HNSCC.

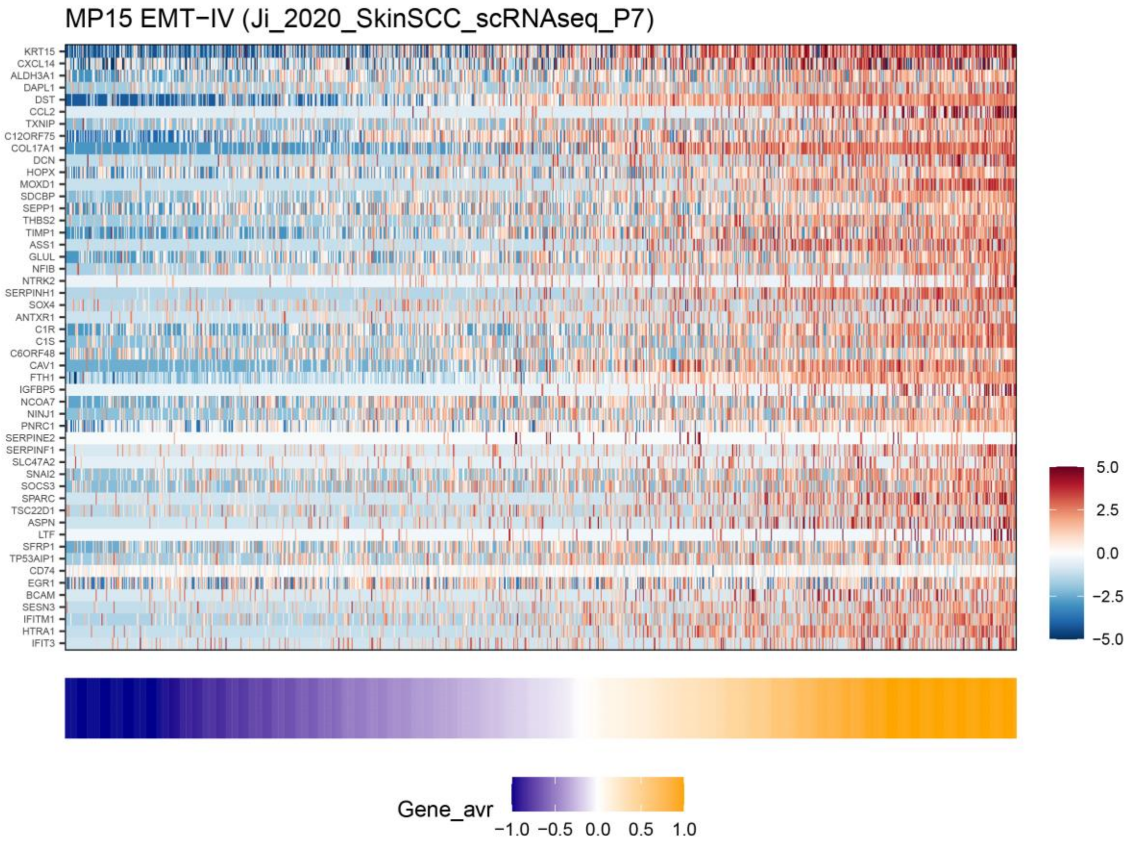

### MP16 – MES (glioma)

Brief description: *Specific* MP that belongs to the mesenchymal hallmark and is specific to glioma.

Functional annotation: Enriched with functional annotations related to EMT (e.g. H.HALLMARK_EPITHELIAL_MESENCHYMAL_TRANSITION).

Selected genes: CD44, ANXA1, ANXA2, WWTR1 (TAZ), TNC

Additional comments: highly abundant in glioma.

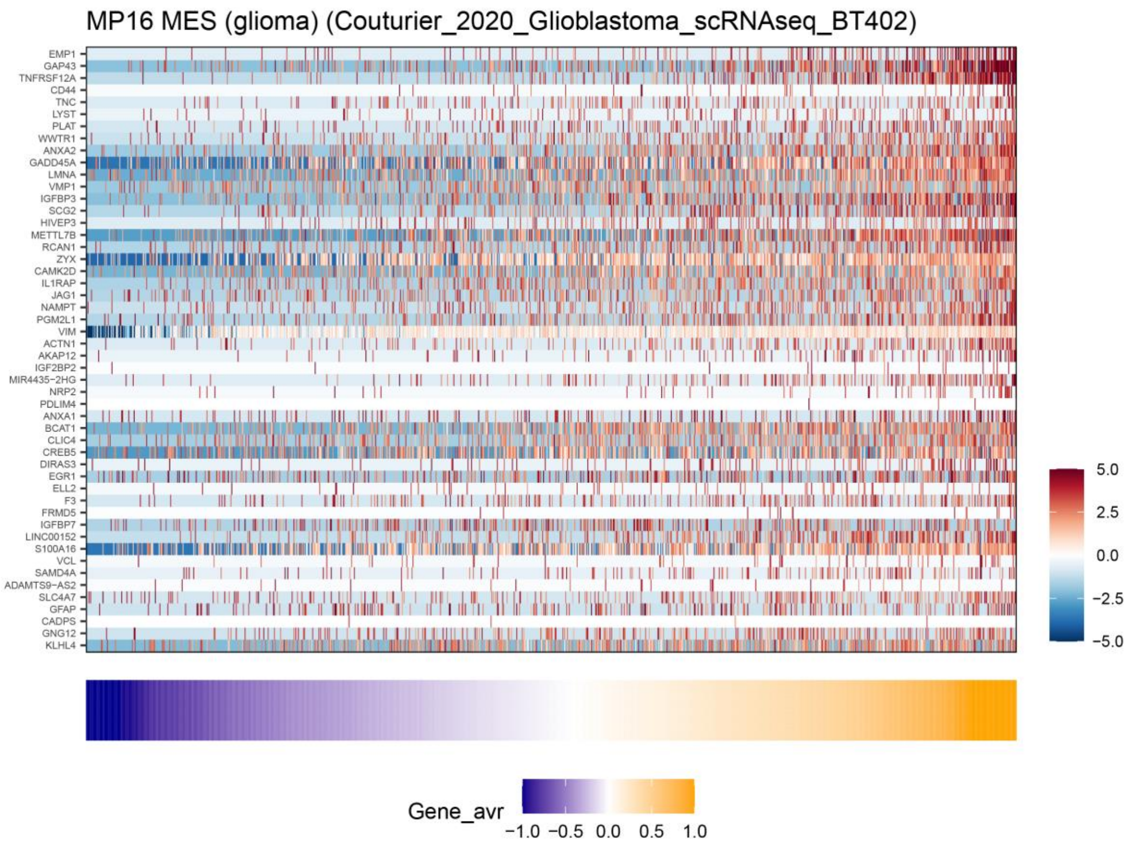

#### <u>MP17 – Interferon/MHC-II (I)</u>

<u>Brief description:</u> *General* MP that belongs to the Interferon/MHC-II hallmark. Compared to the other MP in that hallmark (MP18), this MP consist of more interferon-response and MHC-II genes, also contains MHC-I genes, and less complement genes.

<u>Functional annotations:</u> Enriched with multiple interferon and MHC related annotations (e.g. H.HALLMARK_INTERFERON_ALPHA_RESPONSE and C5.GOCC_MHC_PROTEIN_COMPLEX).

<u>Selected genes:</u> interferon response (ISG15, ISG20, OAS1, MX1, IFI6, IFI27, IFI35, IFI44, IFI44L, IFIT1, IFIT2, IFIT3, IFITM1, IFITM3), MHC-II (CD74, HLA-DMA, HLA-DPB1, HLA-DPA1, HLA-DRA, HLA-DRB1, HLA-DQB1, HLA-DRB5, HLA-DQA1), MHC-I (HLA-A, HLA-B, HLA-C, HLA-E, B2M).

<u>Additional comments:</u> Most abundant in breast cancer, followed by glioma, schwannoma, melanoma, HNSCC, NSCLC, ovarian cancer and cell lines. The interferon and MHC-II components of this MP are highly correlated and hence did not segregate into two distinct MPs.

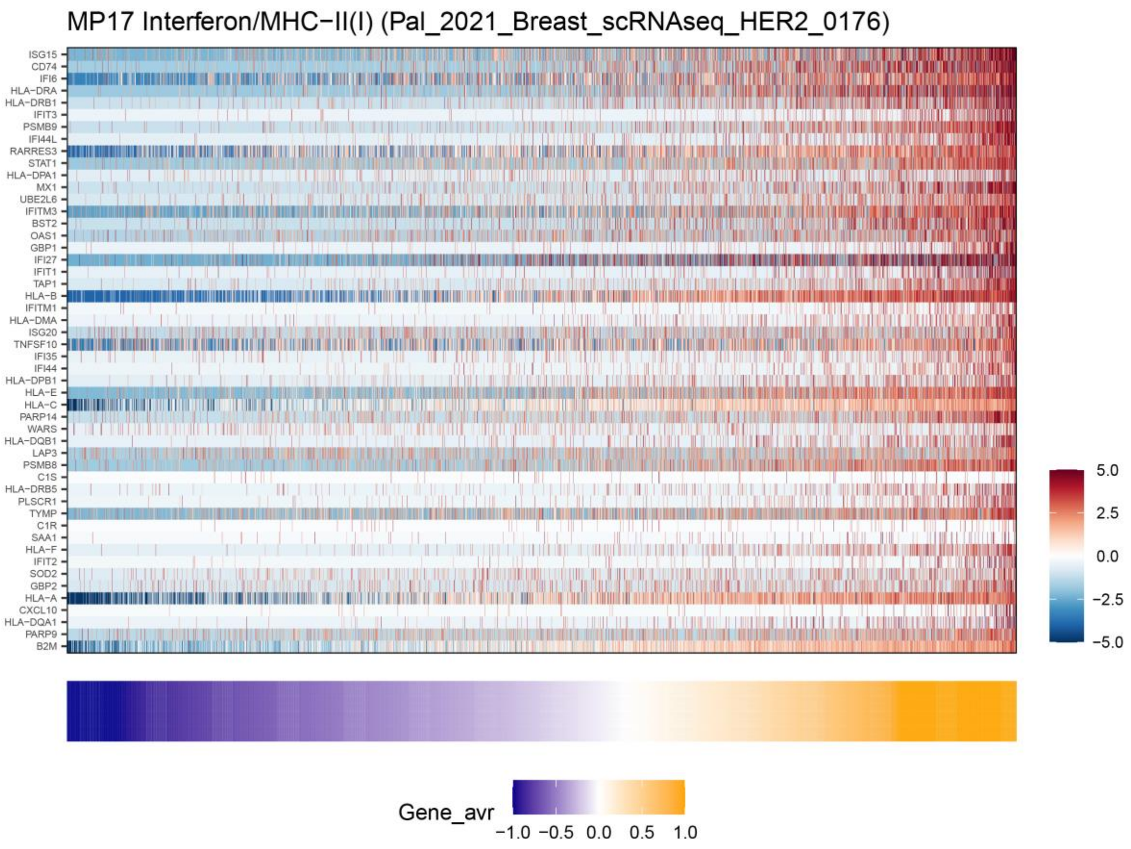

#### MP18 – Interferon/MHC-II (II)

Brief description: *Shared* MP that belongs to the Interferon/MHC-II hallmark. Compared to the other MP in that hallmark (MP17), this MP includes more complement genes, does not contain MHC-I genes and contain fewer interferon-response and MHC-II genes.

Functional annotations: Enriched with interferon and MHC related annotations (e.g. H.HALLMARK_INTERFERON_GAMMA_RESPONSE and C5.GOMF_MHC_CLASS_II_PROTEIN_COMPLEX_BINDING).

Selected genes: MHC-II (CD74, HLA-DMA, HLA-DRA, HLA-DRB1, HLA-DPA1, HLA-DRB5), interferon response (IFI6, IFITM2, IFITM3, ISG15), complement (C1S, C1R, C3, CLU, SERPING1).

Additional comments: abundant in schwannoma, followed by ovarian cancer. The interferon and MHC-II components of this MP are highly correlated and hence did not segregate into two distinct MPs.

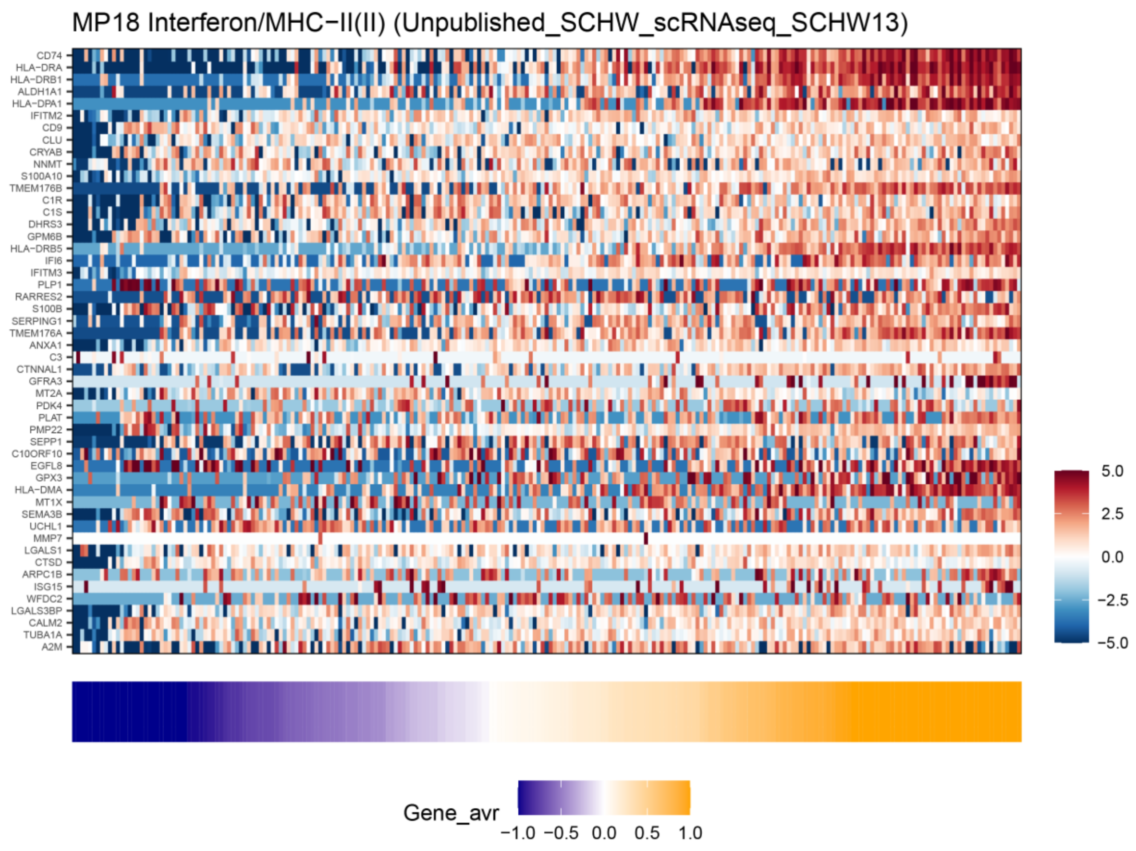

#### MP19 – EpiSen (epithelial senescence)

Brief description: *Shared* MP that constitutes the senescence hallmark. This program was named epithelial senescence because it highly resembles the expression response of keratinocytes and other epithelial cells when entering senescence, and because it correlates with decreased cell cycle (see Kinker et al. for further details). It includes many secreted factors (consistent with the senescence-associated secretory phenotype, or SASP) and many epithelial genes (potentially indicative of a connection between senescence and epithelial differentiation).

Functional annotations: Enriched with epithelial related annotations (e.g. C5.GOCC_KERATIN_FILAMENT) and highly resembles the EpiSen signature previously described by us (Kinker et al. 2020).

Selected genes: S100A8, S100A9, S100A7, S100P, SPRR1B, SPRR2D, SPRR3, SLPI, LCN2, PI3, SAA1, AQP3, CLDN4, LY6D, KRT16, KRT17, KRT6B, KRT6A, SERPINB1, SERPINB3, SERPINB4.

Additional comments: Enriched in multiple types of squamous cell carcinomas (SCC) but also found in other cancer types. Specifically, most abundant in HNSCC followed by skin SCC, NSCLC, PDAC and cell lines. Negatively correlated with cell cycle.

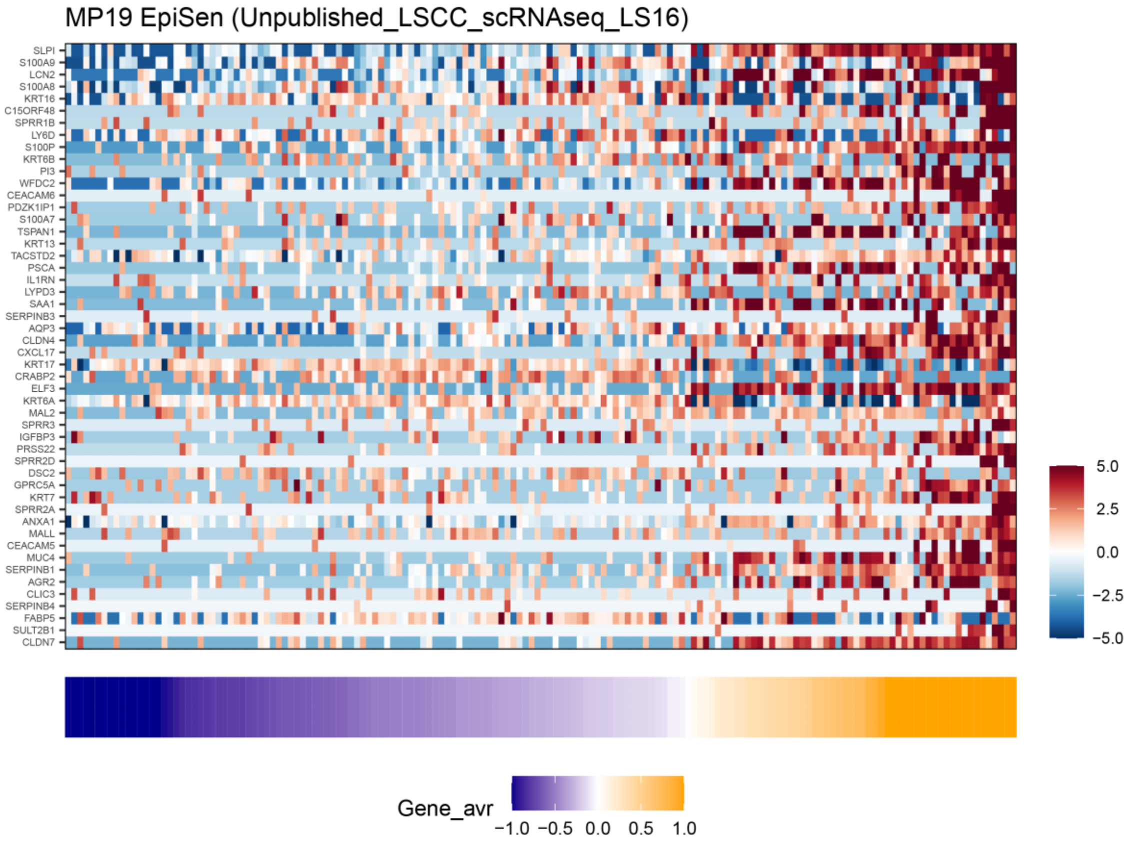

### MP20 – MYC-targets

Brief description: *General* MP that constitutes the oncogenic hallmark and consists of MYC and many of its target genes.

Functional annotations: Enriched with MYC related annotations (e.g. H.HALLMARK_MYC_TARGETS_V1).

Selected genes: MYC, NOP56, NOP16, NOLC1, MRTO4, PHB, PA2G4, HSPE1, DCTPP1, SRM, GNL3, HSPD1, CCT2, ODC1, CCT5, RAN

Additional comments: Most abundant in AML and ALL, followed by skin SCC.

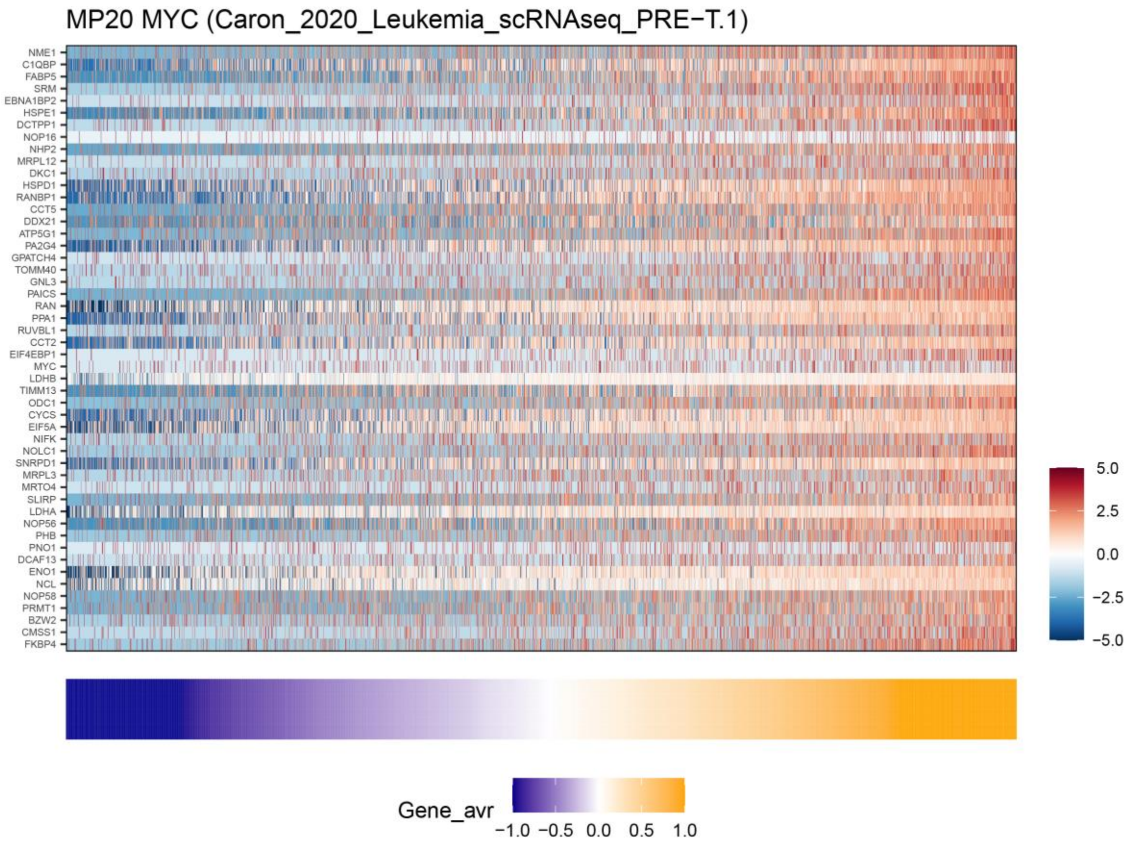

**<u>MP21 – Respiration</u>**

<u>Brief description:</u> *Shared* MP that constitutes the respiration hallmark.

<u>Functional annotations:</u> Enriched with respiration related annotations (e.g. C5.GOCC_RESPIRATORY_CHAIN_COMPLEX).

<u>Selected genes:</u> ATP-synthase (ATP5E, ATP5I, ATP5G1, ATP5J2), NADH Dehydrogenase (NDUFA1, NDUFA3, NDUFA7, NDUFA12, NDUFAB1, NDUFB1, NDUFB2, NDUFB4, NDUFB8, NDUFC1, NDUFS5, NDUFS6), Cytochrome c oxidase (COA3, COX6B1, COX7B, COX17).

<u>Additional comments:</u> Most abundant in glioma followed by medulloblastoma, melanoma, AML and CML.

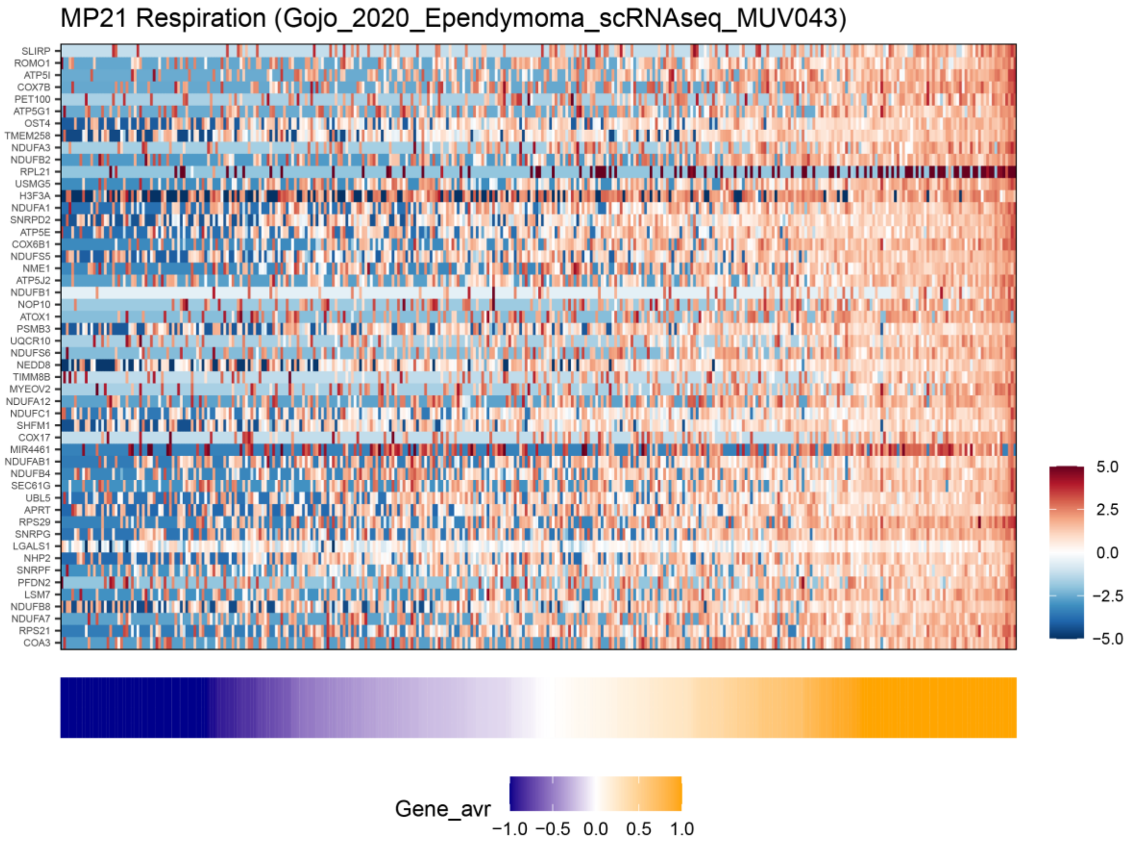

### <u>MP22 – Secreted I</u>

<u>Brief description:</u> *Shared* MP that belongs to the secreted hallmark.

<u>Functional annotations:</u> Enriched with annotations related to secreted compounds (e.g. C5.GOMF_CYTOKINE_ACTIVITY).

<u>Selected genes:</u> cytokine and chemokines (CCL2, CCL20, CXCL1, CXCL2, CXCL8, LIF), EGFR ligands (HBEGF, AREG).

<u>Additional comments:</u> Most abundant in colorectal and renal cancer, followed by prostate cancer.

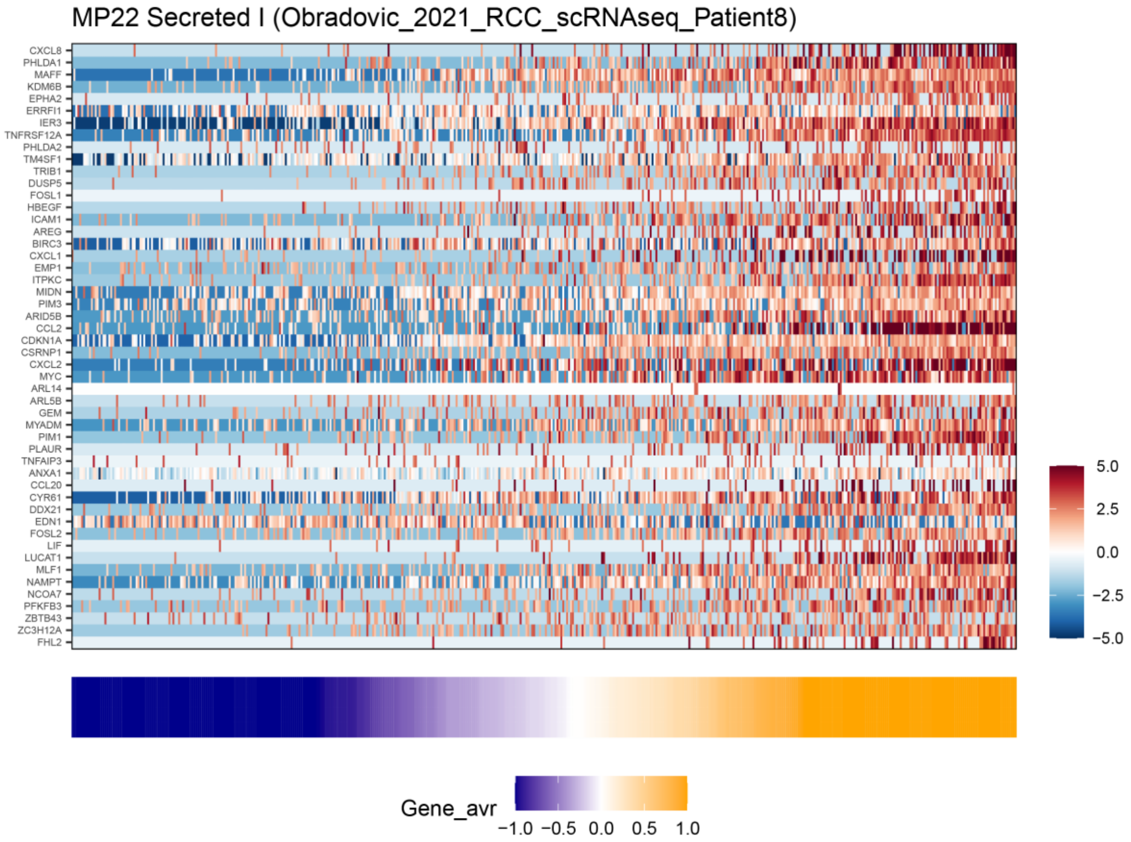

#### MP23 – Secreted II

Brief description: *Shared* MP that belongs to the secreted hallmark.

Functional annotations: Enriched with annotations related to secreted compounds (e.g. H.HALLMARK_COMPLEMENT).

Selected genes: EpiSen-associated secreted proteins (LCN2, SLPI, S100A9, SAA1, SAA2), cytokine and chemokines (MDK, CXCL1, CXCL2, CXCL8), complement (C3, CFB, CLU), MMP7, MUC5B, SERPINF1.

Additional comments: highly abundant in NSCLC. Contains many of the secreted factors seen in EpiSen (MP19), potentially indicating a connection between the two meta-programs.

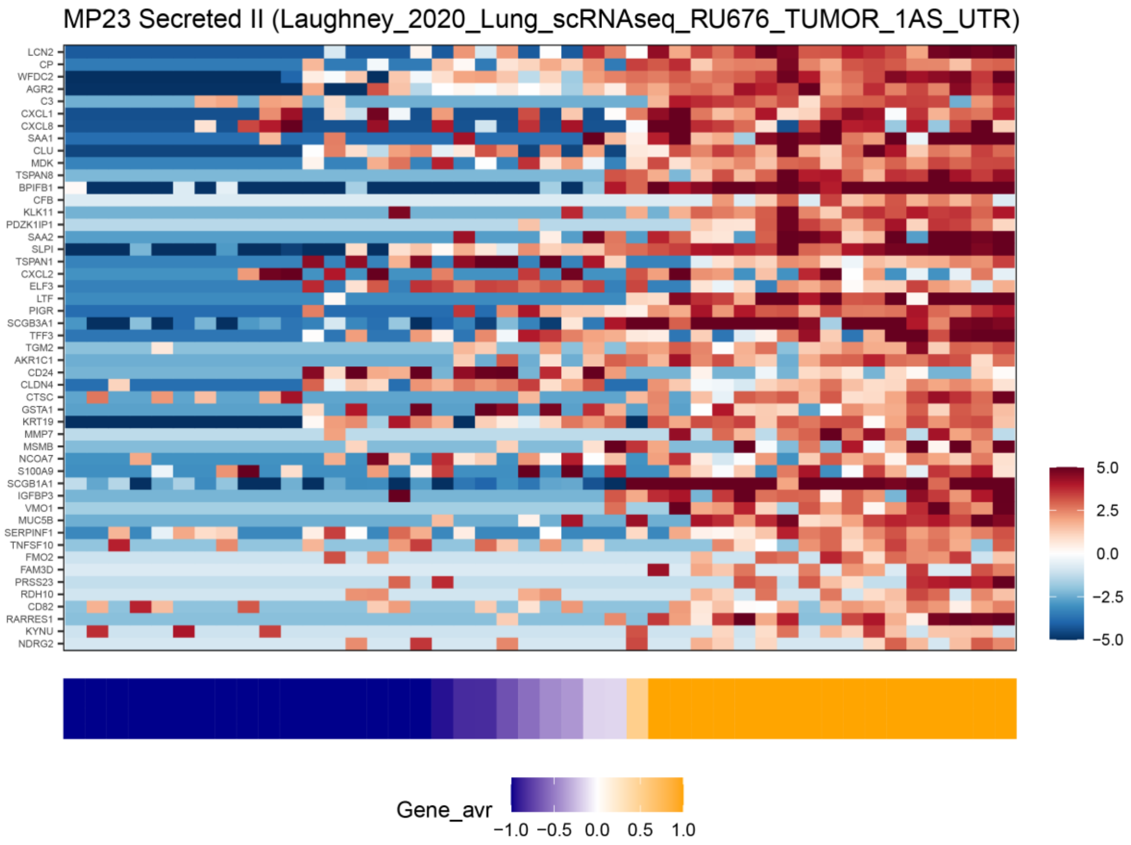

#### <u>MP24 – Cilia</u>

<u>Brief description:</u> *Shared* MP that constitutes the cilia hallmark.

<u>Functional annotations:</u> Enriched with multiple annotations related to cilia (e.g. C5.GOBP_CILIUM_ORGANIZATION).

<u>Selected genes:</u> DNAAF1, DNAH5, ZMYND10, RSPH1, EFHC1, IFT57, IQCG, SPA17

<u>Additional comments:</u> Highly abundant in NSCLC. Also highly abundant in ependymoma, consistent with cilia as a known characteristic of ependymal cells; our main analysis grouped together ependymoma and other gliomas (that do not contain this MP), and therefore it was not identified as abundant in glioma (see Fig. 3A), but when separating glioma to different types we find a highly significant enrichment in ependymoma (not shown). Previously described in the context of ependymoma (Gojo et al. 2020), where it was named “ependymal-like” due to the known association of cilia with ependymal cells.

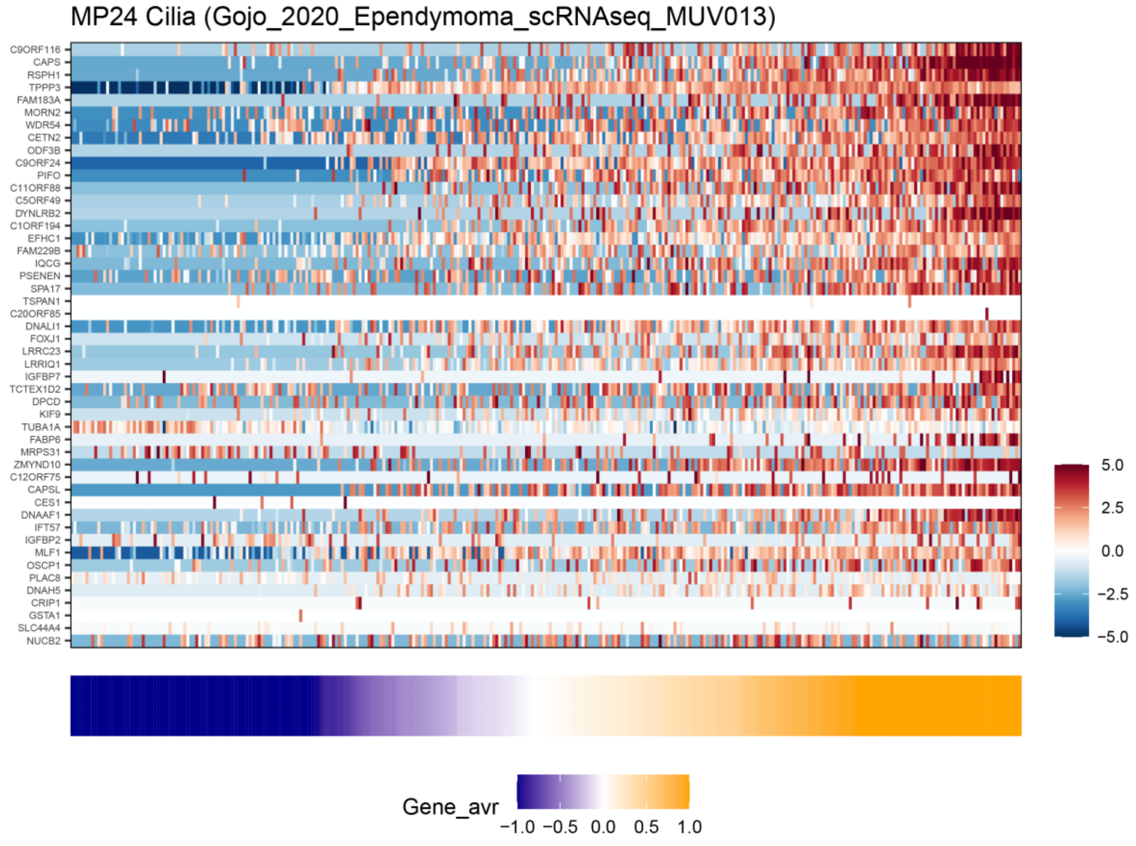

#### MP25 – Astrocytes

Brief description: *Shared* MP that belongs to the lineage-related (neural) hallmark.

Functional annotations: Enriched with multiple annotations related to astrocytes (e.g. C8.ZHONG_PFC_MAJOR_TYPES_ASTROCYTES).

Selected genes: GFAP, CLU, AGT, SLC1A3, HEPN1, AQP4, APOE, SPARC, MLC1

Additional comments: Highly abundant in glioma, in which it was previously described by multiple previous studies and it defines one of four major cellular states (see Neftel et al. 2019)

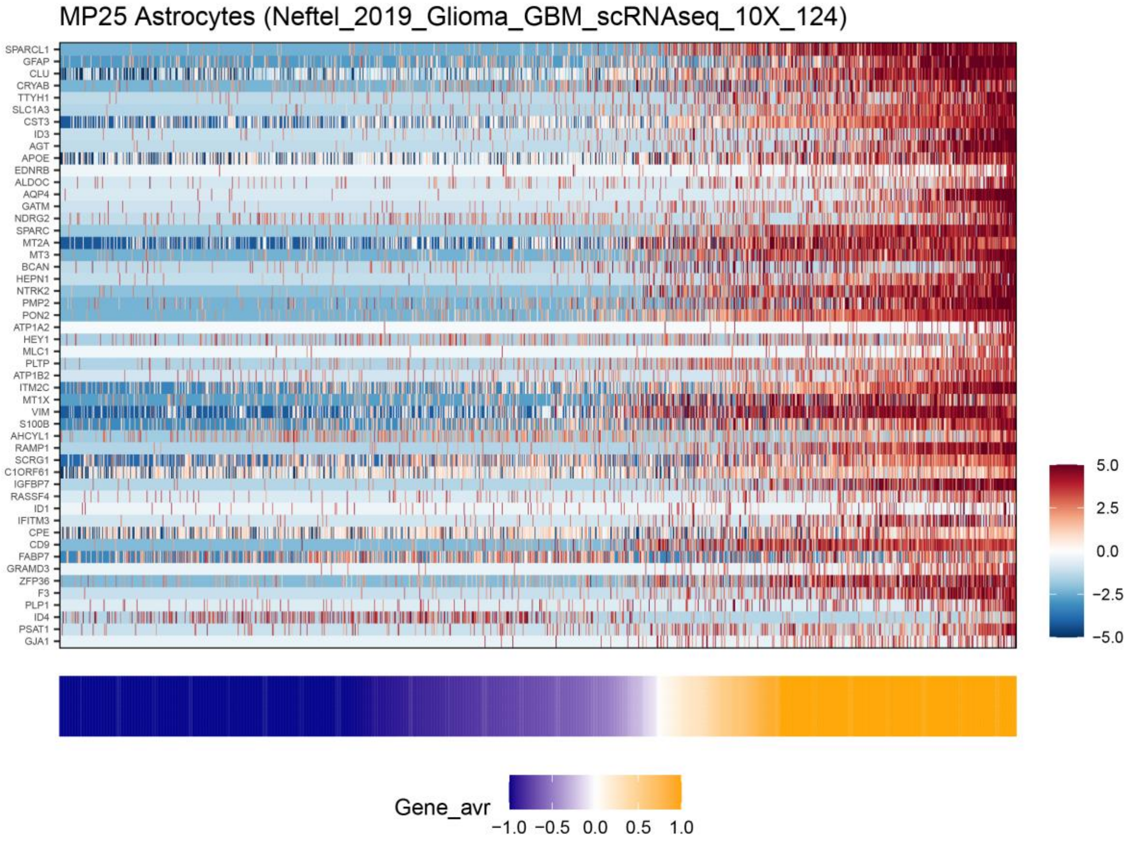

**<u>MP26 – NPC (glioma)</u>**

<u>Brief description:</u> *Shared* MP that belongs to the lineage-related neural hallmark.

<u>Functional annotations:</u> Enriched with annotations related to NPC (e.g. idh_mutant_glioma_sigs.IDH_A_NPC).

<u>Selected genes:</u> DCX, CD24, SOX11, SOX4, STMN1, STMN2, BEX2, DLX5

<u>Additional comments:</u> Abundant in glioma and medulloblastoma. Was previously described by multiple previous glioma studies in which it defines one of four major cellular states (see Neftel et al. 2019).

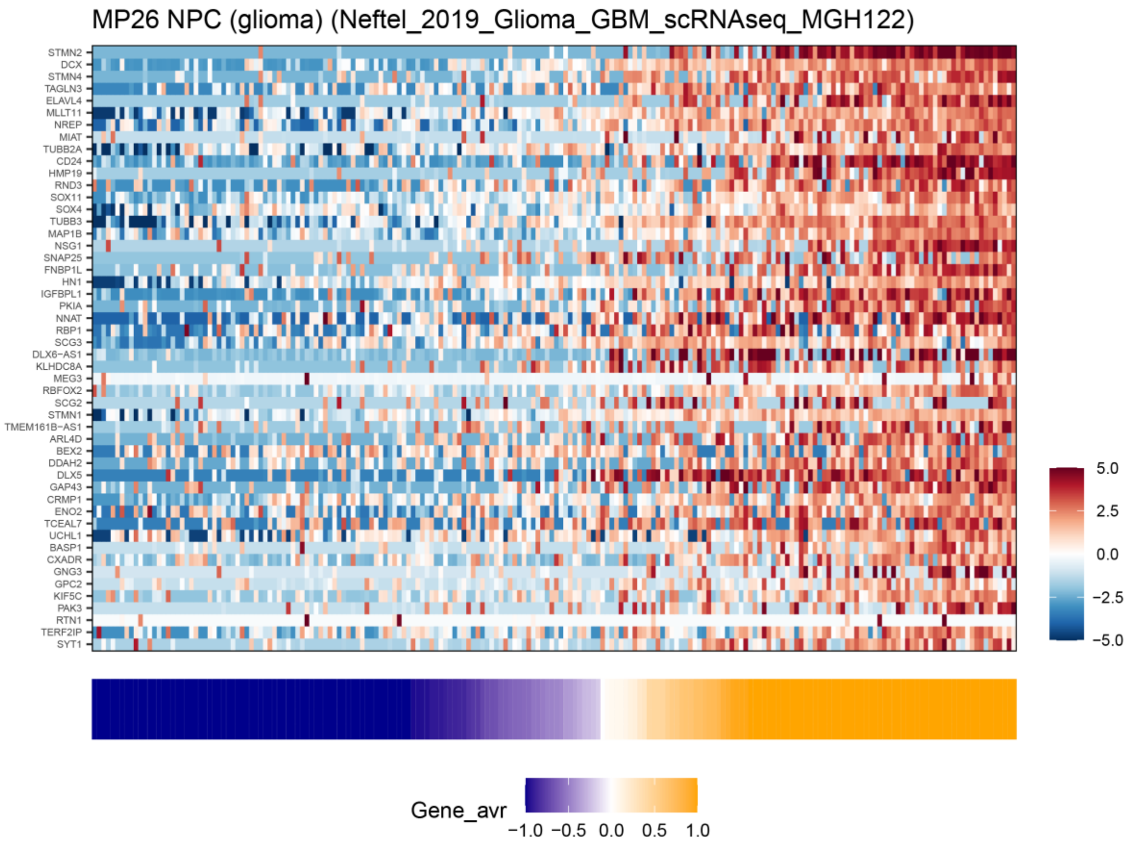

**<u>MP27 – Oligo. progenitor</u>**

<u>Brief description:</u> *Specific* MP that belongs to the lineage-related neural hallmark.

<u>Functional annotations:</u> Enriched with annotations related to oligodendrocyte differentiation (e.g. C5.GOBP_GLIAL_CELL_DIFFERENTIATION).

<u>Selected genes:</u> OLIG1, OLIG2, PDGFRA, NEU4, TNR, LHFPL3 ASCL1,

<u>Additional comments:</u> Abundant in glioma. Was previously described by multiple previous glioma studies in which it defines one of four major cellular states (see Neftel et al. 2019).

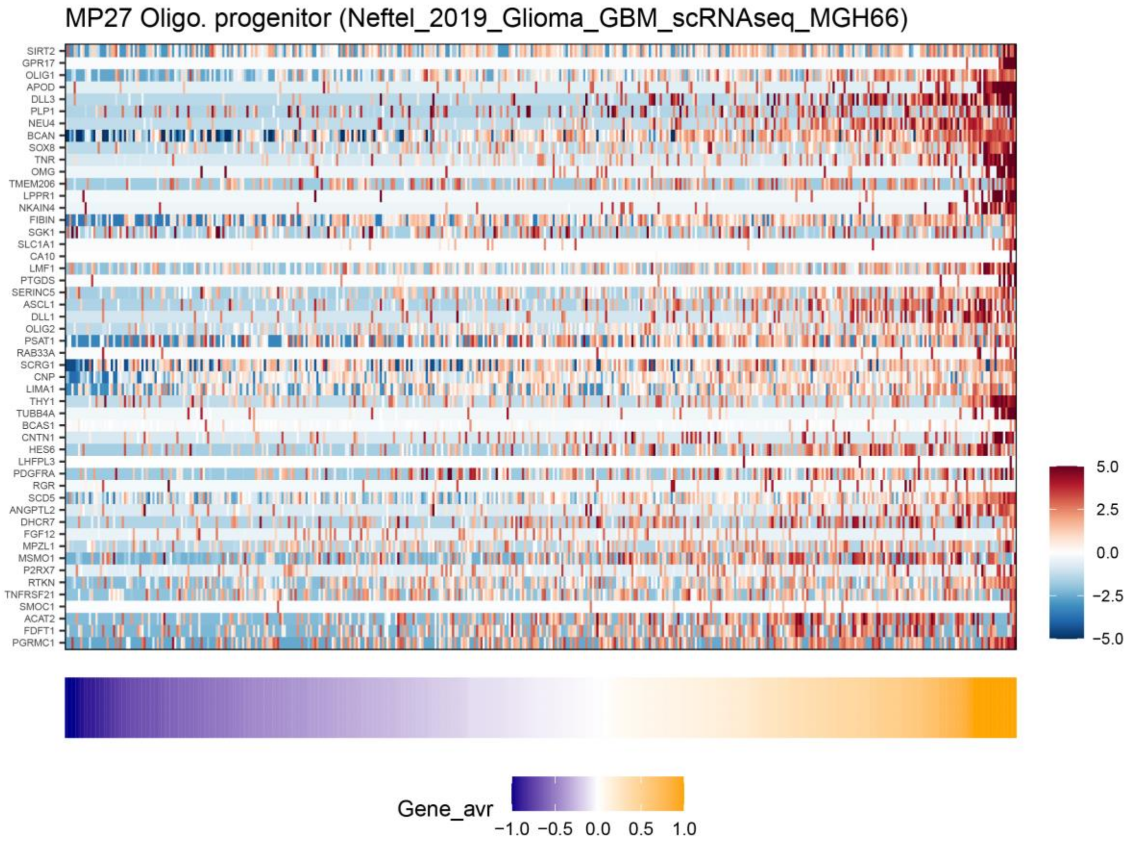

#### MP28 – Oligo. normal

Brief description: *Specific* MP that belongs to the lineage-related neural hallmark.

Functional annotations: Enriched with annotations related to oligodendrocytes (e.g. C5.GOCC_MYELIN_SHEATH).

Selected genes: MAG, MOBP, MAL, PLP1, CLDN11, UGT8, SLC44A1, CMTM5.

Additional comments: Abundant in glioma. Malignant glioma cells typically resemble oligodendrocyte progenitor cells (OPC; MP27), but some malignant cells resemble more differentiated oligodendrocytes, as defined by this program. This program is also identified in the normal (i.e. non-malignant) glial cells within glioma tumors.

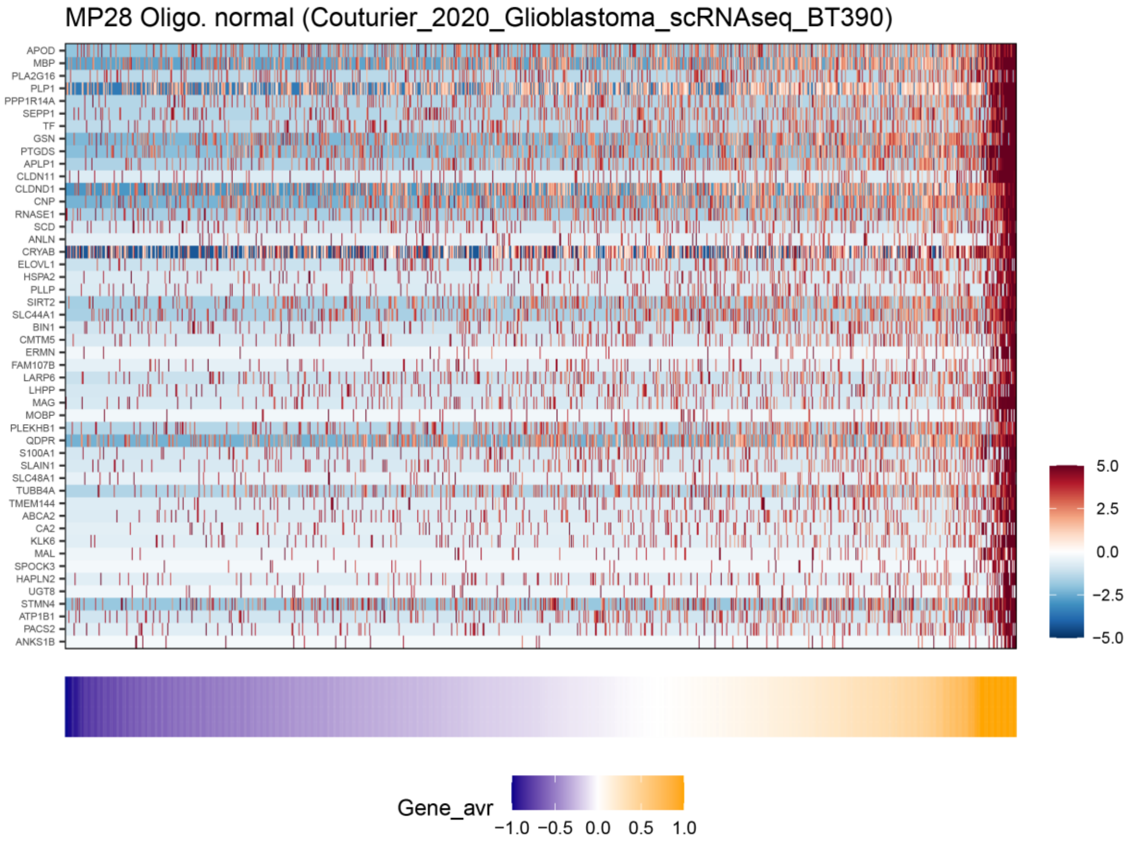

#### MP29 – NPC/OPC

Brief description: *Specific* MP that belongs to the lineage-related neural hallmark. Includes both OPC and NPC markers and hence more difficult to resolve as a specific type of progenitor. Found primarily in glioma cell lines, while glioma tumors primarily contain the more specific programs (either OPC-like or NPC-like, i.e. MP26-27).

Functional annotations: Enriched with annotations related to both neuroal and oligodendrocyte progenitors (NPC or OPC, e.g. BrainSignatures2.OPC.0-poli19).

Selected genes: OPC markers (OLIG1, OLIG2, CHD7), NPC and neuronal markers (DCX, SOX4, SOX5, SOX6, SOX11, DLL3, DCX, MAP2).

Additional comments: Most abundant in glioma cell lines

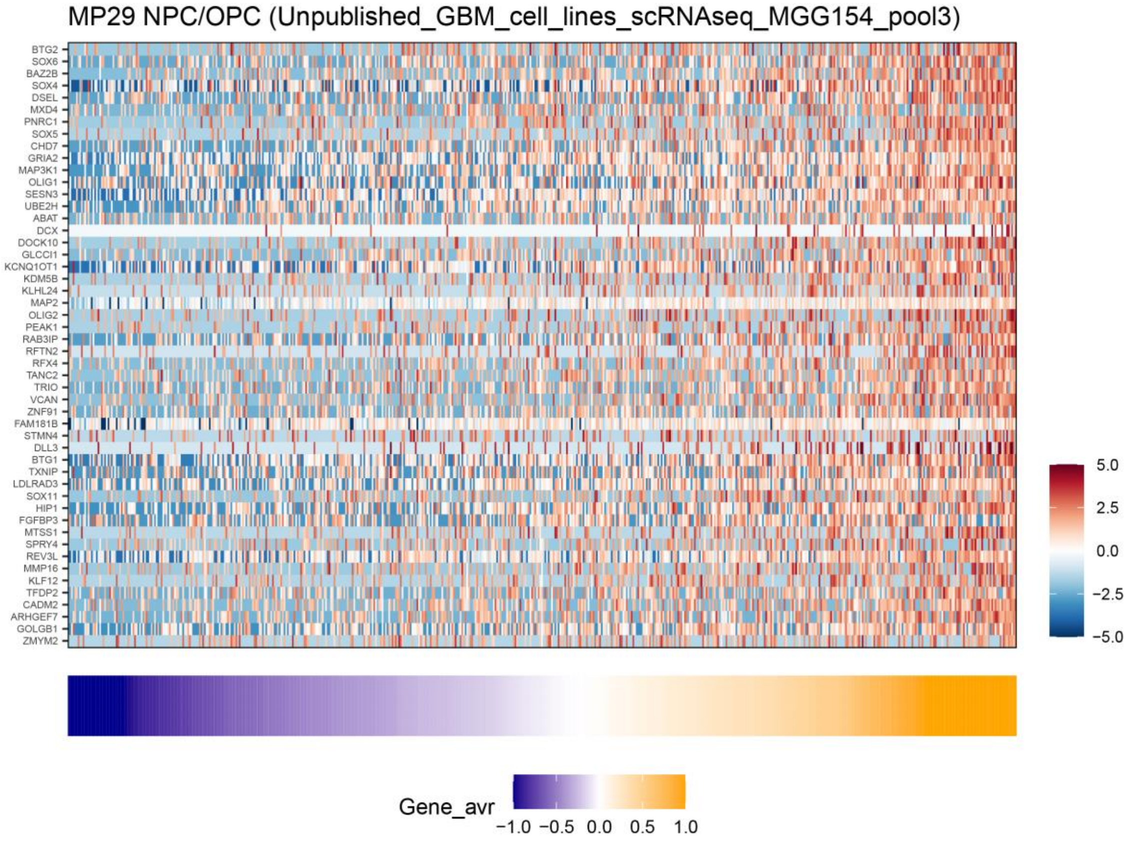

#### MP30 – PDAC-classical

Brief description: *Shared* MP that belongs to the lineage-related hallmark. Highly resembles the signatures of the classical subtype of PDAC, and accordingly detected primarily in PDAC, but also in some non-PDAC tumors.

Functional annotations: Enriched with annotations related to the PDAC classical program (e.g. PDAC.Yue_PDAC_Classical published by Yue et al. 2020).

Selected genes: TFF1, TFF2, TFF3, CEACAM6, AGR2, AGR3, LGALS4, CTSE, MUC13, MUC5AC, MUC5B.

Additional comments: Most abundant in PDAC followed by colorectal cancer.

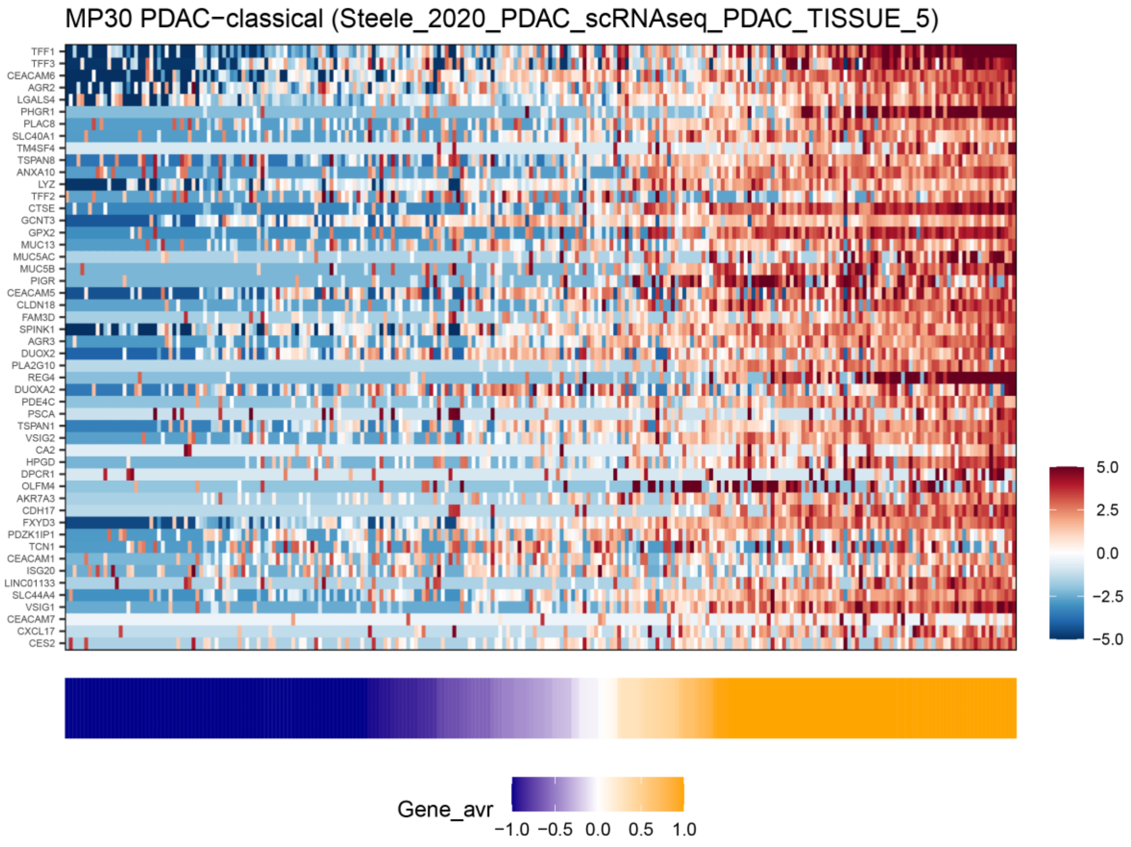

#### MP31 – Alveolar

Brief description: *Specific* MP that belongs to the lineage-related hallmark.

Functional annotations: Enriched with multiple annotations related to alveolar cells (e.g. C8.TRAVAGLINI_LUNG_ALVEOLAR_EPITHELIAL_TYPE_2_CELL).

Selected genes: surfactant protein genes (SFTPA1, SFTPA2, SFTPD, SFTPC, SFTPB, SFTA2, SFTA3, SFTA1P).

Additional comments: highly abundant in NSCLC

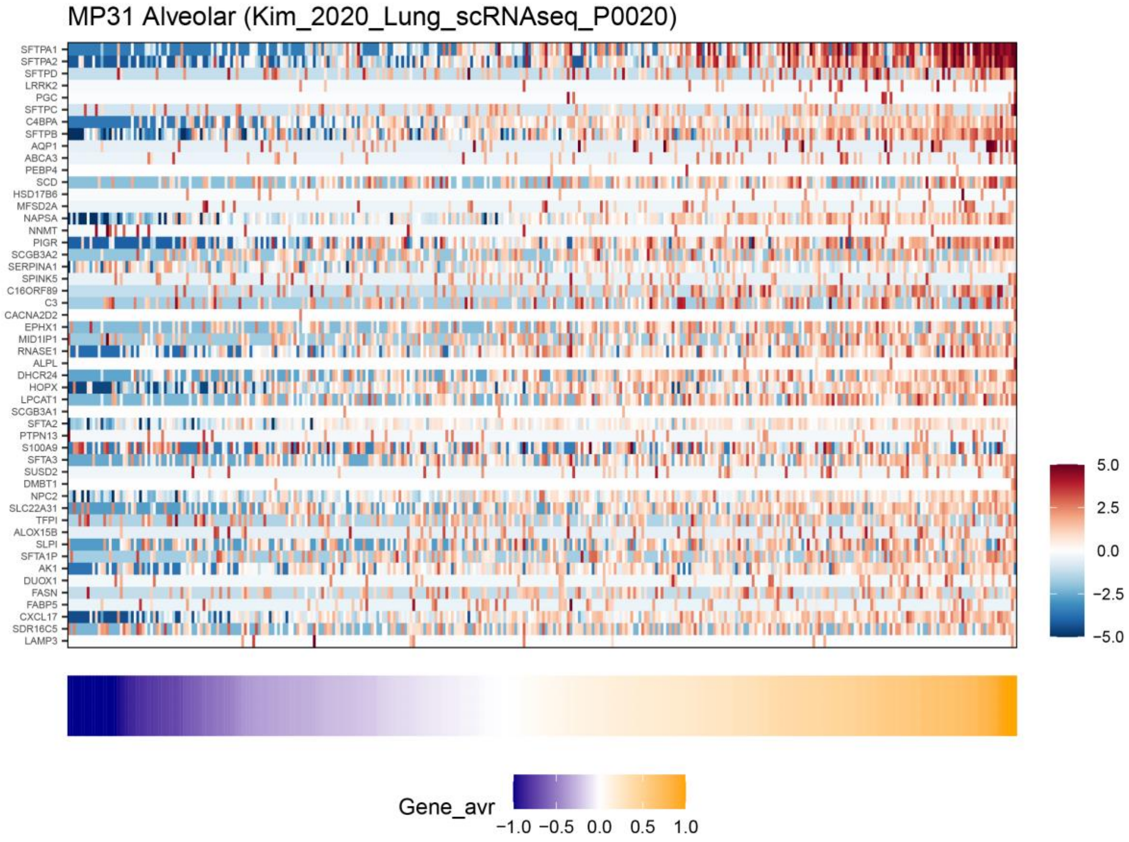

**MP32 – Skin pigmentation**

Brief description: *Specific* MP which belongs to the lineage-related hallmark.

Functional annotations: Enriched with multiple annotations related to pigmentation (e.g. C5.GOBP_PIGMENTATION)

Selected genes: MITF, MLANA, PMEL, TYRP1, GPNMB, DCT.

Additional comments: highly abundant in skin cell lines. Previously described in various melanoma studies, often termed as an MITF-driven program.

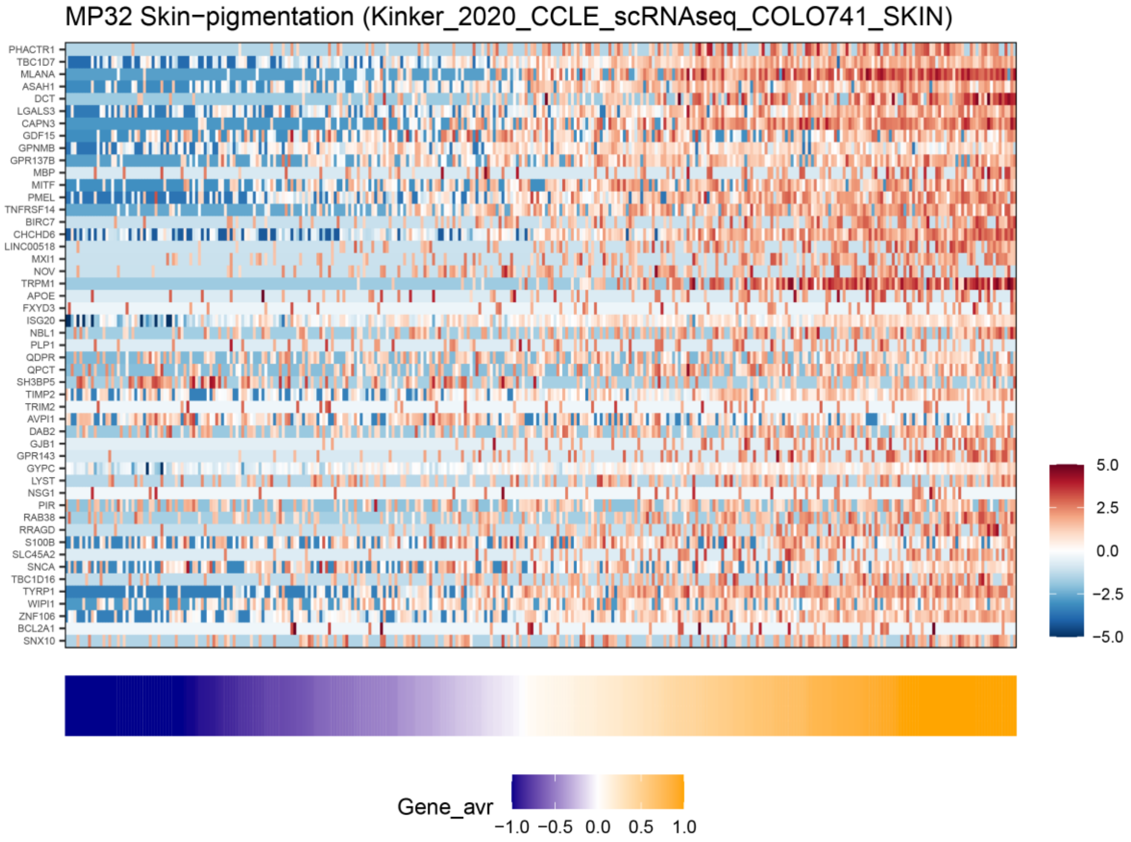

#### MP33 – RBCs

Brief description: *Shared* MP that belongs to the lineage-related (hematological) hallmark, includes many hemoglobin and other red blood cell (RBC) genes.

Functional annotations: Enriched with multiple annotations related to erythrocytes (e.g. H.HALLMARK_HEME_METABOLISM)

Selected genes: hemoglobin genes (HBB, HBG2, HBD, HBA1, HBA2, HBQ1), GYPB, ANK1, GATA1, AHSP

Additional comments: highly abundant in CML, followed by myeloproliferative cancers and CTCs.

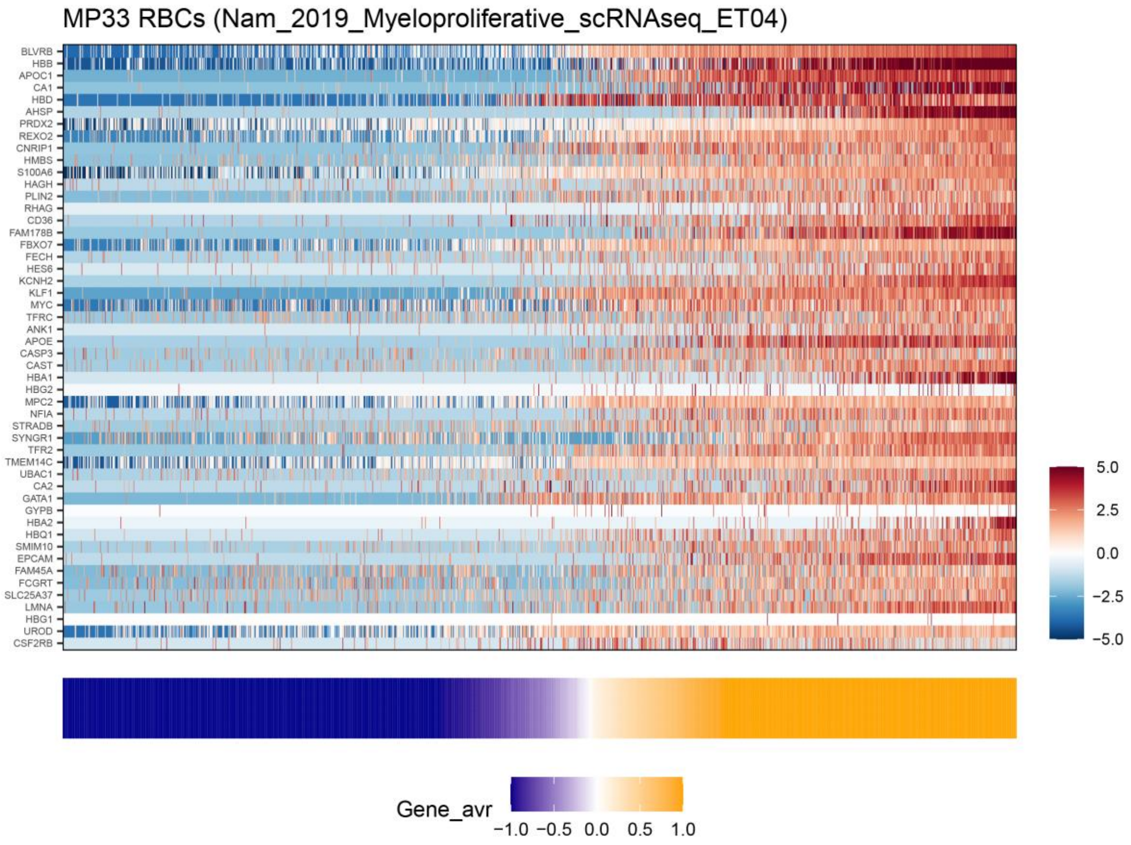

#### MP34 – Platelet-activation

Brief description: *Shared* MP that belongs to the lineage related (hematological) hallmark.

Functional annotations: Enriched with multiple annotations related to platelets (e.g. C5.GOBP_PLATELET_DEGRANULATION).

Selected genes: PLEK, PF4, VASP, RAP1B, GP9, FCER1G, FERMT3.

Additional comments: Most abundant in myeloproliferative cancer and CTCs.

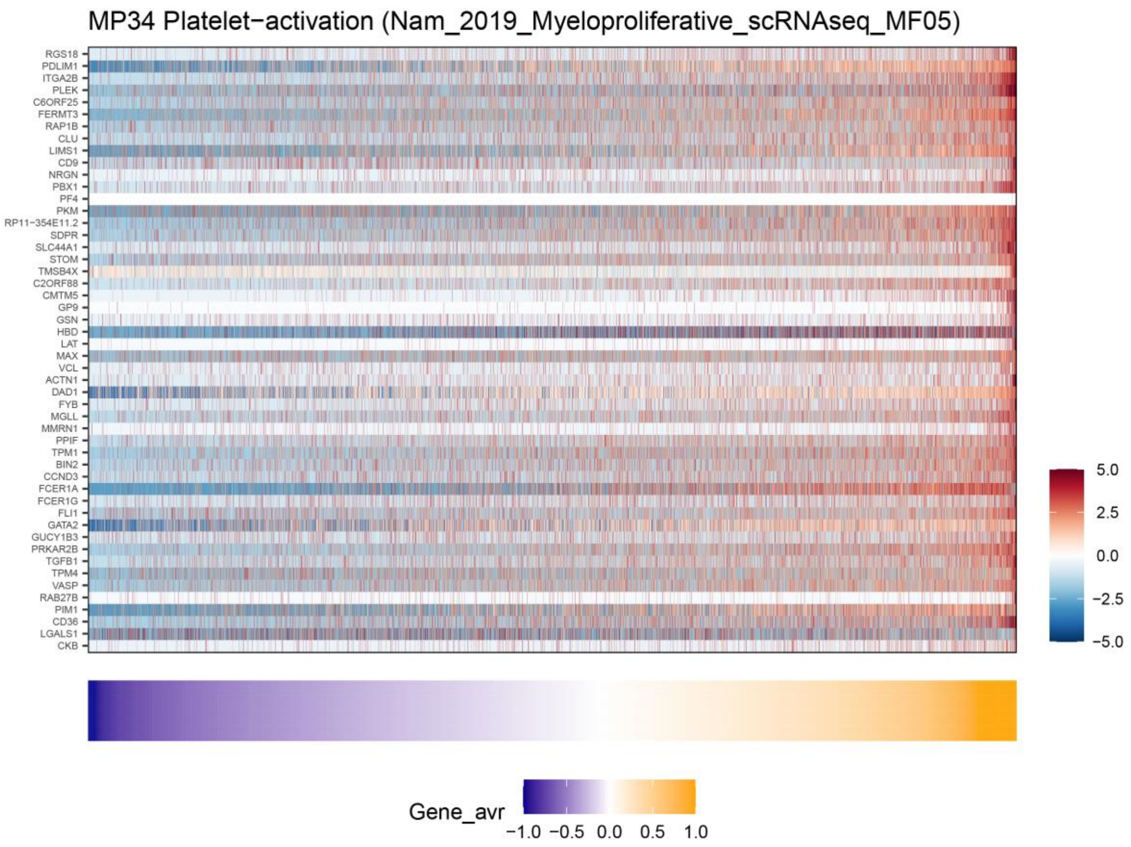

#### MP35 – Hemato. related I

Brief description: *Specific* MP that belongs to the lineage related (hematological) hallmark.

Functional annotations: Enriched with annotations related to the hematological lineage, which we could not map into any specific function.

Selected genes: SPINK2, MS4A3, PRSS57, PRTN3, AZU1, MPO, CD52, SELL, PTPRCAP, IGLL1

Additional comments: Most abundant in myeloproliferative cancers followed by AML.

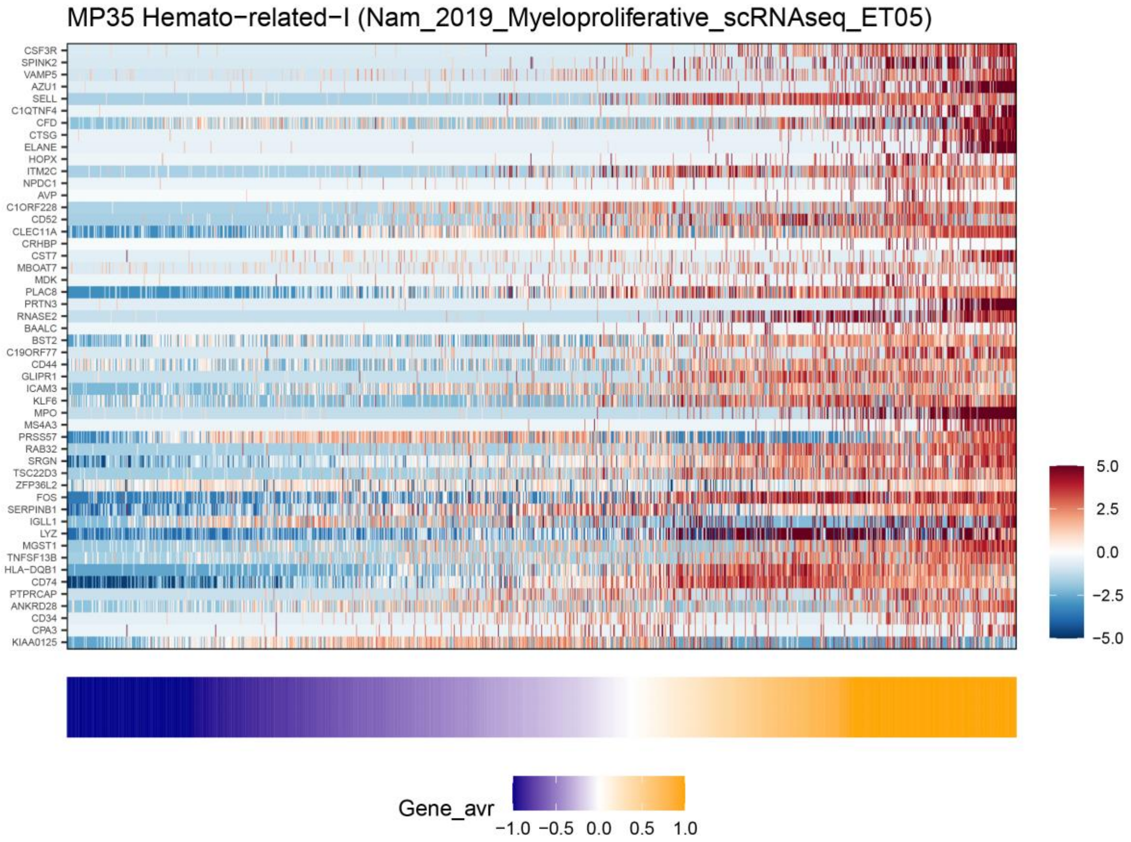

#### MP36 – immunoglobulins (IGs)

Brief description: *Specific* MP that belongs to the lineage related (hematological) hallmark and includes many immunoglobulins.

Functional annotations: Enriched with multiple annotations related to IGs (e.g. C5.GOCC_IMMUNOGLOBULIN_COMPLEX)

Selected genes: IGLC2, IGLC3, IGHA2, IGHA1, IGHG4, IGKC, IGHG1, IGHG3, IGHG2, IGHGP, IGLJ2

Additional comments: Most abundant in multiple myeloma followed by CTCs.

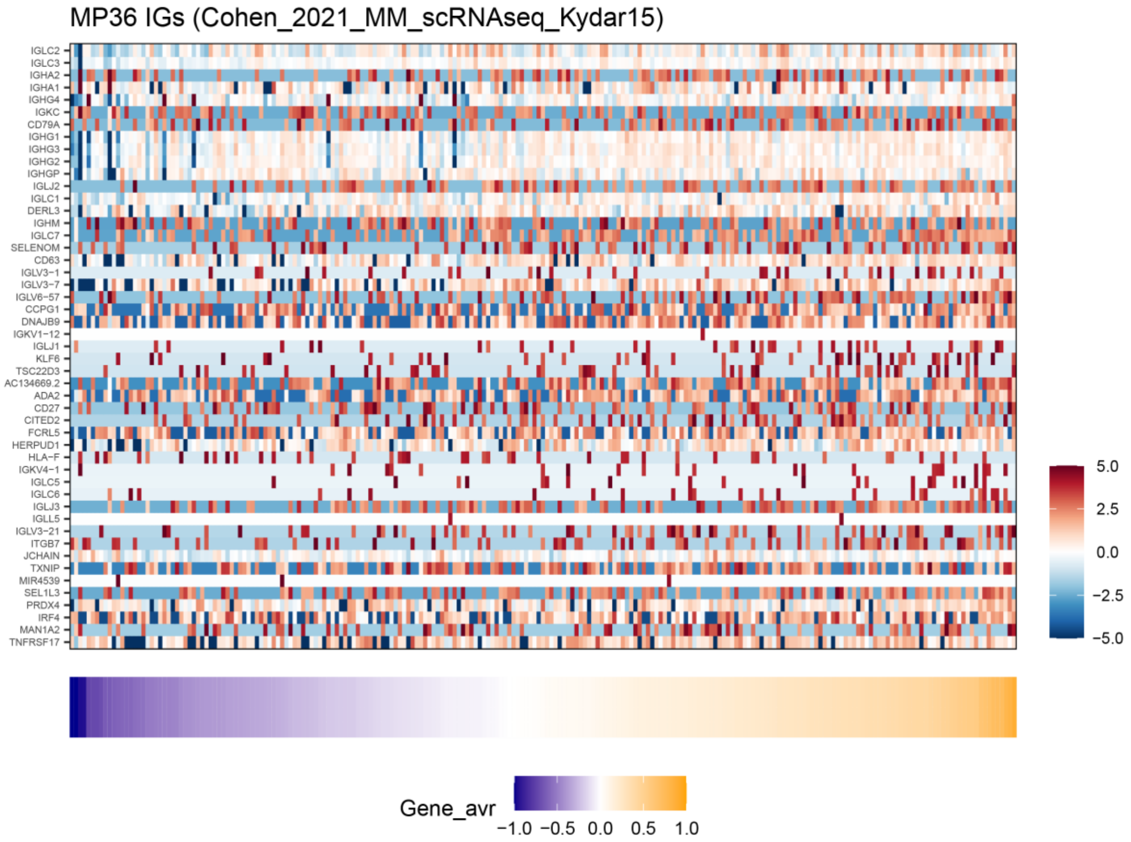

#### MP37 – Hemato. related II

Brief description: *Specific* MP that belongs to the lineage related (hematological) hallmark.

Functional annotations: Enriched with annotations related to hematological lineage, which we could not map into any specific function

Selected genes: NFE2, IL1B, GATA2, KLF1, CPA3, FCER1A, RGS18, MPP1, CD82, CD84, PLEK, CNRIP1

Additional comments: significantly abundant in myeloproliferative cancers.

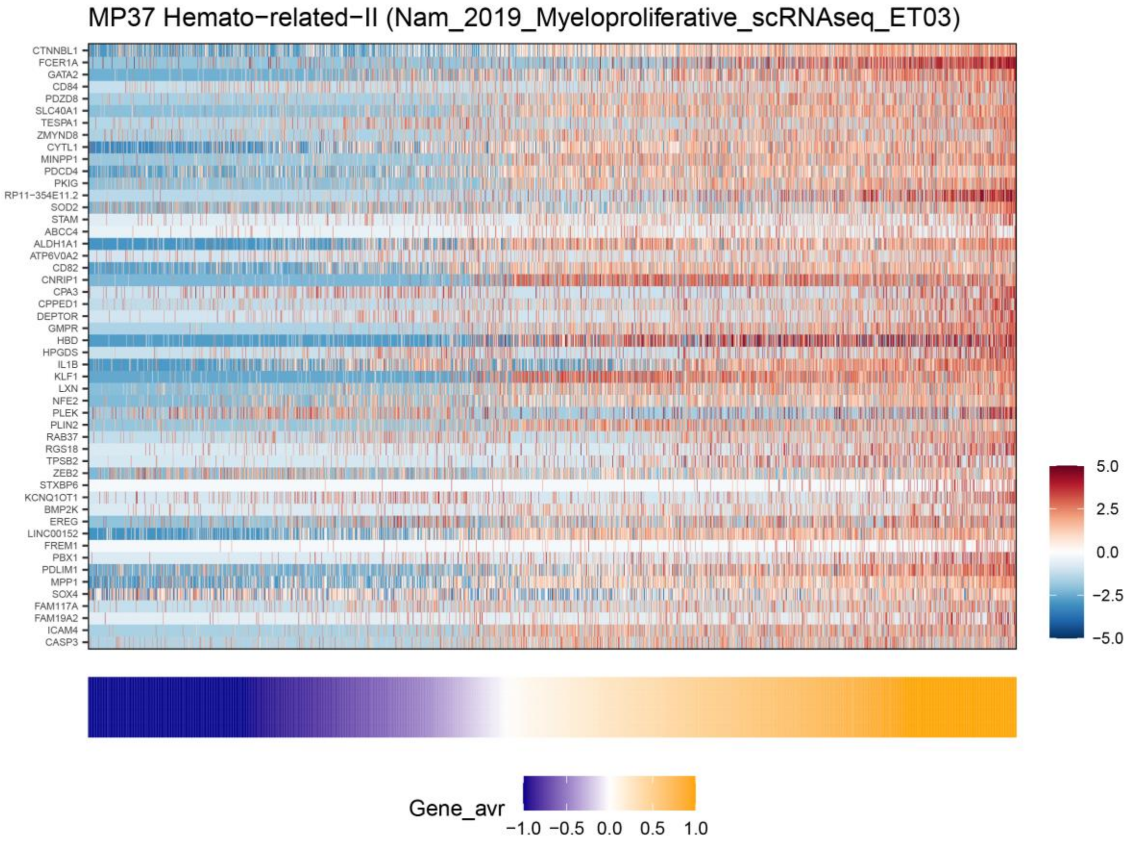

#### MP38 – Glutathione

Brief description: *Specific* MP that contains many glutathione-related genes, found only in renal cancer and not assigned to any hallmark.

Functional annotations: Enriched with annotations related to glutathione metabolism (e.g. C5.GOBP_GLUTATHIONE_METABOLIC_PROCESS).

Selected genes: GSTA1, GSTA2, GPX3, NAT8

Additional comments: Abundant only in renal cancer.

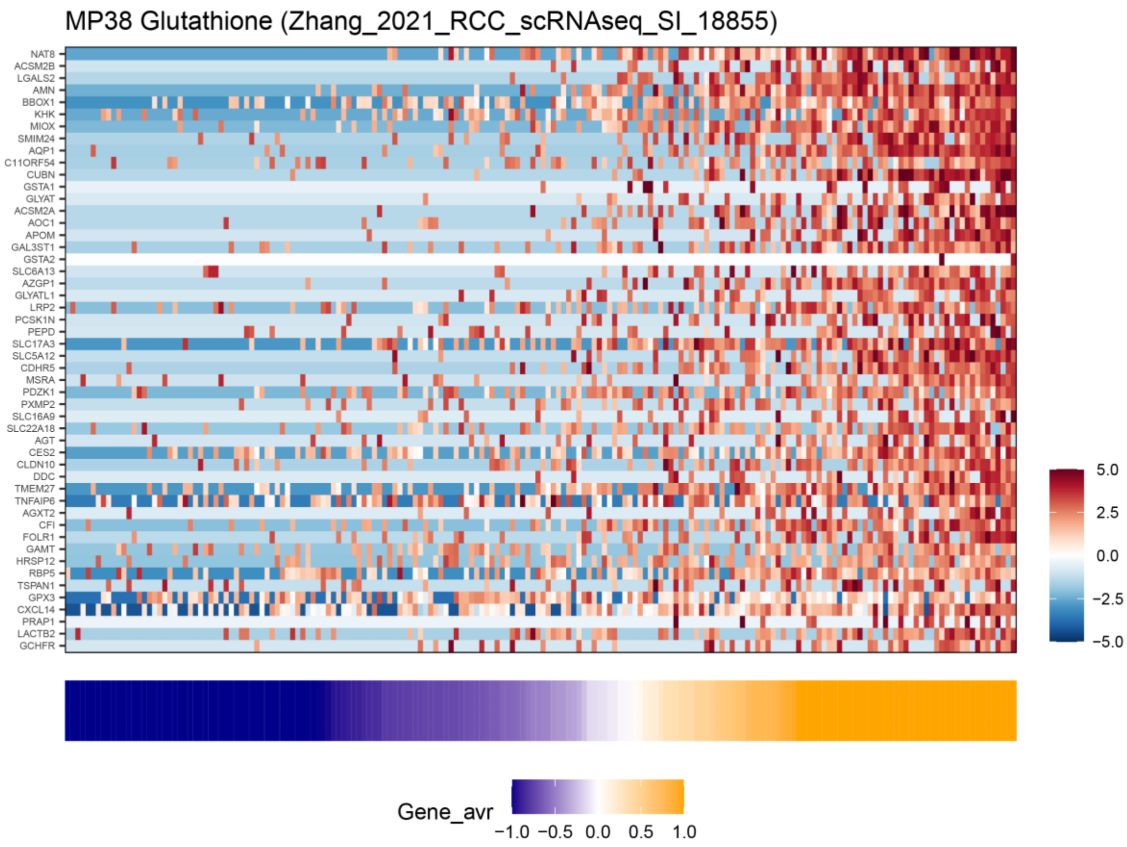

**MP39 – Metal response**

Brief description: *Specific* MP that includes many metal-response genes, and in particular methalothionine genes, found only in renal cancer and not assigned to any hallmark.

Functional annotations: Enriched with annotations related to metal ions (e.g. C5.GOBP_STRESS_RESPONSE_TO_METAL_ION)

Selected genes: MT1X, MT1E, MT1M, MT2A, MT3, MT1F, MT1A

Additional comments: Found only in renal cancer.

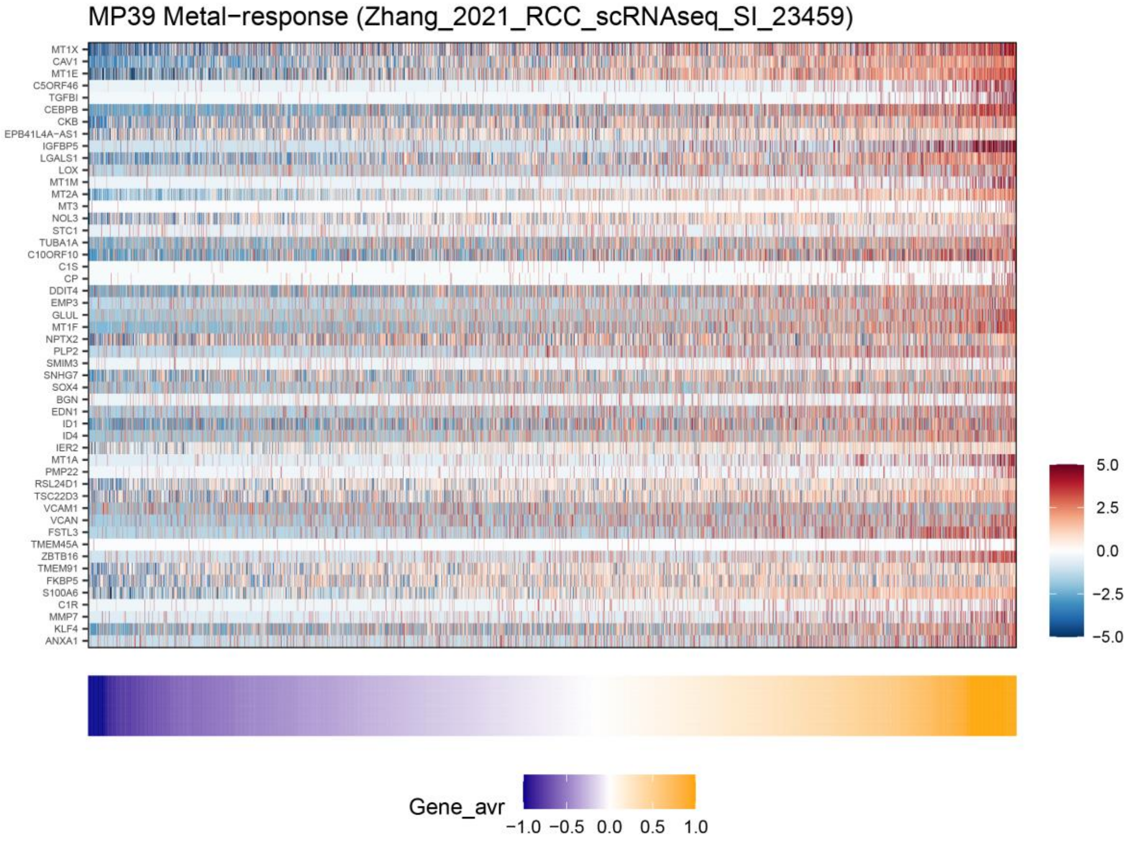

**MP40 – Unassigned I**

Brief description: *Specific* MP that we did not assign to a specific function or to a hallmark. Functional annotations with limited enrichment suggest that this might be another program that is linked to mucous and the PDAC classical program, possibly reflecting a variant of MP30.

Functional annotations: several mucous related annotations, and also one PDAC classical signature.

Additional comments: Found in PDAC and CRC.

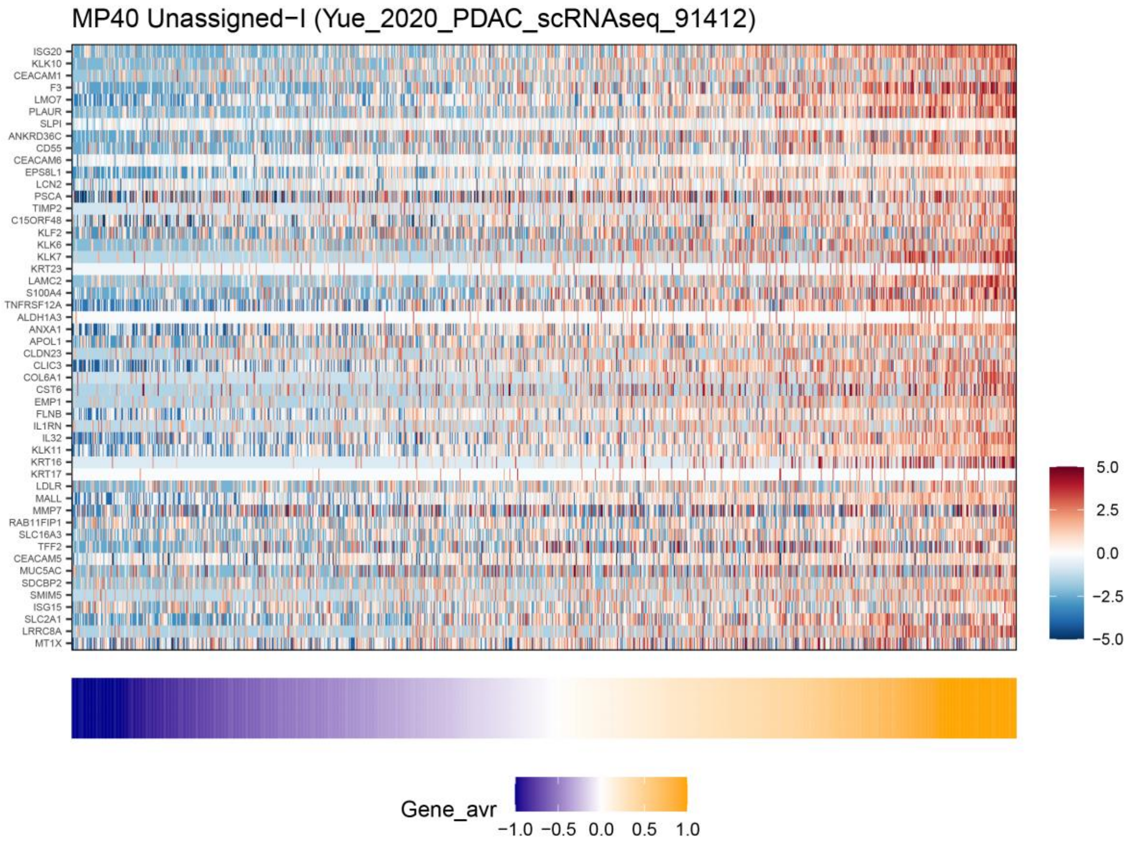

**MP41 – Unassigned II**

Brief description: *Shared* MP, not assigned to a function or a hallmark.

Functional annotations: we did not find a robust pattern in the signature list linking this MP to a certain function or cell type.

Additional comments: Most abundant in multiple myeloma followed by neuroendocrine tumors.

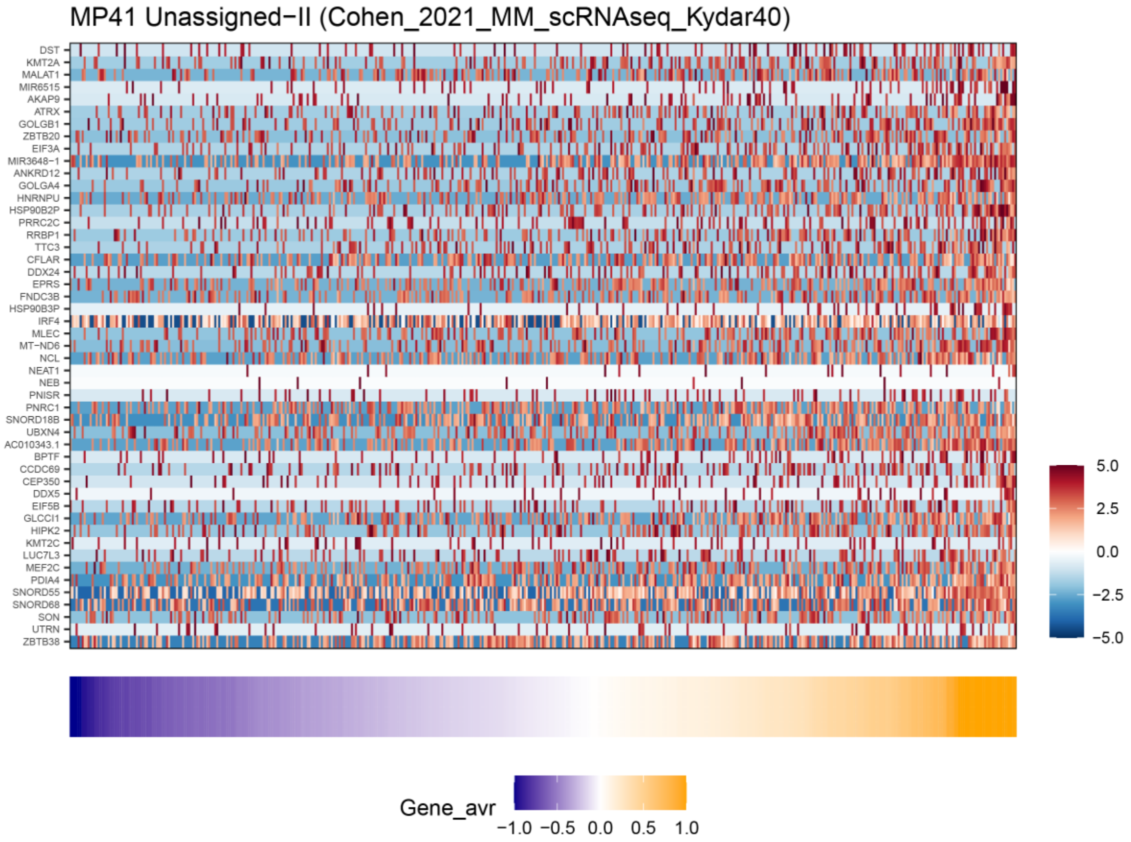
